# A Comprehensive Evaluation of Protein Structure Prediction Models for Short Peptides

**DOI:** 10.64898/2026.07.02.736085

**Authors:** Brataraj Ghosh, Arnab Mukherjee

## Abstract

Short peptides pose distinct challenges for computational structural biology due to their lack of stable tertiary structures, high conformational flexibility, and limited evolutionary signals. To address how modern deep-learning architectures navigate these challenges, we conducted a comprehensive benchmarking of five state-of-the-art protein structure prediction models: AlphaFold2, RoseTTAFold2, ESMFold, OmegaFold, and DMPfold2. Using a curated dataset of experimentally determined short peptide structures (10–49 amino acids) from the Protein Data Bank, we systematically evaluated predictive performance across varying sequence lengths and secondary structure classes. Our results demonstrate that prediction accuracy systematically improves with peptide length. Furthermore, all models perform significantly better on *α*-helical and mixed-structure peptides compared to *β*-sheet-rich and intrinsically disordered sequences. Among the evaluated methods, AlphaFold2 and the single-sequence language models, ESMFold and Omegafold proved to be the most consistent and accurate overall. We also observed that internal model confidence scores are imperfectly calibrated for short peptides, necessitating cautious interpretation. Finally, by extending our analysis to the dbAMP3 dataset of uncharacterized antimicrobial peptides, we demonstrate that a multi-model consensus approach provides a rational framework for identifying robust structural hypotheses in the absence of experimental reference structures.

## I. Introduction

In energy-landscape language, most well-folded proteins are described as minimally frustrated funnels, despite microscopic ruggedness, there is a clear global basin corresponding to the native state, separated from the unfolded ensemble by a noticeable barrier. In larger proteins, folding is often cooperative, a concerted transition stabilized by many non-local interactions (e.g. hydrophobic core packing and long-range contacts). Short peptides, by contrast, lack an extensive hydrophobic core and have relatively few side-chain contacts, so they gain little enthalpic stabilization upon folding [1], [2], [3]. They remain highly flexible because there are not enough favorable intramolecular interactions to overcome thermal fluctuations. Without multiple cooperative interactions locking the structure in place, any secondary structure a short peptide forms is only marginally stable and easily lost. A multitude of structures all lie near the minimum free energy. Kinetically, such a landscape implies barrierless or downhill folding; there are negligible energy barriers between conformations, so the peptide can rapidly hop and interconvert among many structures [4], [5]. This absence of folding cooperativity means small peptides do not exhibit the sharp two-state folding transitions seen in larger proteins, instead, their folding is gradual or continuously fluctuating due to the scarcity of strong stabilizing contacts [6].

When folding is downhill-like or effectively barrierless, as observed in some ultra-fast folding proteins and simple helix–coil peptide systems [7], [5], [8], the process becomes weakly cooperative and diffusion-like, with many shallow microstates rather than a single dominant folding barrier [9], [4], [2]. Such systems are less likely to remain trapped in one well-defined folded basin; instead, they can fluctuate rapidly between transient secondary-structure elements and random-coil-like conformations [10], [8], [11]. Given these thermodynamic and kinetic features, short peptides represent a challenging regime for both secondary-structure and tertiary-structure prediction algorithms. Classical secondary-structure predictors estimate local helix, strand, or coil propensities from sequence-derived patterns, whereas modern tertiary-structure predictors are largely optimized for the sequence-to-structure problem in which a protein sequence is assumed to adopt one dominant native conformation [12], [13], [14], [15]. This assumption is reasonable for many folded globular domains, but becomes less appropriate for short peptides, where the conformational landscape is often shallow, weakly cooperative, and strongly influenced by solvent, terminal effects, and local sequence context. In such systems, the experimentally observed structure may represent one member of a broader conformational ensemble rather than a unique native state. Therefore, disagreement between prediction models and experimental peptide structures should not be interpreted only as algorithmic failure, but also as a consequence of applying single-structure prediction frameworks to molecules whose physical behaviour is intrinsically ensemble-like. For a short and conformationally flexible peptide, the more appropriate physical picture is often one sequence corresponding to an ensemble of many low-lying structures [16], [17]. Therefore, any single static model is, at best, one representative snapshot from that ensemble. Moreover, experimentally observed structures can themselves be condition-dependent; NMR ensembles may depend on solvent, pH, ionic strength, concentration, and temperature, whereas X-ray crystal structures can be influenced by crystal-packing contacts that stabilize particular local conformational basins [10], [18]. They often predict helices or *β*-sheets based on sequence propensities, even if the real peptide is too short or too disordered to form them stably.

Unlike folded proteins, short peptides often have no strong evolutionary constraints or co-evolved contacts which AlphaFold2 exploits, so the predictions rely on generic patterns that may not apply to a floppy sequence. Other secondary-structure predictors, ranging from classical algorithms to newer AI models, similarly tend to over-predict order. They might label a peptide as helical if it has some Ala or Leu residues, or predict a *β*-hairpin in a proline-rich snippet, simply because such patterns occur in folded proteins. In reality, those same sequences, when isolated, might be disordered. Given these challenges, it is important to assess how well current deep-learning models handle peptides. McDonald et al. [19] recently undertook a large-scale benchmark of AlphaFold2 on 588 experimentally determined peptides (10-40 residues) with NMR-derived structures. They found that AlphaFold2 reliably predicted the secondary structure of many peptides, especially *α*-helices, *β*-hairpins, and disulfide-rich folds, often matching or exceeding the performance of peptide-specific methods. However, AlphaFold2 also showed systematic errors on peptides, it struggled to recover accurate backbone *ϕ/ψ* angles and disulfide bonding patterns, and its confidence metric (pLDDT) did not consistently correlate with structural accuracy. In light of these challenges, we conducted a comprehensive benchmark of five state-of-the-art protein structure prediction models, AlphaFold2 (via ColabFold), ESMFold, RoseTTAFold2, OmegaFold, and DMPfold2, assessing their ability to predict short peptide 3D structures. For this study, we compiled a dataset of 2315 experimentally determined short peptide structures from the Protein Data Bank, stratified into four length-based bins (10-19, 20-29, 30-39, and 40-49 amino acids). Predictions from each model are quantitatively evaluated against the experimental structures using confidence scores (pLDDT or equivalent), TM-score, and root-mean-square deviation (RMSD). This analysis aims to identify the strengths and weaknesses of each method across different peptide length ranges, illuminating how predictive performance varies with peptide size and highlighting areas for future improvement.

The selection of AlphaFold2 via ColabFold [20], RoseTTAFold2 [14], ESMFold [15], OmegaFold [21], and DMPfold2 [22], [23] reflects both the diversity of modern protein structure prediction architectures and their practical relevance. AlphaFold2 and RoseTTAFold2 are end-to-end neural predictors that take MSAs (and optionally templates) and directly output 3D coordinates using geometry-aware transformer architectures, without invoking external force fields or classical energy minimization. Physical realism is implicit, learned from structural databases and enforced through geometric priors such as SE(3)-equivariance [24], invariant pair representations, and torsion-based coordinate generation. AlphaFold2, as implemented in ColabFold, represents the current gold standard, employing a two-track transformer with iterative recycling and deep evolutionary information, while RoseTTAFold2’s three-track architecture jointly integrates sequence, pairwise, and coordinate representations, offering a complementary and competitive paradigm [25], [14].

In contrast, ESMFold eliminates MSAs entirely, using a large protein language model (ESM-2) [26] trained on sequence data alone to infer structure via a lightweight folding trunk [27]. It relies solely on learned statistical priors, without explicit physics or refinement. OmegaFold similarly operates without MSAs, combining language model embeddings with SE(3)- equivariant transformers [24] and iterative, diffusion-inspired coordinate refinement [21]. Both methods are computationally efficient and suited for low-homology or short sequences, but are strongly shaped by sequence-derived statistical biases.

DMPfold2 [22], [23] represents a distinct distance-map-based prediction strategy compared with the other evaluated models. It uses MSA-derived features to predict inter-residue distance and precision information, which is then used to reconstruct peptide coordinates through a dedicated coordinate-generation module. This separates DMPfold2 from transformer-based or protein-language-model-based predictors and makes it a useful contrasting model in the benchmark.

Together, these models span complementary prediction paradigms, from single-sequence protein-language-model-based predictors such as ESMFold and OmegaFold, to MSA-dependent geometric deep-learning frameworks such as AlphaFold2 and RoseTTAFold2, to distance-map-guided coordinate reconstruction as implemented in DMPfold2. This diversity is critical for evaluating performance on short peptides, where weak cooperativity, shallow free-energy landscapes, and limited evolutionary signal challenge assumptions learned from globular proteins.

The rest of the paper is organized as follows: Section II outlines the architectural backgrounds of the five deep-learning protein structure prediction models evaluated in this study. Section III details the methodology, including dataset construction, stratification strategies, the structure prediction workflow, and the evaluation metrics employed. Section IV presents the characteristics of the experimental benchmark dataset and introduces the dbAMP3 dataset used for our consensus-based correlation studies. Section V reports the results of our comparative analysis, assessing model performance across varying sequence lengths and secondary structural classes using various metrics. Section VI discusses the broader implications of these findings, highlighting the limitations of current architectures when applied to short, flexible peptides, and evaluating the practical utility of multi-model consensus frameworks. Finally, Section VII summarizes our conclusions.

## II. Background on Models

The selected models for this study span a spectrum of approaches; from single-sequence protein-language-model-based predictors (ESMFold, OmegaFold), to MSA-dependent geometric deep-learning frameworks (AlphaFold2, RoseTTAFold2), to distance-map-guided coordinate reconstruction (DMPfold2). A key distinction among these approaches lies in how structural geometry is represented and enforced. Modern end-to-end predictors do not generally rely on external force fields or classical structure-refinement engines during inference; instead, they learn statistical representations of protein geometry from structural databases through pair representations, attention-based geometric modules, torsion-based coordinate generation, and related structural constraints. In contrast, DMPfold2 separates MSA-derived distance prediction from coordinate reconstruction, making it a useful constraint-driven comparator to the transformer and language-model-based predictors. Understanding this methodological diversity is particularly important when evaluating performance on short peptides, where weak cooperativity, shallow free-energy landscapes, and limited evolutionary signal challenge assumptions learned from globular proteins. Figure 3 depict the architectures of the five models we consider.

**Fig. 1:**
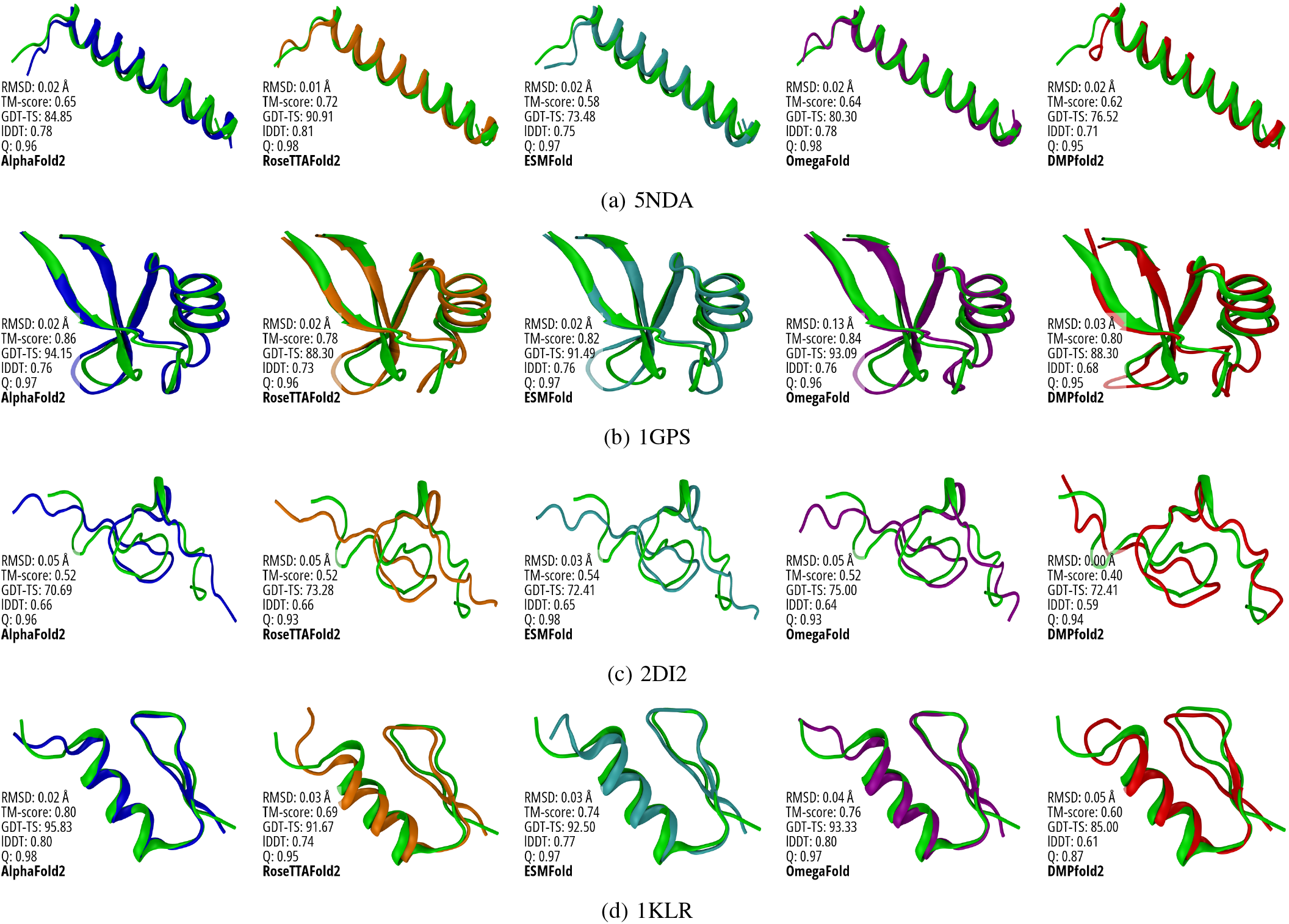
Representative example of high structural agreement between predicted peptide models and the experimentally determined PDB reference. The PDB structure is shown in green, while model predictions are coloured according to model identity: AlphaFold2 in blue, RoseTTAFold2 in orange, ESMFold in cyan, OmegaFold in purple, and DMPfold2 in red. Each predicted structure was aligned to the PDB reference before comparison, and the corresponding structural similarity metrics are reported in each panel, including RMSD, TM-score, GDT-TS, LDDT, and native-contact fraction *Q*. The overlays illustrate both the visual agreement between predicted and experimental conformations and the metric-based differences in local geometry, global topology, and contact recovery across prediction models.

**Fig. 2:**
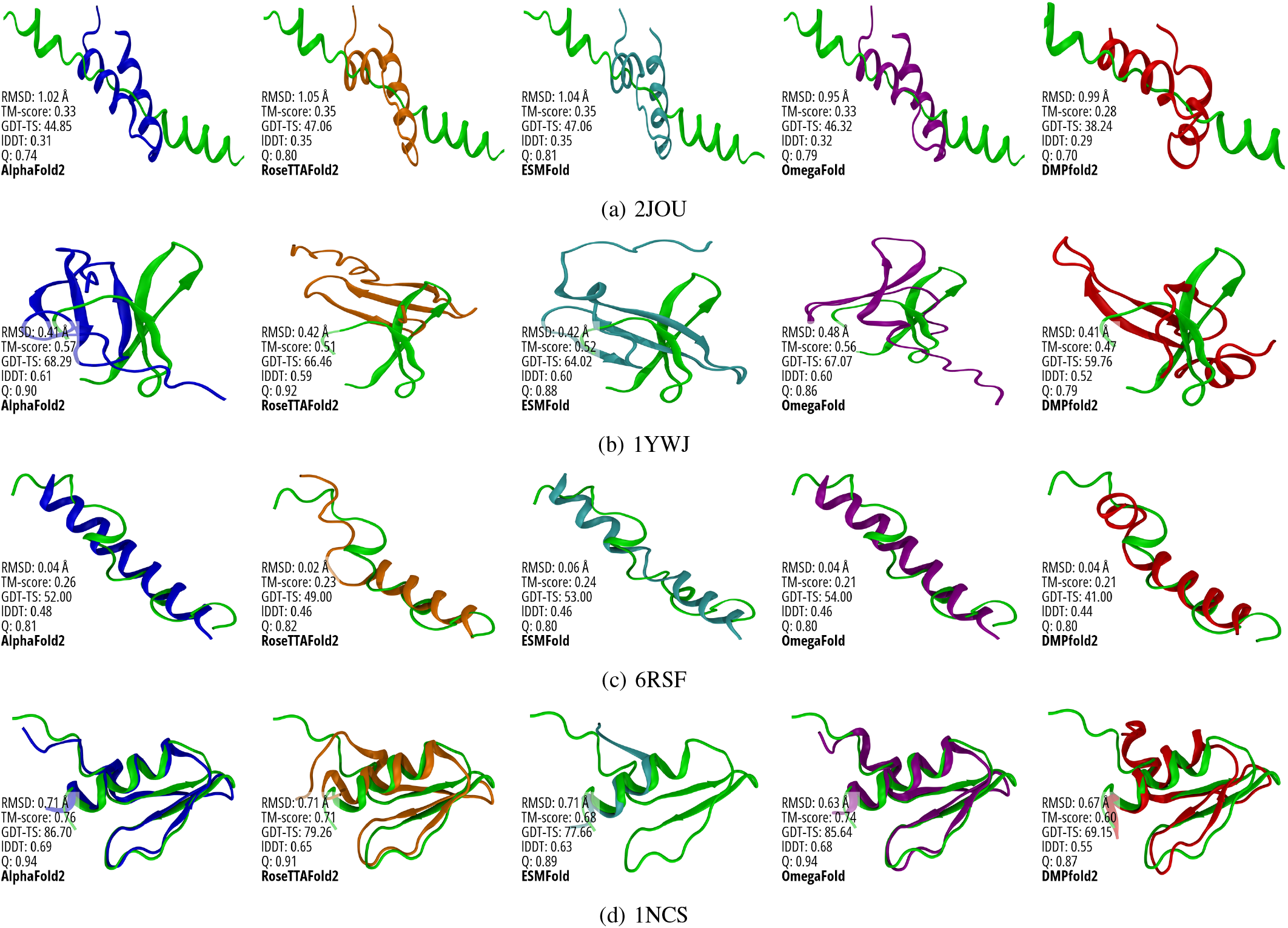
Representative example of low structural agreement between predicted peptide models and the experimentally determined PDB reference. The same colour scheme is used as in the high-agreement example: PDB reference in green, AlphaFold2 in blue, RoseTTAFold2 in orange, ESMFold in cyan, OmegaFold in purple, and DMPfold2 in red. Compared with the high-agreement case, the predicted structures show larger deviations from the PDB conformation, reflected by higher RMSD and reduced TM-score, GDT-TS, LDDT, and native-contact fraction *Q* values.

**Fig. 3:**
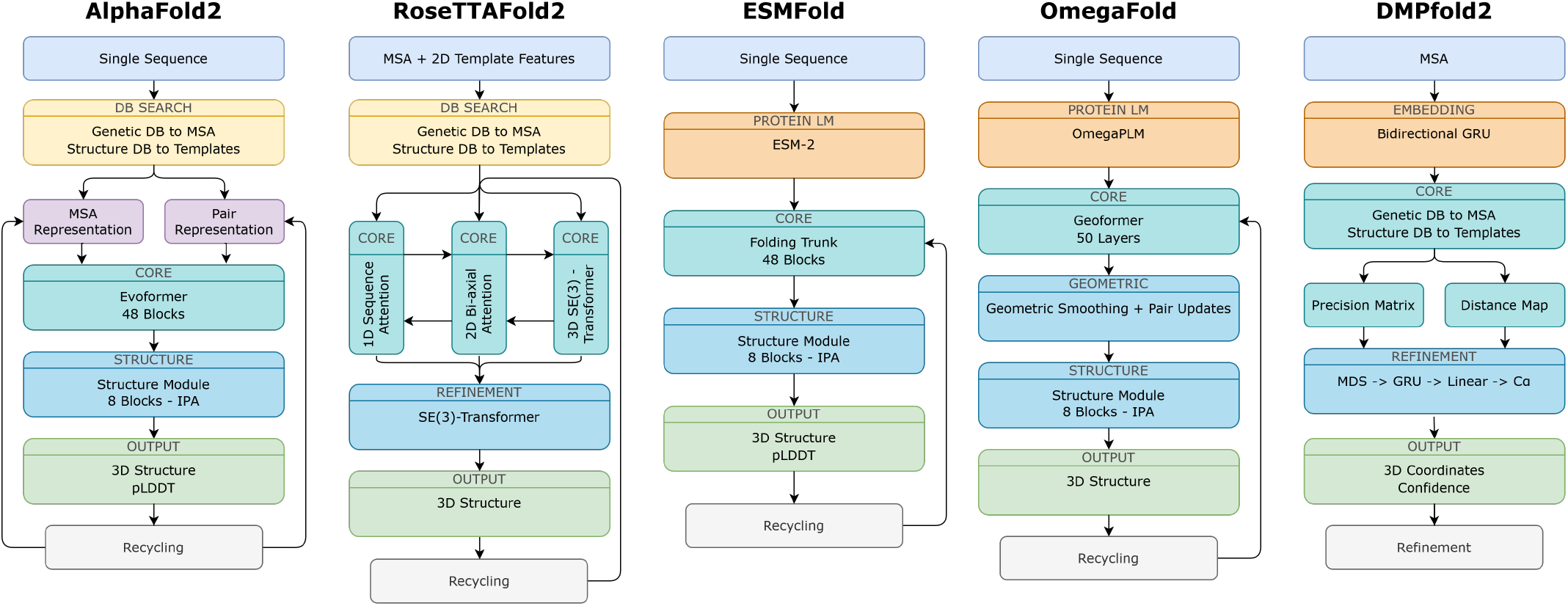
Model Architectures of AlphaFold2, RoseTTAFold2, ESMFold, OmegaFold, and DMPfold2

### a) AlphaFold2

AlphaFold2 [25], [20] represents a fundamental shift in protein structure prediction by replacing explicit physical modeling with an end-to-end learned mapping from sequence and evolutionary information to three-dimensional coordinates. Rather than sampling conformational space with a force field, AlphaFold2 encodes protein sequences and their multiple sequence alignments (MSAs) into rich pairwise and residue-level representations using a deep transformer architecture, iteratively refining these representations through a process known as recycling [25]. Central to the model is the Evoformer module [25], which propagates information between sequence positions and residue pairs, effectively learning statistical regularities associated with native protein folds, including long-range contacts and packing constraints, directly from structural databases. The subsequent structure module converts these abstract representations into atomic coordinates using torsion-based updates and SE(3)-equivariant [24] operations, ensuring rotational and translational consistency while implicitly enforcing realistic covalent geometry. Importantly, AlphaFold2 does not optimize an explicit energy function; instead, its notion of physical plausibility is encoded in learned geometric priors and the statistics of the training data. As implemented in the ColabFold pipeline [20], it represents the current gold standard in structure prediction. However, its strong inductive bias toward single, well-defined native states, reflecting the predominance of globular proteins in the Protein Data Bank, necessitates caution when interpreting predictions for short peptides or intrinsically disordered systems, where a unique deep free-energy minimum may not exist.

### b) RoseTTAFold2

RoseTTAFold2 [14] extends the Rosetta tradition of structure prediction by integrating deep learning directly into the coordinate-generation process, rather than treating it as a post hoc scoring or refinement step. Its defining feature is a three-track neural network architecture that simultaneously processes sequence information, pairwise residue relationships, and three-dimensional coordinates, allowing information to flow bidirectionally between these representations during inference [14]. By updating distances, orientations, and Cartesian coordinates in parallel, RoseTTAFold2 learns to couple local sequence features with emerging global structure, effectively capturing long-range interactions without relying solely on explicit sampling. This architecture provides a complementary paradigm to AlphaFold2 and achieves competitive accuracy, particularly for moderately sized proteins. While less dependent on deep MSAs, its performance still improves with richer evolutionary information. As with other end-to-end predictors, physical realism is imposed implicitly through learned geometric constraints rather than explicit energy evaluation. Consequently, the model tends to favor compact, ordered conformations, and its predictions for short peptides or weakly stable systems should be interpreted as statistically plausible structures rather than definitive native states.

### c) ESMFold

ESMFold [15] represents a departure from MSA-driven structure prediction by relying exclusively on single-sequence information encoded through large protein language models. Built on Meta’s ESM-2 [26] architecture, ESMFold is trained via masked-language modeling on hundreds of millions of protein sequences, allowing it to internalize statistical regularities of amino-acid usage, local motifs, and long-range dependencies without explicit evolutionary alignments. Structural prediction is achieved by passing the learned sequence embeddings through a comparatively lightweight folding trunk, which converts attention-derived representations into backbone frames and atomic coordinates. Unlike AlphaFold2 and RoseTTAFold2, ESMFold does not employ recycling of predicted structures across multiple forward passes, although its folding trunk performs iterative representation updates prior to coordinate prediction. It also does not invoke any explicit geometric or physics-based refinement. Instead, it relies entirely on learned priors from large-scale unsupervised training to generate conformations that are statistically close to physical minima. This makes it computationally efficient and particularly suitable for orphan sequences or peptides lacking evolutionary depth. However, its reliance on generic sequence-structure correlations can lead to overprediction of secondary structure in systems with inherently flat or degenerate free-energy landscapes.

### d) OmegaFold

OmegaFold [21] is a single-sequence protein structure predictor that further expands the landscape of MSA-free methods. It combines embeddings from a large pre-trained protein language model with a structure module based on SE(3)-equivariant transformers [24], enabling direct reasoning over three-dimensional geometry while preserving rotational and translational invariance [28]. The model iteratively refines Cartesian coordinates through equivariant message passing, in a process conceptually reminiscent of diffusion-inspired refinement, as represented as the refinement loop in Figure 3. Although this procedure improves geometric consistency, it does not correspond to minimizing an explicit energy function; instead, structural plausibility emerges from learned statistical patterns and invariant operations. Like ESMFold, OmegaFold is particularly attractive for short sequences and low-homology regimes, but its predictions are strongly influenced by biases learned from globular proteins, often favoring compact conformations even when the true landscape is highly flexible or weakly cooperative.

OmegaFold uses the concept of pseudo-MSA, which tries to imitate the information role of an MSA without actually collecting homologous sequences. Instead of real aligned sequences, the model generates or learns an internal representation that behaves like an evolutionary or contextual feature. A multiple sequence alignment (MSA) is normally generated by searching sequence databases for homologous proteins and aligning them to the query sequence. This provides evolutionary information, because residue pairs that mutate together across homologs often indicate spatial proximity in the folded structure. A pseudo-MSA, in contrast, is an MSA-like representation that is not obtained from real homologous sequence alignment. Instead, it refers to artificial or internally learned sequence features that attempt to provide similar contextual or evolutionary information from a single input sequence.

### e) DMPfold2

DMPfold2 [22] occupies a distinct position by combining machine learning-based restraint prediction with explicit physics-inspired structure generation. A deep convolutional neural network predicts inter-residue distance distributions, hydrogen-bonding patterns, and backbone *ϕ/ψ* torsions from sequence-derived features [22]. DMPfold2 starts from an input multiple sequence alignment (MSA), from which sequence-derived and coevolutionary features are extracted. These features are embedded through recurrent neural-network components, including bidirectional GRU layers, to generate residue-pair representations. The core prediction stage then infers inter-residue structural constraints, including distance maps and contact-like restraint information, which define the expected spatial relationships between residues.

The predicted distance information is then used as internal geometric constraints that guide the model’s own neural coordinate-generation module. In this stage, multidimensional scaling (MDS) provides an initial coordinate representation, which is further processed through GRU and linear layers to generate C-*α* coordinates. Main-chain and C-*β* atoms are then added to obtain full backbone-level structural models, together with a confidence estimate. Because of this, DMPfold2 differs from end-to-end predictors such as AlphaFold2, ESMFold, OmegaFold, and RoseTTAFold2 by explicitly separating MSA embedding, distance-map prediction, and coordinate reconstruction. However, the final structures remain dependent on the accuracy of the predicted restraints, which are themselves learned from datasets dominated by folded proteins. For short peptides, this can result in physically consistent yet overly ordered conformations relative to the underlying free-energy landscape.

## III. Methodology

In this section, we detail the methodology and steps involved in collecting the dataset, dataset pre-processing, simulation setup, and execution and evaluation of the predictions. The complete pipeline consists of seven steps: (1) Dataset collection from PDB [29], (2) Sequence extraction, (3) Dataset filtering and preprocessing, (4) Structure prediction using five models, (5) Structural alignment with reference structures, (6) Metric computation, and (7) Statistical comparison.

### A. Dataset Construction and Pre-processing

The study focuses specifically on short peptides, defined here as sequences between 10 and 50 amino acids in length. This length range was selected because peptide structure prediction presents unique challenges compared to globular proteins, including increased conformational flexibility and a reduced number of long-range contacts.

Experimentally determined peptide structures obtained from the Protein Data Bank were used as reference structures for benchmarking. These structures provide experimentally resolved conformations against which predicted models can be evaluated using coordinate-based, topology-based, and contact-based structural similarity metrics. The following filtering criteria were applied during dataset construction: (1) Sequence length between 10 and 50 amino acids. (2) Structures corresponding to single peptide chains. (3) Removal of multichain complexes. (4) Removal of non-standard amino acid residues. (5) Exclusion of entries with incomplete sequence annotations. Following filtering, the resulting dataset consisted exclusively of single-chain peptide structures composed of standard amino acids. The corresponding amino acid sequences were extracted from the PDB records and stored in FASTA format, which served as the input for structure prediction. The experimentally determined structures from the PDB were used as reference (native) structures for structural comparison and accuracy evaluation.

### B. Dataset Stratification

To enable a detailed analysis of prediction performance across different peptide characteristics, the dataset was stratified using two independent bucketing strategies: sequence length and secondary structure composition. Peptides were categorized into four groups based on sequence length: 10-20, 20-30, 30-40, 40-50. This stratification allows the analysis of how prediction accuracy varies with peptide length, which is an important factor influencing structural complexity and conformational variability.

In addition to length-based grouping, peptides were also categorized according to their dominant secondary structure composition. Secondary structure assignments were derived from the experimentally determined native structures using the DSSP (Dictionary of Secondary Structure of Proteins) secondary structure annotation method [30]. Based on these assignments, peptides were grouped into four structural categories, i.e., *α*-helix-rich peptides, in which *α*-helical elements constitute the dominant structural motif. *β*-sheet-rich peptides, characterized by a predominance of *β*-strand or *β*-sheet conformations. Mixed-structure peptides, containing substantial contributions from both *α*-helical and *β*-sheet elements without a single dominant structural motif. Disordered peptides, in which the structure is primarily composed of coil or irregular conformations with minimal stable secondary structure.

This classification allows the evaluation of prediction performance across peptides with different structural organizations and varying degrees of conformational order. Together, the length-based and secondary-structure-based stratification schemes enable systematic benchmarking of prediction methods across peptides with varying sequence lengths, structural motifs, and conformational complexity, providing a more nuanced assessment of model performance under diverse structural regimes.

### C. Structure Prediction Workflow

For each peptide sequence in the dataset, structures were predicted independently using all five models. The primary input to all prediction pipelines was the FASTA sequence corresponding to each peptide. No template structures were explicitly provided, ensuring that all models performed de novo structure prediction. Structure prediction for models requiring multiple sequence alignments (MSAs), such as AlphaFold2 and RoseTTAFold2, was performed using the ColabFold inference pipeline [20]. ColabFold provides optimized scripts for running AlphaFold-based structure prediction workflows, including automated generation of MSAs and streamlined model inference. For each peptide sequence: (1) The FASTA sequence was provided as input. (2) The ColabFold pipeline generated MSAs using standard sequence databases. (3) The respective prediction models were executed to generate predicted three-dimensional structures.

Models capable of single-sequence inference, such as ESM-Fold and OmegaFold, were run directly using the peptide FASTA sequence without requiring explicit MSA generation. All predicted structures were generated in PDB format for downstream structural comparison and metric computation. One thing to note: for AlphaFold2, only the rank-1 predictions are considered for further analysis.

### D. Computational Environment & and Inference Parameters

All structure prediction pipelines were executed within a standardized computational environment to ensure consistent benchmarking. Model inference was performed utilizing a server with a Nvidia A100 80GB GPU and 256 GB of memory, running on Ubuntu 20.04 with CUDA 11.3. However, due to compatibility issues of the CUDA environment, the DMPfold2 predictions were performed on a different server with Nvidia GTX 1080Ti 12 GB, running CUDA 10.2.

To ensure a robust evaluation across the diverse set of architectural paradigms, model-specific inference parameters were carefully configured, as detailed in Table I. MSA-Dependent Models, such as AlphaFold2 and RoseTTAFold2, was streamlined using the ColabFold inference pipeline. To allow for extensive structural refinement, both AlphaFold2 and RoseTTAFold2 were configured to perform 48 recycle iterations. For AlphaFold2, the maximum MSA and pseudo-MSA depths were set to 512:1024, and the maximum relax iterations (amber [31]) were capped at 200. It is important to note that for AlphaFold2, only the rank-1 predictions were retained for downstream structural analysis and metric computation. RoseTTAFold2 was executed using the RF2_jan24 model weights with a maximum MSA depth of 512, and the use of masked-language modeling (MLM) was disabled. For the without MSA simulations, we specify the MSA mode / method to single sequence, instead of mmseqs2 [32] for AlphaFold2 and RoseTTAFold2, while for DMPfold2, we had to isolate the single sequence at the first generation stage.

**TABLE I:**
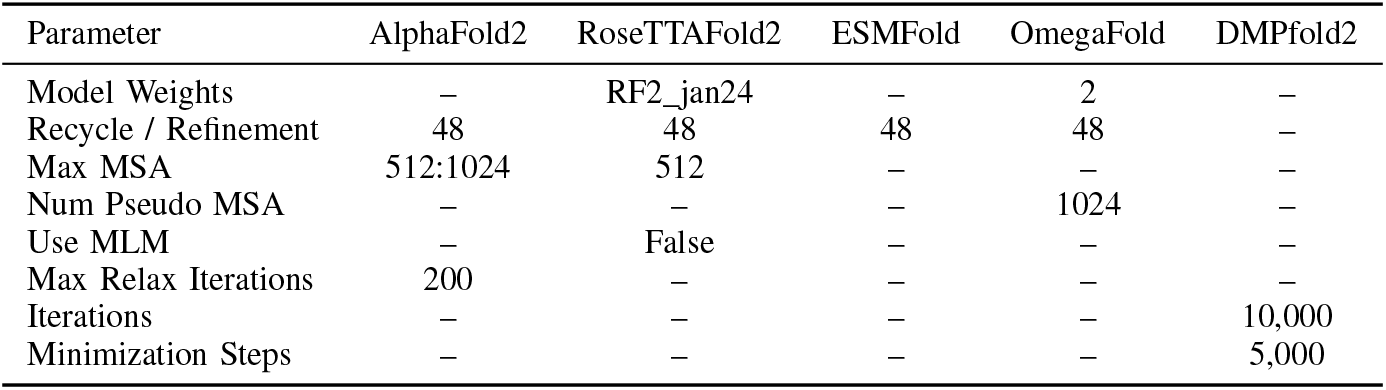
Configuration parameters for different protein structure prediction models.

| Parameter | AlphaFold2 | RoseTTAFold2 | ESMFold | OmegaFold | DMPfold2 |
| --- | --- | --- | --- | --- | --- |
| Model Weights | — | RF2_jan24 | — | 2 | — |
| Recycle / Refinement | 48 | 48 | 48 | 48 | — |
| Max MSA | 512:1024 | 512 | — | — | — |
| Num Pseudo MSA | — | — | — | 1024 | — |
| Use MLM | — | False | — | — | — |
| Max Relax Iterations | 200 | — | — | — | — |
| Iterations | — | — | — | — | 10,000 |
| Minimization Steps | — | — | — | — | 5,000 |

**TABLE II:** Dataset Metadata.

| Property | Details |
| --- | --- |
| Source | Protein Data Bank (PDB) |
| Extraction Date | 2025-06-18 |
| Sequence Length Range | 10-49 Residues |
| Total Peptides | 2315 |
| Unique Peptides | 1964 |

Models capable of single-sequence inference, i.e., ESMFold and OmegaFold were executed directly using the peptide FASTA sequences, circumventing the need for explicit MSA generation. Consistent with the MSA-dependent models, both ESMFold and OmegaFold were set to 48 iterations of refinement cycles. Additionally, OmegaFold was run utilizing model weights designated as “2” and configured with a maximum pseudo-MSA parameter of 1024. For the restraint-guided model DMPfold2, to accommodate its distinct distance geometry and molecular mechanics refinement phase, DMPfold2 was configured to perform 10,000 maximum relax iterations alongside 5,000 minimization steps.

## IV. The Benchmark Dataset

The benchmark dataset used in this study was constructed to provide a representative and systematically curated collection of experimentally determined short peptide structures. All structures were extracted from the Protein Data Bank (PDB), with the dataset finalized on June 18, 2025, as depicted in Table Peptides were selected based on sequence length, retaining entries between 10 and 49 amino acids, spanning the boundary between very short peptides and small, marginally structured protein domains. After filtering, the final dataset comprises 2315 peptide structures corresponding to 1964 unique sequences.

The sequence-length distribution of the dataset is shown in Figure 45. Lengths are broadly distributed across the selected range, with no single length dominating the dataset, although a mild enrichment is observed in the 25-40 residue interval. This range captures peptides long enough to transiently form secondary-structure elements, yet typically too short to support a well-packed hydrophobic core or stable tertiary fold. Shorter peptides (10-19 residues) are also well represented, ensuring coverage of highly flexible, weakly cooperative systems where the free-energy landscape is expected to be particularly shallow. The inclusion of peptides up to 49 residues allows assessment of how predictive performance evolves as chain length increases and cooperative interactions become incrementally more accessible.

Figure 46 summarizes the amino acid composition of the dataset, reported as relative frequencies across all sequences. The distribution broadly reflects typical peptide and protein composition, with no extreme biases toward a narrow subset of residues, indicating that the dataset is not artificially enriched for specific motifs or chemistries. Hydrophobic residues such as leucine, isoleucine, and valine are present at moderate frequencies, consistent with their general prevalence in protein sequences, while charged and polar residues remain well represented, supporting a diverse range of electrostatic and solvation environments. Notably, glycine and proline residues known to disrupt regular secondary structure occur at appreciable levels, reinforcing the intrinsically flexible character of many peptides in the dataset and increasing the challenge for structure prediction methods that implicitly favor ordered conformations.

Together, these characteristics make the dataset well-suited for benchmarking structure prediction models in a regime that departs from the canonical “one sequence, one folded structure” assumption. The combination of broad length coverage, chemically diverse sequences, and experimentally determined structures, many of which reflect conformational heterogeneity rather than a single rigid fold, provides a stringent and realistic testbed for evaluating how modern deep-learning-based predictors perform on short peptides. In particular, the dataset enables systematic analysis of how prediction accuracy, confidence metrics, and structural biases vary as a function of peptide length and sequence composition, thereby illuminating the strengths and limitations of current models when applied outside their primary training domain.

Additionally, beyond the experimental benchmark dataset, the dbAMP3 dataset [33] was utilized to perform broader correlation studies across a large set of uncharacterized antimicrobial peptides. These studies allowed us to investigate whether the general trends and behavioral relationships observed in our reference-based analysis persist within a larger, biologically relevant population. Consequently, this correlation analysis provides critical context for understanding how consistently these deep-learning predictors operate when applied to realworld sequence databases.

## V. Results

### A. Structural Deviations & RMSD Analysis

Root-mean-square deviation (RMSD) between predicted and experimental C-*α* coordinates is a widely used metric for structural accuracy. RMSD was calculated only over C-*α* atoms belonging to residues assigned to structured regions, following the convention used in previous peptide-structure benchmarking work [19]. C-*α* atoms from structureless or unassigned regions were excluded from the RMSD calculation. Therefore, the RMSD values primarily report the agreement of the ordered peptide core rather than the deviation of the entire peptide chain. This convention reduces the influence of flexible termini and poorly defined structureless segments, which can otherwise dominate RMSD despite having limited relevance to the folded structural region. Low RMSD indicates tight atomic agreement, while large RMSD shows gross mismatch in backbone placement. Across all five predictors, a clear length-dependent improvement in structural accuracy is observed. Both the mean and median RMSD values decrease as peptide length increases from the shortest bin to ∼ 30-49 residues. For example, in AlphaFold2 the mean RMSD decreases from 0.30 ± 0.54 Å in the 10-19 residue bin to ∼0.12 ± 0.16 Å for peptides between 30-49 residues, as depicted in Table III. Similar trends are observed for RoseTTAFold2 and ESMFold, where mean RMSD values shift from 0.43±0.54 Å to ∼ 0.15 Å and 0.36 ± 0.60 Å to ∼ 0.13 Å, respectively. This matches expectations, as longer peptides can form more intra-chain contacts and stabilising interactions, producing deeper local minima that predictors can recover. As it can be observed in Figure 4, AlphaFold2 produces moderate RMSD in the 10-19 bin (many examples in the *>* 0.25 Å regime) with a small bump around 0.5 Å, but RMSD distributions shift left and narrow for lengths scale from 20-39 residues and above; in the 30-49 bin a substantial fraction of peptides fall into the 0.1 Å range, with a tail of near-native predictions. While RoseTTAFold2 closely mirrors AlphaFold2, it is slightly less accurate on average within the shortest sequence bins. This discrepancy is particularly notable in the 10-19 amino acid (aa) range, where the median deviation increases from 0.09 Å in AlphaFold2 to 0.25 Å in RoseTTAFold2. A similar trend is observed in the 20-29 aa range, with the mean shifting from 0.08 Å in case of AlphaFold2 to 0.15 Å in RoseTTAFold2. Analysis of length-wise RMSD distributions reveals that ESMFold outperforms RoseTTAFold2 in the shortest sequence bins (10-19 aa and 20-29 aa). Compared to AlphaFold2, for RMSD distribution, the mean and median values of the ESMFold predicted structures are very close to RoseTTAFold2. But the tail of the distribution is noticeably broader, characterized by a thicker tail of high-RMSD outliers. In contrast, AlphaFold2 maintains a tighter distribution with fewer catastrophic failures. As sequence length increases (30-39 aa and 40-49 aa), distribution shifts toward lower RMSD values. This suggests that ESMFold’s language-model priors require more residue context to resolve structures reliably; while its single-sequence embeddings can effectively recover stable motifs, they are more prone to error in short, context-poor sequences than AlphaFold2.

**Fig. 4:**
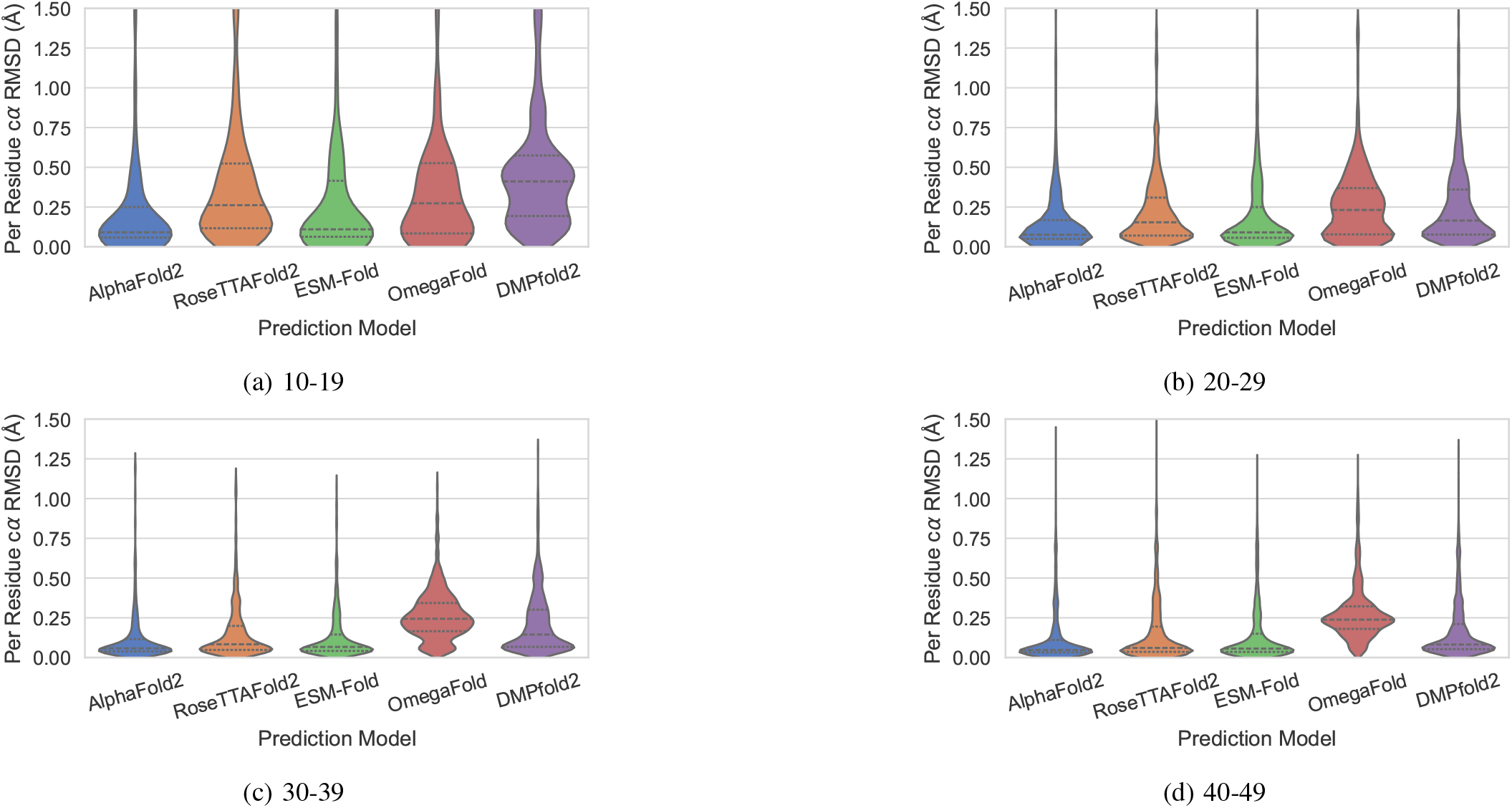
Length-wise comparison of C(*α*) RMSD distributions across prediction models. Violin plots show the distribution of model-to-reference RMSD (Å) values within each peptide length bin: 10-19, 20-29, 30-39, and 40-49 residues. Lower RMSD values indicate closer agreement with the experimental reference structure, whereas broader distributions and extended upper tails indicate larger model-dependent deviations.

Across all length bins, DMPfold2 shows the weakest RMSD match of any method, and this is especially evident in the tails of its distributions. Even in the shortest 10-19 aa bin, where one might expect the problem to be easiest, the DMPfold2 RMSD histogram is very broad and the right-hand tail extends well beyond 1.0 Å, reaching up to ∼ 1.75 Å for some peptides. This heavy tail persists in the longer bins: for 20-29, 30-39, and 40-50 aa, the bulk of the distribution remains centered at relatively high RMSD, and in every case there is a substantial fraction of structures with RMSD *>* 1.0 Å, indicating frequent large-scale misplacements rather than just small local deviations. In contrast, the end-to-end neural models (AlphaFold2, ESMFold, RoseTTAFold2, and especially OmegaFold for 30-49 aa) show much tighter, left-shifted distributions, where RMSD values above 1 Å are rare outliers rather than a routine outcome. OmegaFold shows intermediate performance that is noticeably better than DMPfold2 but consistently more variable than AlphaFold2 and ESMFold. In the 10-19 aa bin, OmegaFold produces moderately compact RMSD distributions but already exhibits a clear right-hand tail reaching 0.8 to 1.0 Å, indicating that even very short peptides occasionally deviate substantially from the experimental reference. As peptide length increases to 20-29 aa and especially 30-39 aa, the RMSD distributions broaden further, often becoming multi-modal with distinct secondary peaks around 0.6-0.8 Å which is also can be observed in case of DMPfold2, suggesting heterogeneous prediction behaviours across peptides with different secondary structure types. In the 40-49 aa bin, OmegaFold’s variability becomes pronounced; the mean shifts upward and the high-RMSD tail extends to ∼1.2-1.4 Å (Figure 4), substantially higher than AlphaFold2 and ESMFold in the same length category.

Across structural classes, (*α*)-helix-rich and mixed-content peptides consistently yield the lowest RMSD values for all five models as indicated in Figure 5, indicating that these two categories are the easiest for current predictors. In the *α*-helix class, the RMSD distributions for AlphaFold2 and ESMFold are extremely tight (means ∼ 0.12-0.16 Å), with narrow variance and almost no long-RMSD tail, reflecting the fact that helices have strong local hydrogen-bonding patterns and predictable *ϕ/ψ* correlations. RoseTTAFold2 performs slightly worse (mean ∼ 0.18 Å), followed by OmegaFold (mean ∼ 0.17-0.22 Å), which shows modest but clear widening of the tail. DMPfold2 is the least accurate even for helices (means ∼ 0.22 Å), showing extended tails up to 0.7 Å, demonstrating that its restraint-driven folding approach struggles to reproduce subtle geometric regularities in short peptides. A similar pattern occurs in mixed-structure peptides (helix-turn-coil or helix-hairpin combinations), where AlphaFold2 and ESMFold again show tight distributions (means 0.11-0.14 Å), RoseTTAFold2 is moderately broader, OmegaFold exhibits noticeable variability, and DMPfold2 shows the largest spread. Mixed peptides benefit from partial local structure while not relying heavily on long-range contacts, making them more learnable for most models. In contrast, *β*-sheet-rich and disordered peptides exhibit substantially higher mean RMSDs and broader, heavier-tailed distributions, indicating that these categories remain distinctly challenging. *β*-sheet peptides require correct strand pairing, registry, and long-range hydrogen-bond geometry features that are hard to infer in short chains with limited coevolutionary signal. As a result, mean RMSDs rise to ∼ 0.19-0.22 Å for AlphaFold2/ESMFold but increase dramatically for RoseTTAFold2 (means ∼ 0.28 Å) and especially OmegaFold, which shows large variance and tails often exceeding 1.0-1.5 Å. DMPfold2 performs by far the worst on *β*-sheets (means ∼ 0.38 Å and tails to ∼ 1.4 Å), reflecting the difficulty of reconstructing sheet topology from noisy restraints. The disordered category shows the greatest overall dispersion: even AlphaFold2 and ESMFold generate means around 0.21-0.23 Å, but with clear tailing up to ∼ 1.0 Å due to the mismatch between a dynamic experimental ensemble and a single predicted structure. RoseTTAFold2 and OmegaFold show much higher means and multimodal distributions, while DMPfold2 again exhibits the broadest and highest-RMSD profiles (means ∼ 0.41 *−* 0.45 Å; tails *>* 1.4 Å). The secondary-structure comparison clearly shows that *α*-helix and mixed peptides are predicted most accurately, while *β*-sheet-rich and disordered peptides systematically yield higher RMSDs across every model.

**Fig. 5:**
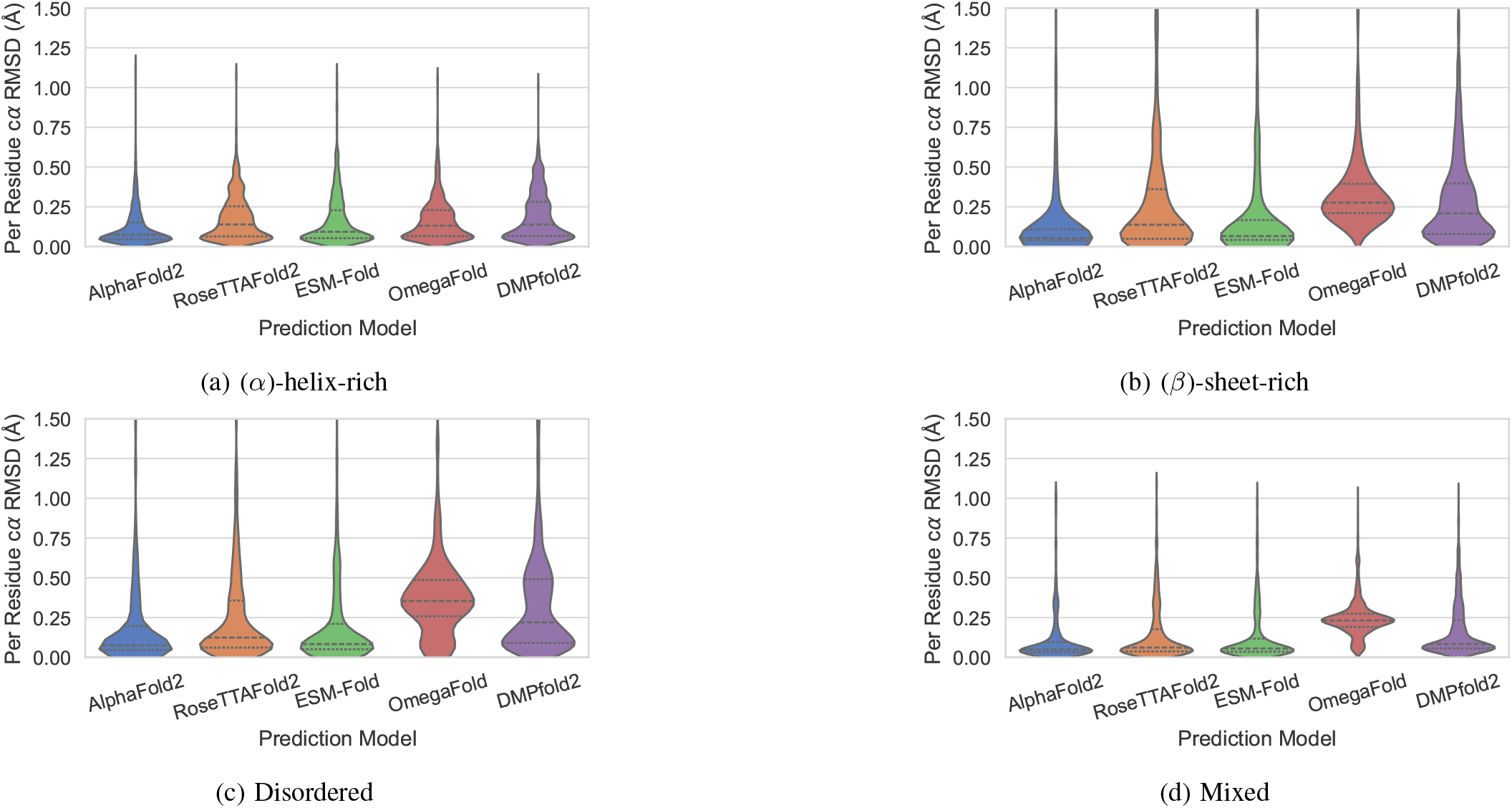
Secondary-structure-wise comparison of C(*α*) RMSD (Å) distributions across prediction models. Peptides were grouped into (*α*)-helix-rich, (*β*)-sheet-rich, disordered, and mixed classes. Lower RMSD values indicate better structural agreement with the experimental reference, while broader violin distributions reflect increased variability in model accuracy within a given structural class.

### B. Model Confidence vs. Local Accuracy (pLDDT/lDDT)

We quantify local structural accuracy using LDDT (local distance difference test). LDDT assesses, without global superposition, the fraction of a residue’s short-range interatomic distances (typically within a 15 Å cutoff) that are accurately reproduced in the model compared to the experimental structure. Scores are normalized to 0 *−* 1 and primarily reflect local geometry, specifically, the side-chain and backbone neighbourhoods, rather than overall global alignment. In contrast, pLDDT (predicted LDDT) represents the model’s self-reported confidence. This is a per-residue prediction of LDDT, often generated by the uncertainty head of AlphaFold-class models and their variants. OmegaFold was not included in the pLDDT-based calibration analysis because it does not report pLDDT in the same definition and calibration framework as AlphaFold2 and ESMFold. Although OmegaFold provides model-associated confidence information, this score is not directly equivalent to the predicted LDDT values used by AlphaFold2-class confidence heads. Including OmegaFold in the same pLDDT-LDDT comparison would therefore mix confidence measures with different definitions and calibration properties. For this reason, the LDDT-pLDDT correlation analysis was restricted to models that explicitly report pLDDT-like per-residue confidence scores comparable to the reference-based LDDT metric. A high predicted local distance difference test (pLDDT) score typically signifies stable local environments with minimal predicted errors. In contrast, low pLDDT values are commonly associated with flexible regions such as loops, termini, and flexible segments, or areas where the model’s confidence is low. A strong correlation between pLDDT and the measured LDDT suggests a well-calibrated prediction model. Conversely, miscalibration manifests as either high confidence in inaccurate regions (high pLDDT, low LDDT) or overly pessimistic confidence (low pLDDT, high LDDT). For AlphaFold2, as depicted in Figure 25, the correlation improves once peptide length exceeds ∼ 20 residues, increasing from 0.52 for 10-19 residues to 0.65 for 20-29 residues, before remaining moderately strong in the 30-39 bin (0.62) and decreasing again for the longest peptides (0.52 for 40-50 residues). The weaker correlation for very short peptides likely reflects the high conformational flexibility of short chains, where even models with similar confidence values can produce structurally diverse conformations. A similar pattern is observed for RoseTTAFold2, as depicted in Figure 29, with correlations of 0.52, 0.64, 0.67, and 0.53 across increasing length bins, showing the strongest calibration for peptides in the 30-39 residue range. For the single-sequence predictor ESMFold, the correlation gradually improves with peptide length, increasing from 0.57 for 10-19 residues to ∼ 0.68 for both the 20-29 and 30-39 bins, before slightly decreasing to 0.59 for the longest peptides, as depicted in Figure 33. This suggests that the language-model-derived confidence scores become more informative once sufficient sequence context is available. In contrast, as observed from Figure 39 DMPfold2 shows weaker and less stable correlations, with coefficients of 0.41, 0.60, 0.60, and 0.40 across the four length bins. From these observations, it can be inferred that none of the predictors achieve extremely strong correlations (*>* 0.8), indicating that while pLDDT/confidence is useful as a qualitative confidence indicator, it does not perfectly track true structural accuracy for short peptides, particularly at the extremes of the length range.

When we test the correlation of LDDT and pLDDT based on secondary structures, for AlphaFold2, the correlations remain relatively consistent across categories, ranging from 0.57 for *α*-helix-rich peptides to ∼ 0.56 for *β*-sheet-rich, 0.55 for disordered, and 0.53 for mixed structures, as depicted in Figure 26. This indicates that AlphaFold2’s pLDDT is moderately informative regardless of secondary-structure composition, although the spread of points is wider for disordered peptides, reflecting the intrinsic structural heterogeneity of these sequences. RoseTTAFold2 exhibits a similar pattern, as observed in Figure 30 but with slightly stronger calibration for *β*-sheet-rich peptides (*r ≈* 0.77), while *α*-helix-rich, disordered, and mixed peptides show moderate correlations around 0.56-0.62. For ESMFold, the correlations remain moderately strong across most classes, with coefficients of approximately 0.67 for *α*-helix-rich peptides, 0.72 for *β*-sheet-rich peptides, 0.62 for disordered peptides, and 0.53 for mixed structures, as depicted in Figure 34. This indicates that the language-model-derived confidence is generally well aligned with actual structural accuracy, particularly for structured motifs such as helices and sheets. However, the broader scatter observed in disordered and mixed peptides again reflects the difficulty of assigning reliable confidence to flexible or multi-state conformations. In contrast, as evident from Figure 40, DMPfold2 shows substantially weaker calibration, particularly for disordered peptides where the correlation drops to ∼ 0.21, while *α*-helix-rich and *β*-sheet-rich peptides show only modest correlations (∼ 0.50 and ∼ 0.54, respectively). The mixed class also exhibits weak association (∼ 0.38), suggesting that the scalar confidence score produced by DMPfold2 does not reliably track local structural correctness. When grouped by secondary-structure class, the correlation tends to be stronger for (*β*)-sheet-rich peptides, moderate for *alpha*-helical and mixed structures, and weakest for disordered peptides.

### C. Global Structural Similarity (TM-score & GDT-TS)

To assess how closely a predicted peptide structure matches its experimentally determined conformation, we used two widely adopted similarity metrics, TM-score and GDT-TS. The template modelling score (TM-score)[34] is a length-normalised measure of topological similarity between two protein structures. Unlike the root-mean-square deviation (RMSD), TM-score weights small distance errors more strongly than large ones and introduces a length-dependent scaling factor, making the score relatively independent of protein length. TM-scores range from 0 to 1; values above ∼ 0.50 indicate the two structures share the same fold, whereas scores below about 0.17 correspond to random structural similarity. The global distance test-total score (GDT-TS)[35] also measures structural similarity but is reported on a 0-100 scale. It superimposes the C-*α* atoms of a predicted and experimental structure and calculates the fraction of residues whose positions fall within various distance cut-offs (typically 1, 2, 4 and 8 Å), averaging these fractions to yield a final percentage. Higher GDT-TS values denote closer agreement between the predicted and experimental structures, and the metric is widely used in Critical Assessment of Methods of Protein Structure Prediction (CASP)[36] assessments to compare structure-prediction methods. Across all five models, prediction accuracy increased with peptide length, as observable from Figure 8. As depicted in Figure 6, for AlphaFold2, mean TM-scores increase from 0.31 ± 0.12 in the 10-19 residue bin to ∼ 0.59 ± 0.15 for 40-49 residue peptides, with corresponding GDT-TS values increasing from ∼ 66 to ∼ 69. RoseTTAFold2 follows a similar trend but performs slightly worse for shorter peptides: its mean TM-score rises from 0.27 ± 0.09 (10-19 residues) to 0.55 ± 0.15 (40-49 residues), with GDT-TS increasing from 59 to 64.

**Fig. 6:**
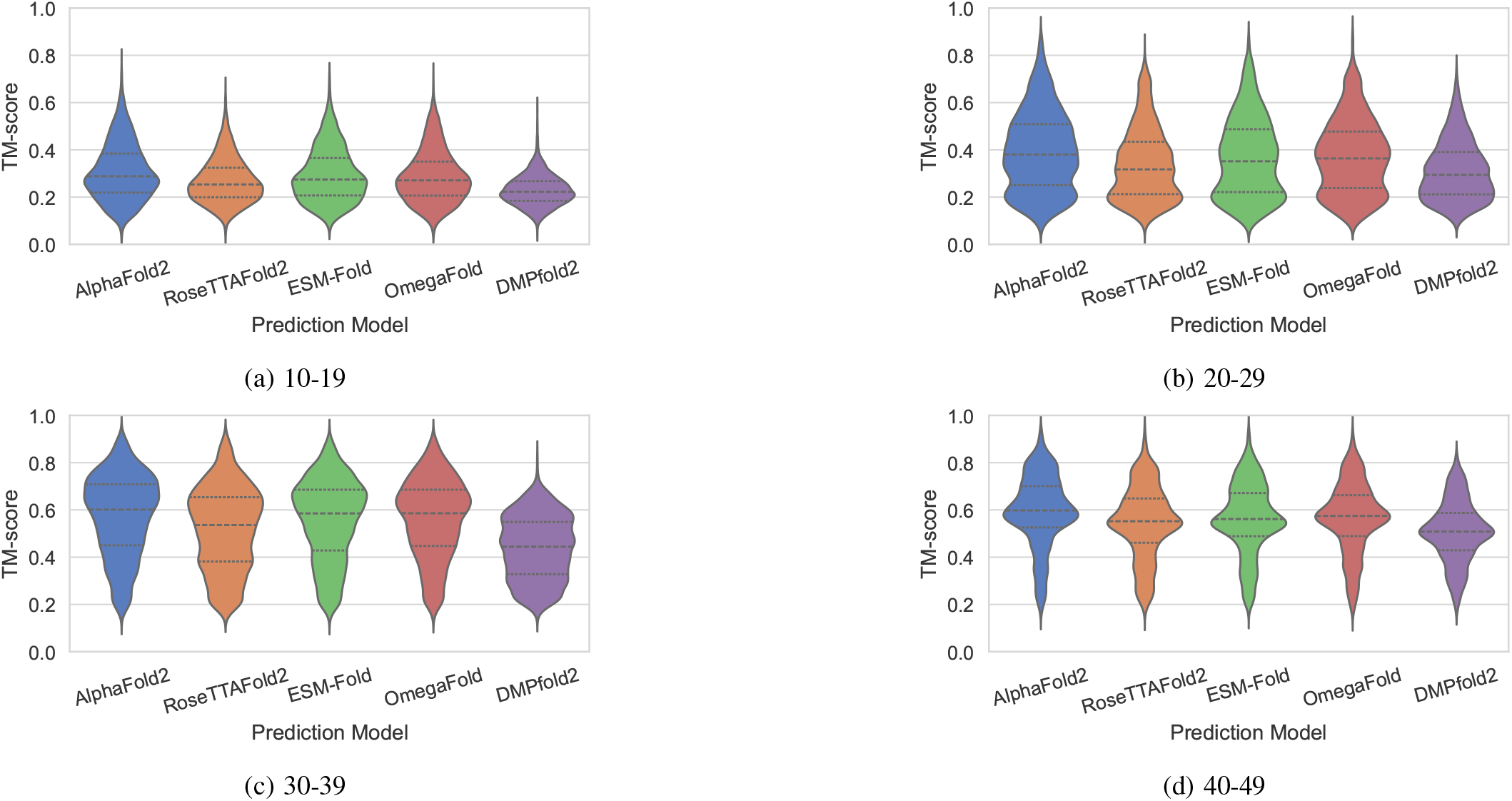
Length-wise comparison of TM-score distributions across prediction models. Violin plots show model-to-reference TM-scores for peptides grouped by sequence length. Higher TM-scores indicate better recovery of the overall structural topology, while lower scores indicate weaker global structural agreement.

For the single-sequence models, length dependence was more pronounced. ESMFold improves from 0.29 ± 0.11 in the shortest bin to 0.56 ± 0.15 in the longest, with GDT-TS increasing from 64 to 65. OmegaFold shows a similar improvement, with TM-scores rising from 0.29 ± 0.11 to 0.57 ± 0.15, while GDT-TS values increase from 63 to 66. In the intermediate length bins (30-49 residues), both models achieve TM-scores comparable to those of AlphaFold2 and RoseTTAFold2, suggesting that large protein language models can effectively compensate for the absence of evolutionary information when sufficient sequence context is available. In contrast, DMPfold2 lagged behind the other methods: TM-scores rarely exceeded 0.45 even for the longest peptides, and GDT-TS values remained below 70, although both metrics improved slightly with increasing length. These results suggest that methods using pre-trained language models (ESMFold and OmegaFold) can compensate for the absence of evolutionary information in longer peptides, whereas distance-geometry-based DMPfold2 struggles with these very short sequences. Classifying by secondary-structure class (*α*-helix-rich, *β*-sheet-rich, mixed, disordered) revealed a strong class effect that cut across models, as observed from Figures 12 and 10. For AlphaFold2, peptides containing well-defined secondary structures achieve the highest structural similarity scores. *β*-sheet-rich peptides show the strongest agreement with experimental structures, with a mean TM-score of 0.57 ± 0.21 and GDT-TS of 81.87 ± 14.47, followed closely by mixed and *α*-helical peptides (TM-scores 0.48-0.57 and GDT-TS 71-74). RoseTTAFold2 followed the same ordering with slightly lower medians in the structured classes. ESMFold and OmegaFold show similar structural trends but with slightly different strengths. ESMFold performs best for *β*-sheet-rich peptides (TM-score ∼ 0.54 ± 0.21; GDT-TS ~ 78), while *α*-helical and mixed peptides yield comparable but slightly lower scores (∼ 0.45-0.57 TM-score; ∼ 66-71 GDT-TS). OmegaFold also performs well on structured peptides, with *β*-rich and mixed classes showing the highest similarity (TM-score ∼ 0.53-0.59; GDT-TS ∼ 76-82). DMPfold2 was weakest overall but showed a modest relative advantage on *β*-rich peptides compared with *α*-rich, consistent with its constraint-driven modelling aligning more naturally with strand pairing than with short, solvent-exposed helices. These results highlight that MSA-dependent methods such as AlphaFold2 and RoseTTAFold2 provide reasonable models of 30-50 residue peptides but are only modestly accurate for peptides below 20 residues.

### D. Native Contact Evaluation

In addition to conventional structural metrics such as RMSD, we also evaluated prediction quality using native contact analysis, following the framework introduced by Best et al. [37]. In this approach, the structural similarity between a predicted model and the experimental structure is quantified by the fraction of native contacts (Q), defined as the proportion of residue-residue contacts present in the experimentally derived structure that are also formed in the predicted structure. Native contacts are typically defined using a distance threshold between heavy atoms of residue pairs in the reference structure, and Q therefore provides a measure of how well the predicted structure preserves the native contact network rather than only the coordinates. Using native contacts as an evaluation metric provides a physically meaningful complement to RMSD-based measures. While RMSD captures geometric deviations in Cartesian coordinates, it can be sensitive to local structural shifts or flexible regions that do not necessarily disrupt the overall fold. In contrast, the native contact fraction directly probes whether the correct residue-residue interactions and packing topology are reproduced, which are the key determinants of protein stability and folding mechanisms. Thus, we employ native contact analysis here as a physics-based structural descriptor, allowing us to assess whether predicted peptide structures reproduce the correct interaction network. As depicted in Figure 14, AlphaFold2 shows the strongest and most stable performance across all length bins, with mean Q values increasing slightly from 0.86 ± 0.11 (10-19 aa) to 0.92 ± 0.09 (40-49 aa), indicating reliable reconstruction of native residue-residue contacts even for the shortest peptides. ESMFold exhibits a very similar trend, with mean Q values ranging from 0.87 ± 0.10 to 0.91 ± 0.08, suggesting that its single-sequence language-model architecture is capable of recovering native contact patterns with accuracy comparable to AlphaFold2, particularly for peptides longer than 20 residues. RoseTTAFold2 also shows a clear improvement with increasing peptide length, with mean **Q** values rising from 0.82 ± 0.12 in the 10-19 aa bin to 0.92 ± 0.10 for 40-49 aa peptides, suggesting that the model benefits from additional sequence context when reconstructing residue-residue interactions, the slightly larger variance implies less consistent structural accuracy compared to AlphaFold2 and ESMFold. DMPfold2 consistently shows the lowest native-contact recovery across the dataset. Mean Q values decrease from 0.83 ± 0.11 (10-19 aa) to 0.79 ± 0.15 (30-39 aa), indicating that the model has greater difficulty reconstructing accurate interaction networks for short peptides.

When we consider dataset stratified based on secondary structure, across most models, *α*-helix-rich and mixed secondary-structure peptides show the highest native-contact fractions, as depicted in Figure 15, indicating that these structural classes are most reliably reconstructed. AlphaFold2 achieves mean Q values of ∼ 0.92 for both *α*-helical and mixed peptides, while ESMFold, while RoseTTAFold2 and OmegaFold show comparable values (0.91-0.92), suggesting that regular hydrogen-bonding patterns in backbone geometry, which is characteristic of helices are easier for current predictors to capture. DMPfold2, although slightly lower, still maintains reasonable contact recovery (0.87), indicating that even restraint-driven approaches can reconstruct the relatively local interaction pattern of helices. In contrast, *β*-sheet-rich peptides exhibit slightly lower and more variable contact recovery, particularly for RoseTTAFold2 and DMPfold2. The disordered peptide class shows the lowest native-contact fractions across all models, with Q values decreasing to 0.86 for AlphaFold2 and ESMFold, 0.83 for RoseTTAFold2, 0.86 for OmegaFold, and as low as 0.75 for DMPfold2. This behaviour is expected because intrinsically flexible peptides lack a single dominant contact network, making it inherently difficult for structure predictors to reproduce the experimentally observed interaction pattern.

### E. Effect of MSA on structure prediction

Multiple sequence alignments provide evolutionary information by aligning homologous sequences and identifying conserved positions, correlated mutations, and co-evolving residue pairs. In structure-prediction models, this information is useful because correlated substitutions can indicate spatial proximity between residues and therefore help define residue-residue contacts, secondary-structure packing, and global topology. However, the usefulness of MSA information is expected to be limited for short peptides, because they often have fewer homologous sequences, weaker co-evolutionary signal, and fewer long-range contacts than folded protein domains. To evaluate this effect, we compared AlphaFold2, RoseTTAFold2, and DMPfold2 predictions generated with and without MSA information. ESMFold and OmegaFold were not included in this MSA-ablation comparison because they do not rely on externally supplied MSAs during inference. Instead, both models are designed as single-sequence predictors. ESMFold uses representations from a large protein language model trained on massive sequence databases, allowing it to encode evolutionary and structural regularities directly from the input sequence without constructing an explicit MSA. OmegaFold similarly predicts structures from single sequences using learned sequence and pair representations, with geometric reasoning incorporated through its folding network rather than through direct MSA-derived co-evolutionary constraints. Therefore, ESMFold and OmegaFold provide useful MSA-independent baselines, whereas AlphaFold2, RoseTTAFold2, and DMPfold2 allow direct testing of how removing MSA information affects prediction quality.

For AlphaFold2, removing MSA information has only a modest effect in the shortest peptides but a clearer effect in longer sequences, as evident from Figures 7 and 9. In the 10-19 residue bin, the mean Q value remains unchanged at 0.86, and TM-score and GDT-TS remain very similar with and without MSA information. However, the effect becomes stronger as peptide length increases. In the 40-49 residue bin, Q decreases from 0.92 with MSA to 0.80 without MSA, TM-score decreases from 0.59 to 0.45, and GDT-TS decreases from 69.26 to 52.62. This indicates that AlphaFold2 can still recover many local structural features without MSA information, especially for very short peptides, but MSA-derived constraints become increasingly important for maintaining global topology in longer peptides. A similar but more pronounced trend is observed for RoseTTAFold2. With MSA information, RoseTTAFold2 shows increasing TM-score with peptide length, reaching 0.52 and 0.55 in the 30-39 and 40-49 residue bins, respectively. Without MSA information, these values fall to 0.41 and 0.39. The corresponding GDT-TS values also decrease substantially, from 68.68 to 56.00 in the 30-39 bin and from 63.99 to 47.04 in the 40-49 bin. Q values are also reduced in longer peptides, decreasing from 0.88 to 0.76 in the 40-49 residue bin. Thus, RoseTTAFold2 shows greater dependence on MSA information for preserving both contact-level and topology-level accuracy, particularly in longer peptides where nonlocal interactions and segment organization become more important. DMPfold2 shows the strongest dependence on MSA-derived information. In the shortest 10-19 residue bin, performance is relatively similar with and without MSA information, with Q values remaining at 0.83 and only minor differences in TM-score and GDT-TS. However, performance drops sharply in the longer bins when MSA information is removed. In the 30-39 residue bin, Q decreases from 0.79 to 0.70, TM-score decreases from 0.44 to 0.36, and GDT-TS decreases from 61.32 to 50.42. The effect is even stronger in the 40-49 residue bin, where Q decreases from 0.80 to 0.64, TM-score from 0.51 to 0.37, and GDT-TS from 60.37 to 43.93. This behaviour is consistent with the design of DMPfold2, which relies heavily on MSA-derived covariance or co-evolutionary constraints to infer residue-residue contacts and global structure. When this information is unavailable, its ability to reconstruct reliable peptide topologies is substantially reduced.

**Fig. 7:**
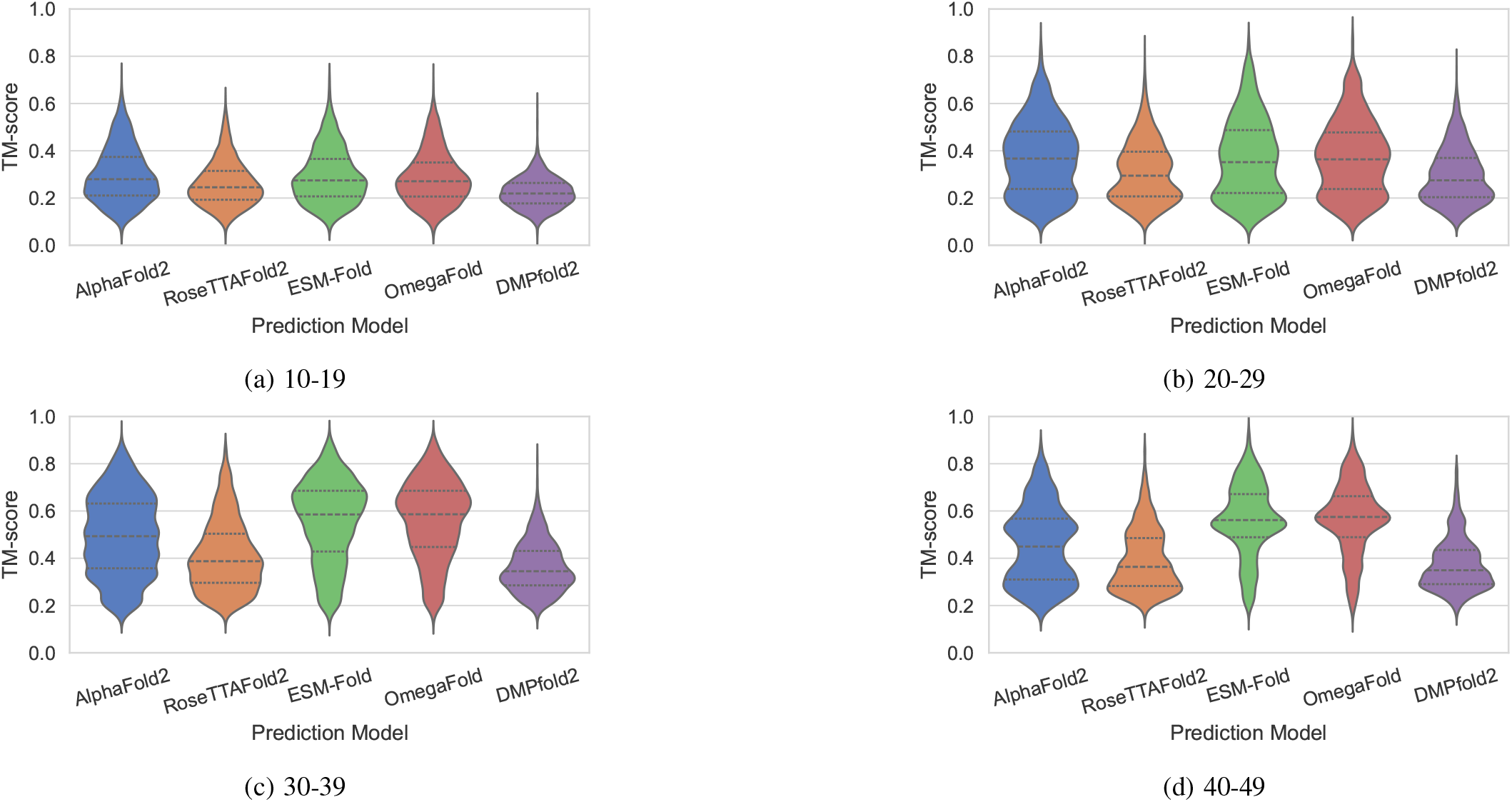
Length-wise comparison of TM-score distributions across prediction models. Violin plots show model-to-reference TM-scores for peptides grouped by sequence length. Higher TM-scores indicate better recovery of the overall structural topology, while lower scores indicate weaker global structural agreement. For AlphaFold2, RoseTTAFold2, and DMPfold2, predictions were generated using only the input peptide sequence, without providing multiple sequence alignment (MSA) information (i.e., single-sequence inference).

**Fig. 8:**
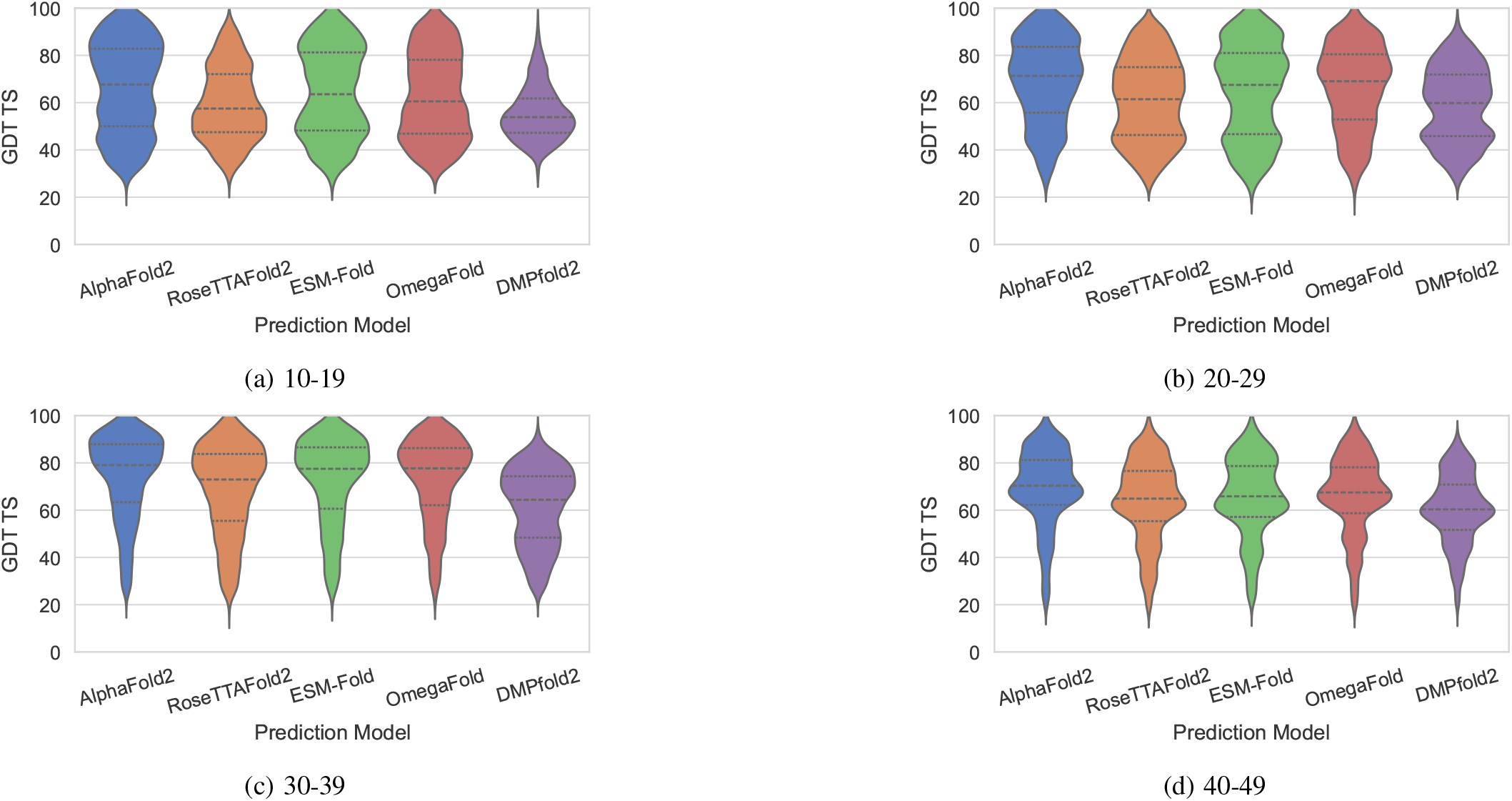
Length-wise comparison of GDT-TS distributions across prediction models. Violin plots show the distribution of model-to-reference GDT-TS values for each peptide length bin. Higher GDT-TS values indicate a larger fraction of residues superposed within standard distance thresholds, reflecting better global structural agreement.

**Fig. 9:**
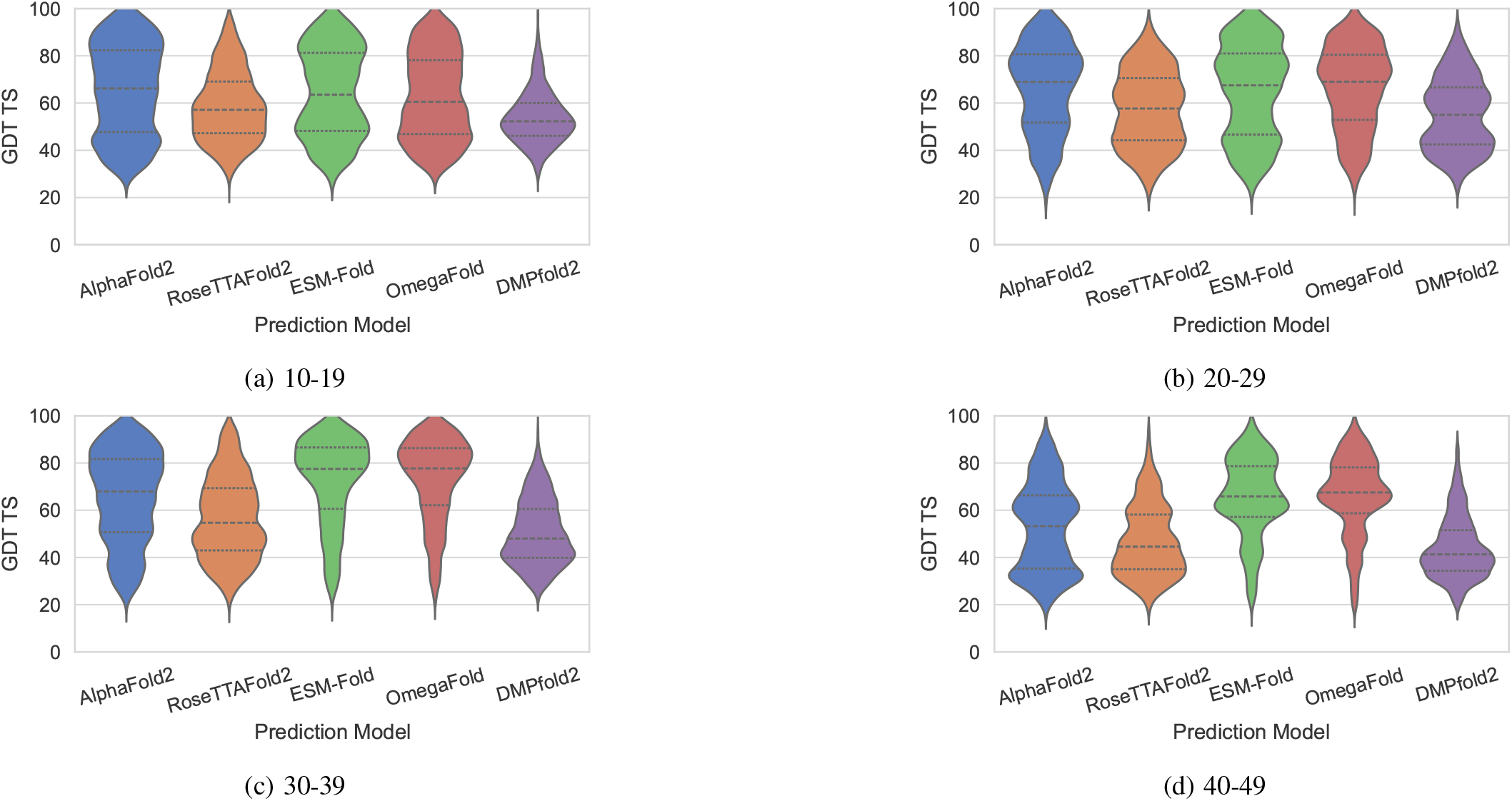
Length-wise comparison of GDT-TS distributions across prediction models. Violin plots show the distribution of model-to-reference GDT-TS values for each peptide length bin. Higher GDT-TS values indicate a larger fraction of residues superposed within standard distance thresholds, reflecting better global structural agreement. For AlphaFold2, RoseTTAFold2, and DMPfold2, predictions were generated using only the input peptide sequence, without providing multiple sequence alignment (MSA) information (i.e., single-sequence inference).

**Fig. 10:**
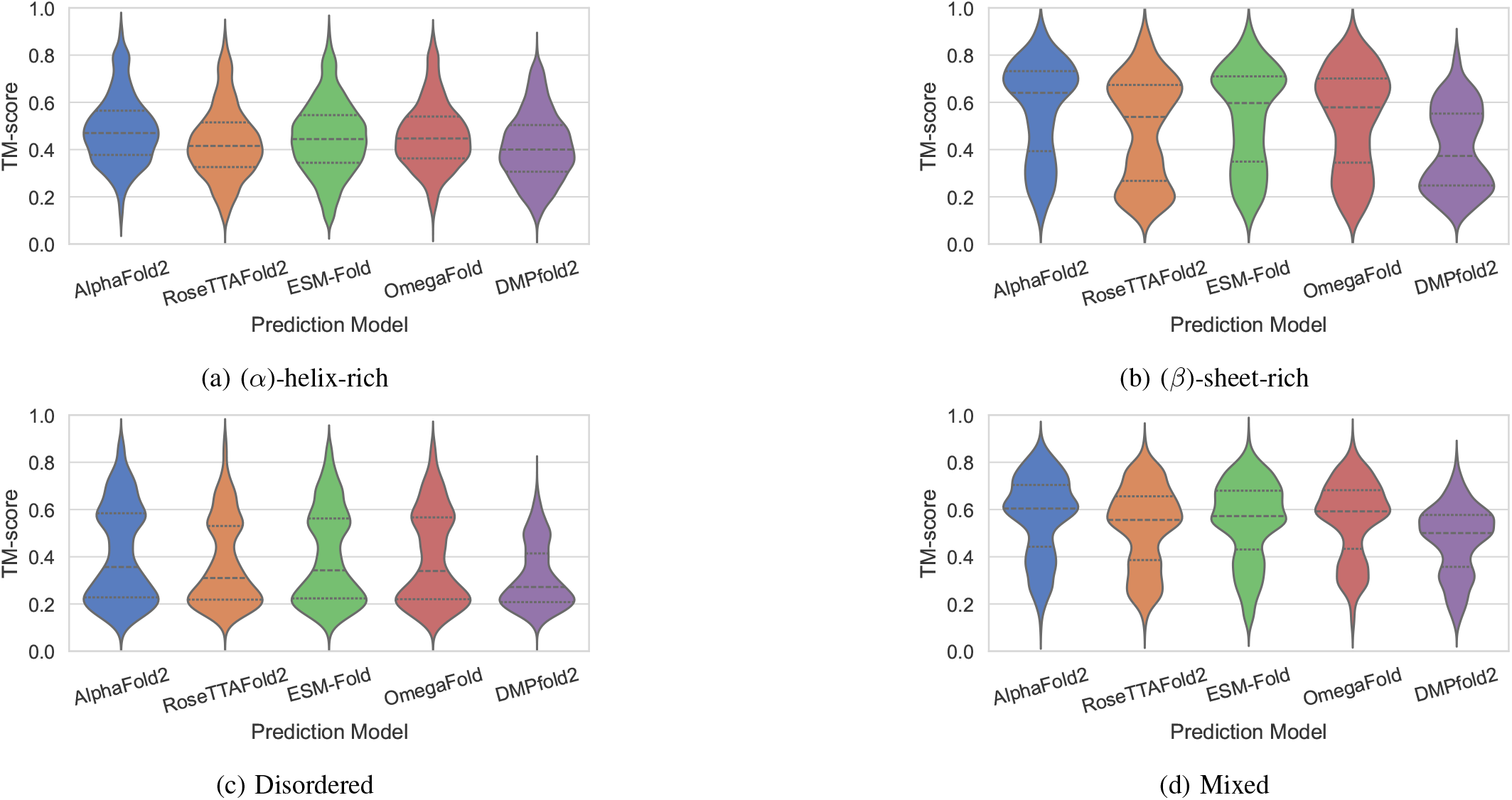
Secondary-structure-wise comparison of TM-score distributions across prediction models. Peptides were classified as (*α*)-helix-rich, (*β*)-sheet-rich, disordered, or mixed. Higher TM-scores indicate better global topological agreement with the experimental reference structure.

The secondary-structure-wise analysis further shows that the benefit of MSA information is not uniform across structural classes. For AlphaFold2, removal of MSA information reduces performance most clearly for *β*-rich, disordered, and mixed peptides, as evident from Figures 11 and 13. For example, *beta*-sheet-rich peptides show a decrease in GDT-TS from 81.87 with MSA to 73.76 without MSA, while mixed peptides decrease from 74.29 to 60.54. RoseTTAFold2 shows an even stronger MSA dependence in *β*-rich and mixed classes, where GDT-TS drops from 70.90 to 57.30 for *β*-rich peptides and from 69.44 to 54.40 for mixed peptides. DMPfold2 shows the largest secondary-structure-dependent degradation; *β*-rich peptides decrease from 64.36 to 53.33 in GDT-TS, disordered peptides from 53.04 to 45.36, and mixed peptides from 64.27 to 48.87. These trends suggest that MSA information is particularly important for peptides containing nonlocal contacts or multiple structural elements, whereas short *α*-helical peptides are less dependent on MSA because their geometry is largely determined by local sequence and backbone hydrogen-bonding propensities. The MSA-removal analysis shows that evolutionary information contributes most strongly when peptides are long enough to contain meaningful contact topology. For very short peptides, removing MSA information has relatively small effects, likely because these sequences contain limited co-evolutionary signal and are dominated by local structural propensities. In contrast, for longer, *β*-rich, mixed, or structurally complex peptides, MSA information improves contact recovery, TM-score, and GDT-TS, indicating better preservation of both local interaction networks and global topology. Among the tested models, AlphaFold2 appears most robust to MSA removal, RoseTTAFold2 shows intermediate dependence, and DMPfold2 is the most strongly affected. This supports the interpretation that MSA information is helpful but not uniformly decisive for short-peptide prediction; its value increases with peptide length, contact complexity, and secondary-structure organization.

**Fig. 11:**
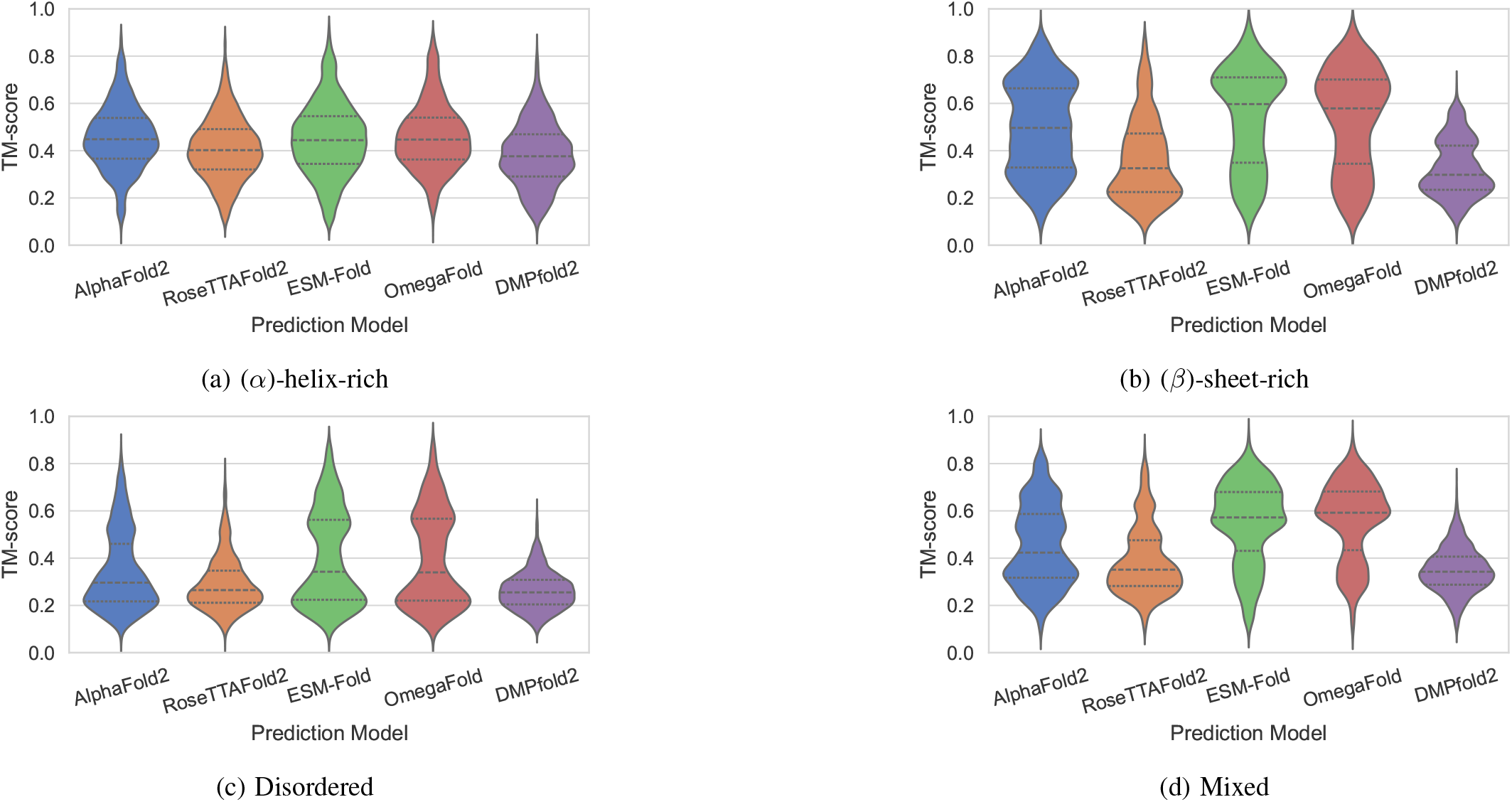
Secondary-structure-wise comparison of TM-score distributions across prediction models. Peptides were classified as *α*-helix-rich, *β*-sheet-rich, disordered, or mixed. Higher TM-scores indicate better global topological agreement with the experimental reference structure. For AlphaFold2, RoseTTAFold2, and DMPfold2, predictions were generated using only the input peptide sequence, without providing multiple sequence alignment (MSA) information (i.e., single-sequence inference).

**Fig. 12:**
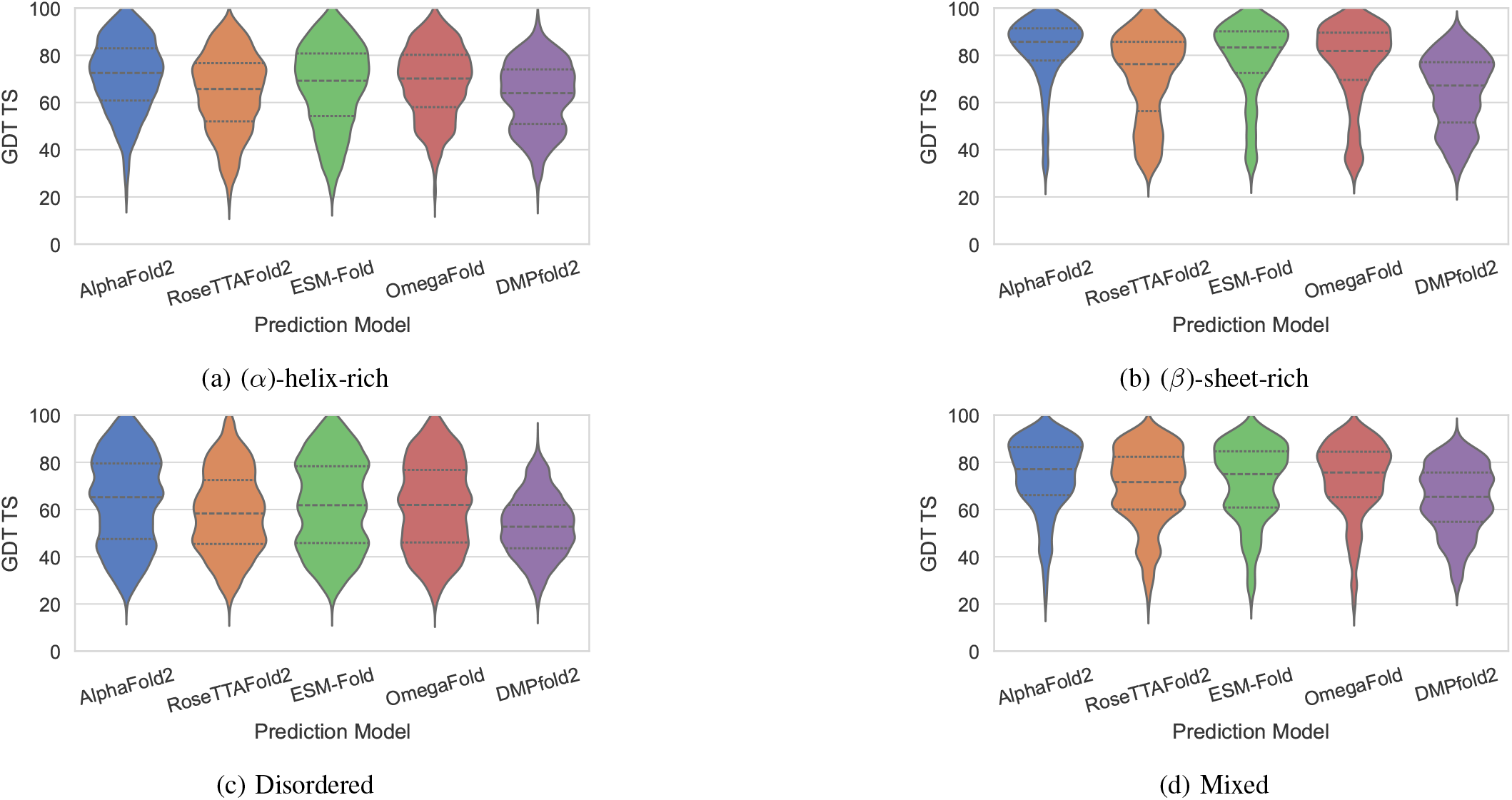
Secondary-structure-wise comparison of GDT-TS distributions across prediction models. Peptides were grouped by dominant structural class, and GDT-TS was used to quantify global agreement with the experimental reference. Higher values indicate better structural agreement, whereas broader distributions indicate greater variability in model performance within that class.

**Fig. 13:**
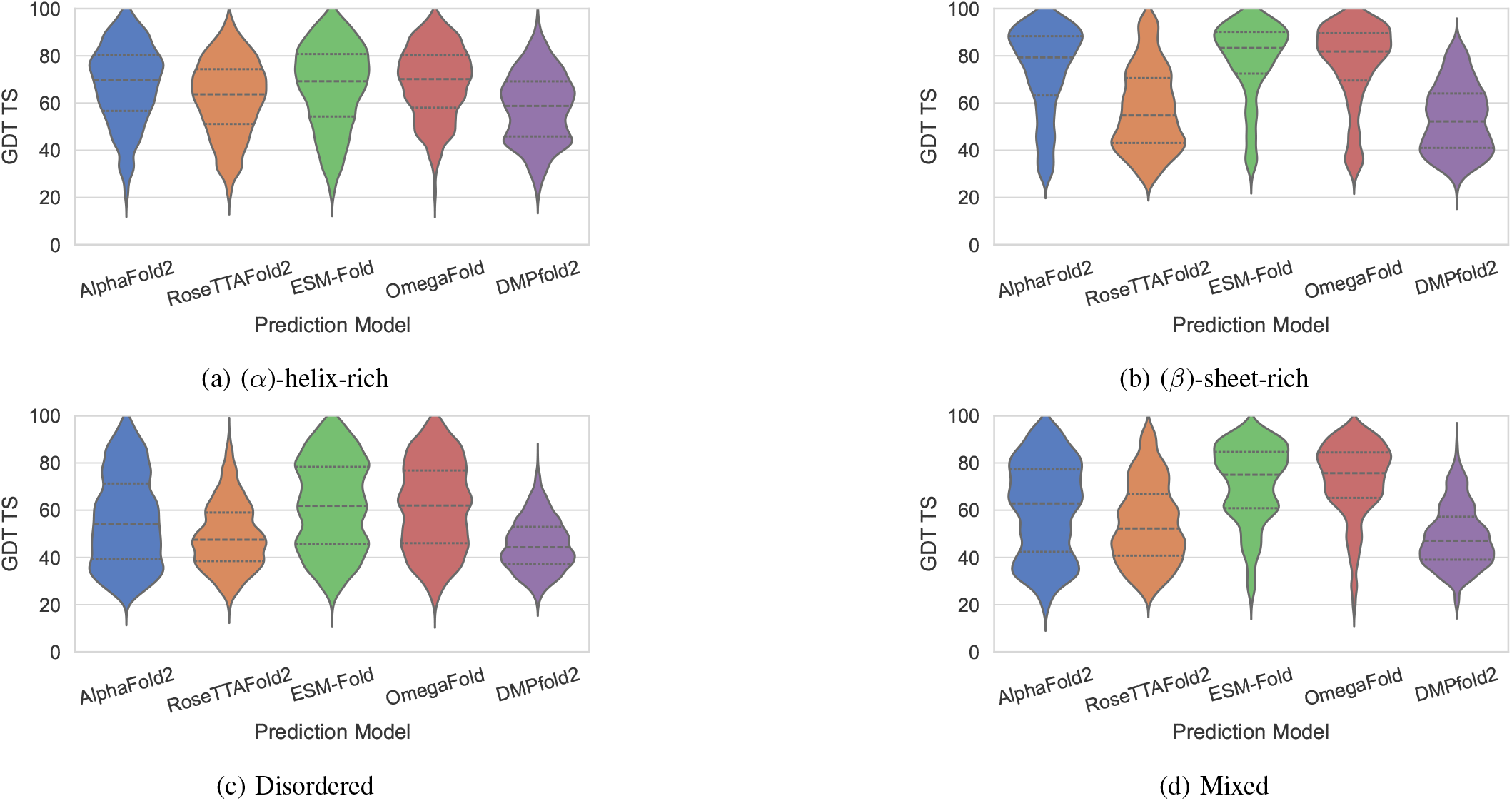
Secondary-structure-wise comparison of GDT-TS distributions across prediction models. Peptides were grouped by dominant structural class, and GDT-TS was used to quantify global agreement with the experimental reference. Higher values indicate better structural agreement, whereas broader distributions indicate greater variability in model performance within that class. For AlphaFold2, RoseTTAFold2, and DMPfold2, predictions were generated using only the input peptide sequence, without providing MSA information (i.e., single-sequence inference).

**Fig. 14:**
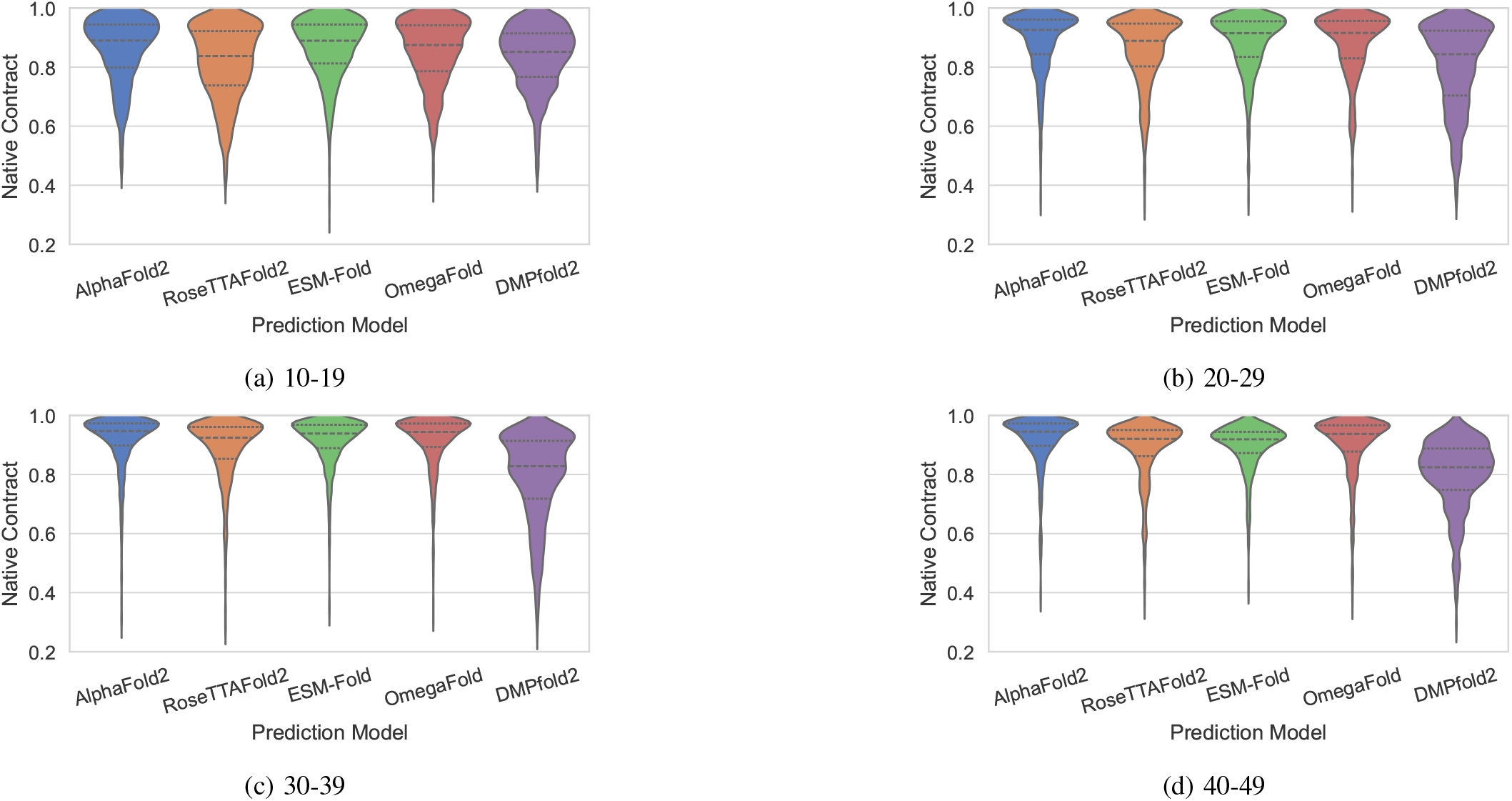
Length-wise comparison of native-contact preservation, (Q), across prediction models. Violin plots show the distribution of (Q) values within each peptide length bin. Higher (Q) values indicate greater recovery of the reference contact network, whereas lower values indicate poorer preservation of native residue-residue contacts.

**Fig. 15:**
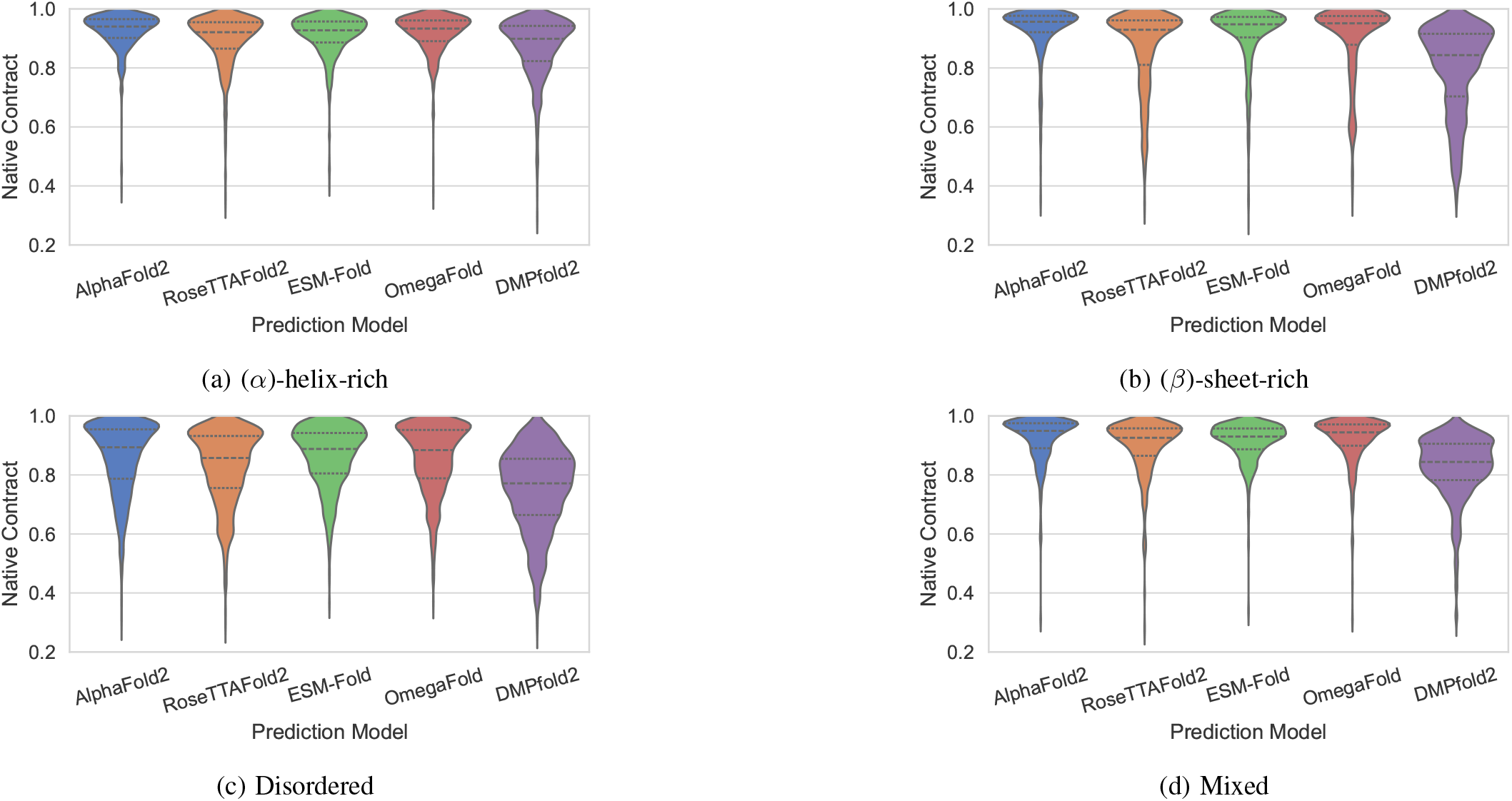
Secondary-structure-wise comparison of native-contact preservation, (Q), across prediction models. Peptides were grouped according to dominant secondary-structure class. Higher (Q) values indicate stronger preservation of the reference contact network, allowing comparison of how contact-level accuracy varies between helical, sheet-rich, mixed, and disordered peptides.

### F. dbAMP3 Prediction Correlation

To assess the practical applicability of the peptide-structure prediction workflow beyond experimentally solved PDB peptides, we extended the analysis to antimicrobial peptide sequences from the dbAMP3 [33] dataset. After filtering for sequence length, the final dataset contained 21,904 peptide sequences ranging from 10 to 50 residues. The average sequence length was 22.72 residues, with a median length of 20 residues. The N50 length was 25 residues, meaning that when peptides are sorted from longest to shortest, 50% of the total residue count is contained in peptides of length 25 residues or longer. Because experimental reference structures were not available for these dbAMP3 peptides, direct model-to-experiment accuracy assessment was not possible. Therefore, this analysis was designed as a model-referenced inter-model agreement study rather than a true accuracy benchmark. For each peptide, structures predicted by AlphaFold2, ESMFold, and RoseTTAFold2 were compared by selecting one model prediction as the reference and evaluating the structural agreement of the remaining predictions against it. AlphaFold2 was selected as the primary high-performing MSA-dependent reference model, ESMFold as a representative MSA-free protein-language-model predictor suitable for large-scale single-sequence peptide prediction, and RoseTTAFold2 as an independent MSA-dependent architecture distinct from AlphaFold2. RoseTTAFold2 was included as a deliberately independent comparator rather than as a presumed top-performing model. The PDB benchmark showed that RoseTTAFold2 was generally more separated from AlphaFold2 than ESMFold or OmegaFold, but this difference makes it useful in the dbAMP3 consensus analysis. Because the dbAMP3 dataset lacks experimental reference structures, the goal was not to rank predictors by accuracy, but to assess whether different modelling paradigms converged on similar peptide structures. RoseTTAFold2 provides an additional MSA-dependent but architecturally distinct reference point; therefore, cases where RoseTTAFold2 agrees with AlphaFold2 and ESMFold can be interpreted as stronger consensus predictions, while cases where it diverges highlight lower-confidence or architecture-sensitive structural assignments. The correlation was quantified using the same metrics as the rest; RMSD, TM-score, GDT-TS, and model-referenced native contact overlap. Because the contacts are derived from predicted structures rather than experimentally determined native states, they should not be described as true native contacts in this section. A more accurate term is model-referenced contact overlap or inter-model contact consistency. This design allows us to ask whether independent prediction frameworks converge on similar structural solutions. In this context, agreement between models should be interpreted as prediction consensus rather than proof of native correctness; nevertheless, consensus across architectures provides a useful measure of structural robustness when experimental references are absent.

Among the model comparisons, AlphaFold2 and ESMFold show the strongest coordinate-level agreement, with compact per-residue RMSD distributions across most length bins, as depicted in Figure 16. Their mean RMSD values remain low, ranging from 0.10 to 0.16 Å, with the lowest deviation observed in the 30–39 residue bin. This suggests that AlphaFold2 and ESMFold generally converge to similar backbone geometries across the dbAMP3 peptide set. Comparisons involving RoseTTAFold2 show slightly greater variability. AlphaFold2-RoseTTAFold2 RMSD values are higher for shorter peptides but decrease with increasing length, from approximately 0.21 Å in the 10-19 bin to 0.12 Å in the 40-49 bin, indicating improved agreement for longer sequences. RoseTTAFold2-ESMFold shows intermediate behaviour, with mean RMSD values remaining around 0.13-0.15 Å across length bins but with broader distributions. The RoseTTAFold2 provides a useful third-model check but introduces greater coordinate-level variability. RoseTTAFold2 comparisons show broader distributions, particularly in comparisons involving AlphaFold2 or ESMFold, suggesting greater model-to-model variability in backbone placement. Although the median per-residue RMSD remains low for many peptides, all comparisons retain long right-hand tails, especially in the longer 30-39 and 40-50 residue bins. These values indicate a subset of dbAMP3 peptides for which the predictors diverge substantially, likely because longer antimicrobial peptides can contain multiple helices, turns, flexible termini, or alternative packing arrangements that allow different models to converge on different conformational solutions. This per-residue RMSD trend should be interpreted together with TM-score and GDT-TS. Scenarios where per-residue RMSD remains elevated but TM-score and GDT-TS are high suggest that the models may agree on the overall peptide topology while differing in the exact spatial placement, bending, or register of structural elements. In this sense, the dbAMP3 RMSD analysis highlights coordinate-level uncertainty, whereas the topology-based metrics indicate whether the same general fold or structural organization is preserved across predictors.

**Fig. 16:**
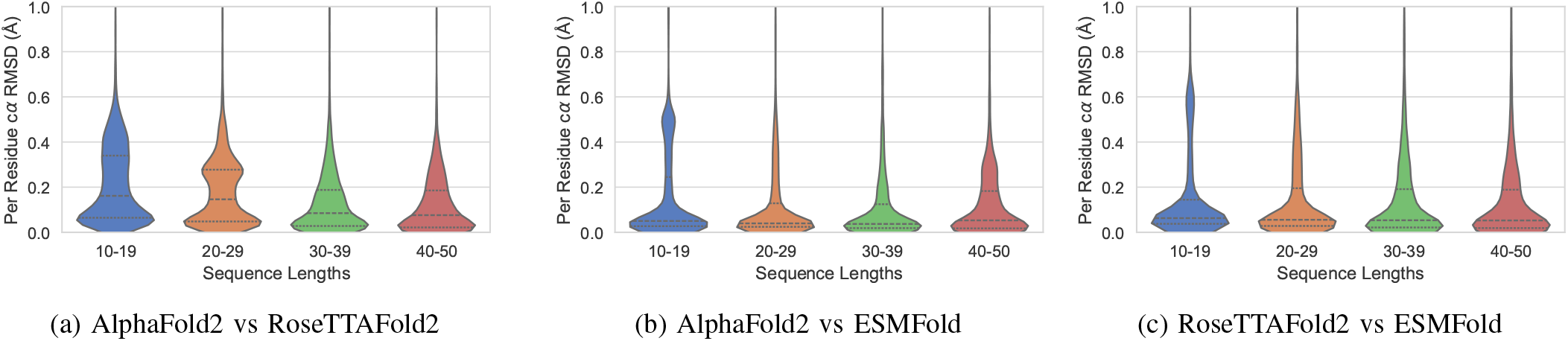
Pairwise comparison of RMSD distributions between peptide structures predicted by AlphaFold2, RoseTTAFold2, and ESMFold on the dbAMP3 dataset. Violin plots show the distribution of backbone root-mean-square deviation (RMSD) values calculated between the predicted structures generated by each pair of models, as no experimental reference structures are available for this dataset. Lower RMSD values indicate greater structural agreement between model predictions, whereas broader distributions reflect increased variability in the extent of agreement across peptides.

The topology-based metrics provide a complementary view to the per-residue RMSD analysis. As shown above, AlphaFold2-ESMFold showed the strongest coordinate-level agreement, with mean per-residue RMSD values decreasing with increasing peptide length and remaining consistently low for the longer peptides. While per-residue RMSD captures fine coordinate-level disagreement between model predictions, TM-score and GDT-TS are more tolerant of local shifts and instead report whether two structures preserve a similar overall topology. The topology-based metrics support this agreement, with TM-scores increasing across the length bins and GDT-TS values remaining consistently high across all peptide lengths. across the four length bins, suggesting that these two models often converge on similar dbAMP3 peptide conformations. In the dbAMP3 dataset, TM-score and GDT-TS generally improve with increasing peptide length, particularly in the 30-39 and 40-50 residue bins, as depicted in Figures 17 and 19, suggesting that longer antimicrobial peptides provide sufficient structural context for AlphaFold2, RoseTTAFold2, and ESMFold to converge on broadly similar global conformations. This trend contrasts with the RMSD distributions, which remain broader in the longer length bins. The apparent discrepancy is not contradictory; it indicates that models may agree on the global organization of a peptide while differing in local backbone placement, helix bending, terminal flexibility, or the relative orientation of structural segments. This interpretation is especially relevant for antimicrobial peptides, which are often amphipathic, flexible, and environment-dependent, and may adopt different conformations depending on solvent, membrane association, oligomeric state, or binding context.

**Fig. 17:**
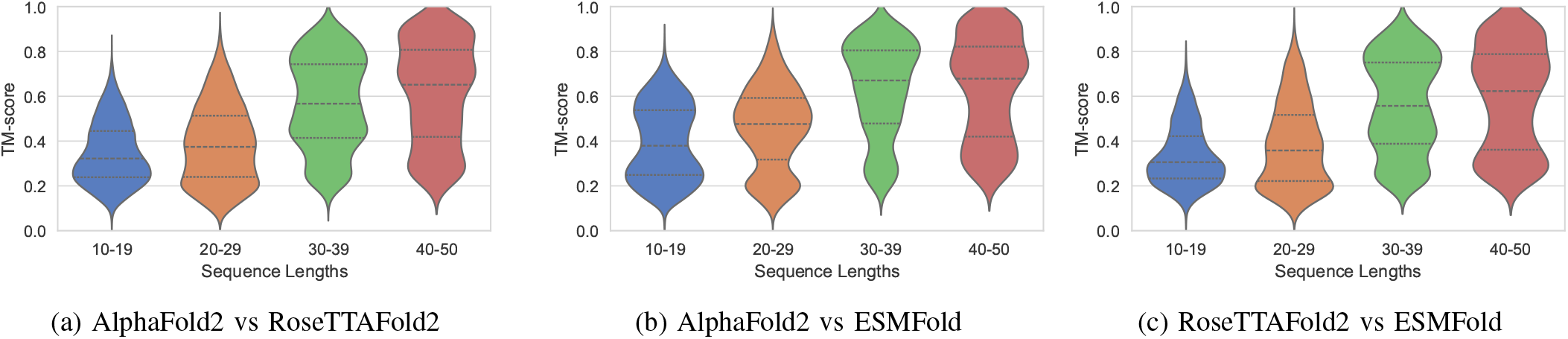
Pairwise comparison of TM-score distributions between peptide structures predicted by AlphaFold2, RoseTTAFold2, and ESMFold on the dbAMP3 dataset. Violin plots show the distribution of TM-scores calculated between the predicted structures generated by each pair of models, as no experimental reference structures are available for this dataset. Higher TM-scores indicate greater agreement in the overall structural topology between model predictions, whereas broader distributions reflect increased variability in inter-model structural similarity across peptides.

Contact-overlap analysis provides an intermediate level of comparison between coordinate-sensitive RMSD and topology-sensitive TM-score or GDT-TS. The contact-overlap distributions remain concentrated at high values across most length bins, indicating that AlphaFold2, RoseTTAFold2, and ESMFold frequently preserve similar local interaction patterns. Among the three comparisons, AlphaFold2-ESMFold again shows the strongest contact-level agreement, with mean Q values of 0.91 ± 0.10, 0.94 ± 0.08, 0.94 ± 0.08, and 0.92 ± 0.09 across the 10-19, 20-29, 30-39, and 40-49 residue bins, respectively, as depicted in Figure 18. The corresponding median values remain very high, ranging from 0.95 to 0.97 across length bins, indicating that most AlphaFold2 and ESMFold predictions preserve highly similar local contact patterns. Comparisons involving RoseTTAFold2 also show high contact overlap, although with slightly lower values. For AlphaFold2-RoseTTAFold2, mean Q increases from 0.85 ± 0.14 in the 10-19 bin to 0.92 ± 0.10 and 0.91 ± 0.10 in the 30-39 and 40-49 bins, respectively. RoseTTAFold2-ESMFold shows a similar but slightly lower pattern, with mean Q values of 0.86 ± 0.12, 0.88 ± 0.12, 0.90 ± 0.11, and 0.88 ± 0.11 across the four length bins. For dbAMP3 peptides, the combined patterns observed across the metrics indicate that the predictors often agree on local structural organization and overall topology, but not necessarily on precise backbone geometry. High contact overlap suggests that similar residue-residue interaction patterns are preserved across models, while improved TM-score and GDT-TS indicate convergence toward comparable global folds. In contrast, broader per-residue RMSD distributions reveal residual uncertainty in the exact placement, bending, or orientation of structural elements. The dbAMP3 peptides showing strong agreement across contact overlap, TM-score, and GDT-TS can be considered more robust structural hypotheses, whereas cases with high RMSD divergence despite reasonable topology scores should be treated as structurally ambiguous and may require experimental validation or ensemble-based modelling.

**Fig. 18:**
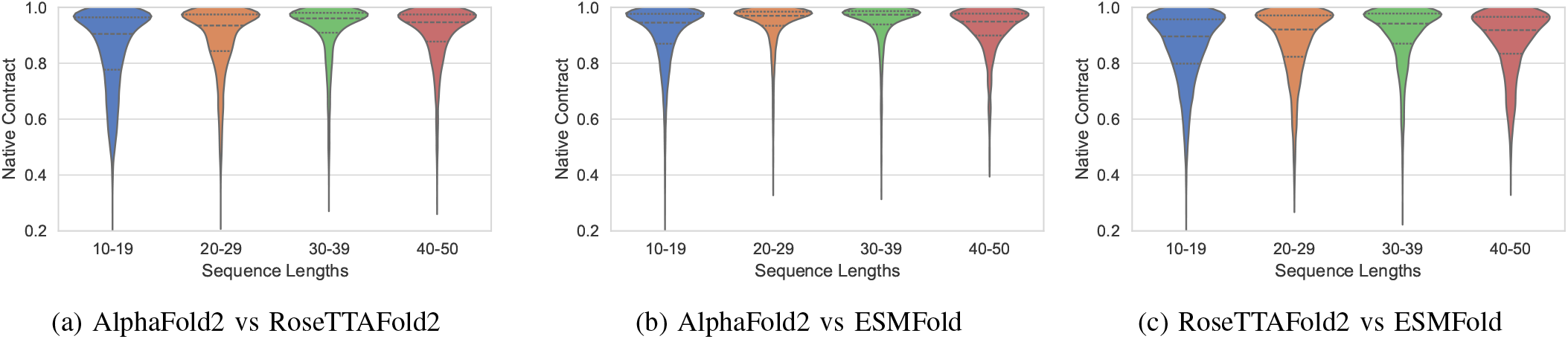
Pairwise comparison of native-contact preservation (Q) between peptide structures predicted by AlphaFold2, RoseTTAFold2, and ESMFold on the dbAMP3 dataset. Violin plots show the distribution of (Q) values calculated between the predicted structures generated by each pair of models, as no experimental reference structures are available for this dataset. Higher (Q) values indicate greater agreement in residue–residue contact patterns between model predictions, whereas lower values indicate weaker preservation of contact networks across models.

**Fig. 19:**
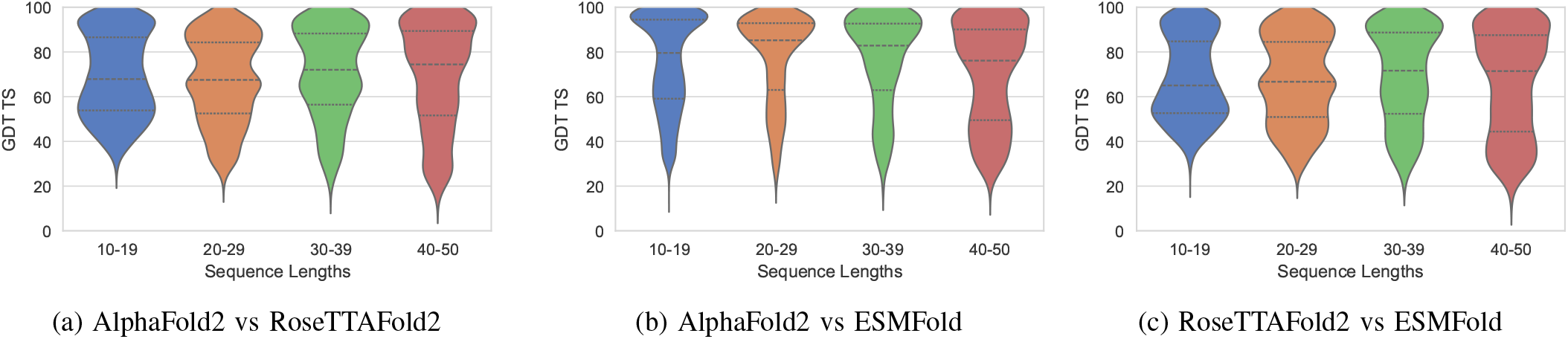
Pairwise comparison of GDT-TS distributions between peptide structures predicted by AlphaFold2, RoseTTAFold2, and ESMFold on the dbAMP3 dataset. Violin plots show the distribution of GDT-TS values calculated between the predicted structures generated by each pair of models, as no experimental reference structures are available for this dataset. Higher GDT-TS values indicate greater agreement in global backbone superposition between model predictions, whereas lower values indicate weaker structural consistency across models.

**Fig. 20:**
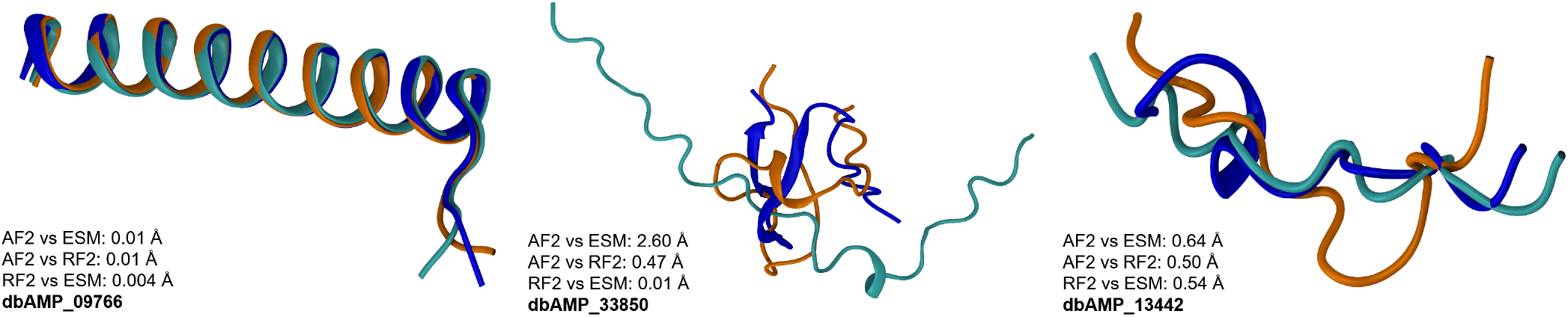
Aligned structural representations of three selected peptides from the dbAMP3 dataset. Structures predicted by AlphaFold2 (AF2), RoseTTAFold2 (RF2), and ESMFold (ESM) are shown in superposition to visually compare predicted conformations across models. AF2 structures are shown in blue, RF2 in orange, and ESMFold in cyan.

**Fig. 21:**
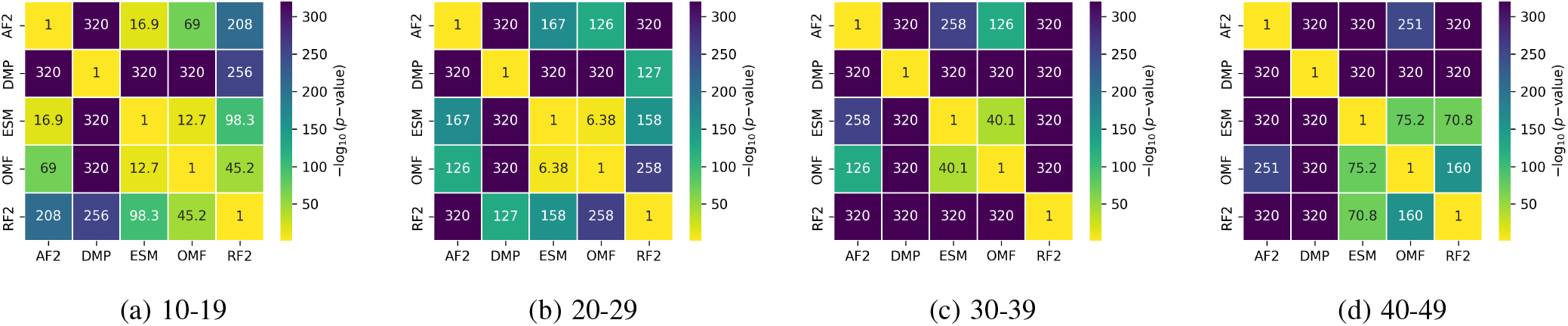
Wilcoxon signed-rank test significance heatmap for pairwise model comparisons using TM-score across peptide length bins.

**Fig. 22:**
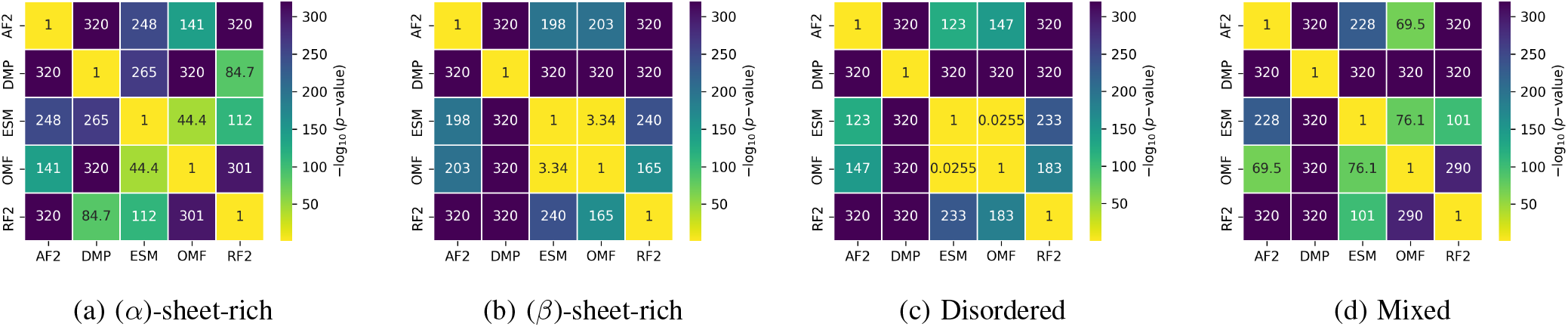
Wilcoxon signed-rank test significance heatmap for pairwise model comparisons using TM-score across secondary-structure classes.

The pairwise comparisons reveal that AlphaFold2 and ESMFold generally show the strongest agreement, especially in topology-sensitive metrics. Their per-residue RMSD distributions remain relatively compact across most length bins, while TM-score and GDT-TS increase markedly for longer peptides, indicating that these two models frequently recover similar global structures despite being based on different principles; AlphaFold2 relies on MSA-derived evolutionary information, whereas ESMFold uses single-sequence language-model representations. This cross-architecture agreement is particularly important because it suggests that the predicted fold is not merely an artifact of one modelling strategy. In contrast, comparisons involving RoseTTAFold2; particularly, AlphaFold2 versus RoseTTAFold2 and RoseTTAFold2 versus ESMFold, show broader RMSD distributions and more pronounced tails, suggesting greater variability in coordinate-level placement. However, the corresponding TM-score and GDT-TS distributions still improve with peptide length, indicating that RoseTTAFold2 often agrees with the other models at the level of broad topology even when the exact backbone geometry differs.

The combined behaviour of RMSD, TM-score, GDT-TS, and contact overlap provides a practical way to rank confidence in dbAMP3 predictions. Peptides for which AlphaFold2, ESM-Fold, and RoseTTAFold2 all show low per-residue RMSD, high TM-score, GDT-TS, and high contact overlap can be considered high-confidence structural hypotheses, because independent models converge on both local contacts and global topology. Peptides with high TM-score, GDT-TS and high contact overlap but broader RMSD should be treated as moderate-confidence cases; the models likely agree on the overall fold and residue-interaction network, but differ in helix bending, terminal placement, loop geometry, or relative orientation of structural segments. Finally, peptides with poor agreement across RMSD, TM-score, GDT-TS, and contact overlap should be regarded as structurally ambiguous, reflecting either genuine conformational heterogeneity or model uncertainty. Hence, in the absence of experimentally reported structures, multi-model agreement provides a rational confidence filter; the most reliable dbAMP3 structural predictions are those supported simultaneously by AlphaFold2-ESMFold agreement and by RoseTTAFold2 as an independent third-model check.

If AlphaFold2, ESMFold, and RoseTTAFold2 all converge to similar structures, the prediction can be considered a high-confidence structural hypothesis, because multiple architectures recover similar local contacts and global topology. If AlphaFold2 and ESMFold agree strongly but RoseTTAFold2 differs, the prediction may still be considered moderately robust, but the discrepancy suggests possible sensitivity to modelling architecture or MSA-derived features. Conversely, if RoseTTAFold2 and ESMFold agree while AlphaFold2 disagrees, or if AlphaFold2 and RoseTTAFold2 agree while ESMFold disagrees, the case should be treated more cautiously because the agreement may reflect either shared MSA-driven bias or alternative sequence-based interpretation. Thus, the most reliable dbAMP3 predictions are not those supported by any single pair alone, but those showing consistent agreement across all the metrics classes, low per-residue RMSD, high TM-score, GDT-TS, and high model-referenced contact overlap across multiple model pairs.

This multi-model comparison provides a practical confidence filter for dbAMP3 peptides. High contact overlap indicates that models preserve similar residue-residue interaction networks, high TM-score and GDT-TS indicate convergence toward similar global topology, and low per-residue RMSD indicates close coordinate-level agreement. When all three measures agree across AlphaFold2, ESMFold, and RoseTTAFold2, the predicted structure is likely to be a robust structural hypothesis. When TM-score, GDT-TS and contact overlap remain high but RMSD is elevated, the models may agree on the overall fold while differing in helix bending, terminal placement, or local backbone geometry. In contrast, peptides showing poor agreement across multiple model pairs should be considered structurally ambiguous and may require experimental validation, membrane-associated modelling, or ensemble-based conformational analysis.

## VI. Discussion

The accurate structural prediction of short peptides remains a distinct and formidable challenge in computational structural biology. While modern deep-learning models have revolutionized the prediction of large, globular proteins, their application to short sequences tests the limits of algorithms trained primarily on cooperative, well-folded domains. Ultimately, our findings highlight the critical interplay between sequence length, structural class, and evolutionary information, revealing how these models behave when forced to navigate the shallow, highly flexible conformational landscapes characteristic of short peptides.

### A. The Impact of Peptide Length and Structural Cooperativity

In this work, we systematically evaluated how modern structure prediction methods perform in this regime. Across all evaluated models, prediction accuracy improves systematically with peptide length. RMSD decreases while TM-score and GDT-TS increase as peptide length grows from 10–19 residues to 40–49 residues. This trend is particularly evident for AlphaFold2, Omegafold, and ESMFold, where longer peptides exhibit narrower RMSD distributions and fewer catastrophic outliers. The improvement likely arises because longer peptides possess increased opportunities for intra-chain stabilization, including transient tertiary contacts and more persistent secondary-structure motifs, resulting in deeper and more learnable free-energy minima.

### B. Challenges Across Secondary-Structure Classes

The secondary-structure analysis shows that model performance is strongly dependent on structural class. *α*-helical and mixed peptides generally exhibit lower RMSD and stronger native-contact recovery, indicating that local backbone geometry and short-range interaction patterns are more consistently reproduced. This is expected because helices are largely stabilized by local sequence and backbone hydrogen-bonding patterns, making them easier for modern predictors to recognize.

In contrast, *β*-sheet-rich and disordered peptides consistently represent the most difficult classes for all evaluated prediction methods. In *β*-sheet-rich peptides, predictors must correctly infer strand pairing, registry, and long-range hydrogen-bond topology despite limited sequence length and weak coevolutionary signals. Consequently, these systems exhibit broader RMSD distributions and increased structural variability, particularly for OmegaFold and DMPfold2. Disordered peptides present an even greater challenge because their experimentally observed structures often correspond to conformational ensembles rather than a single well-defined native state. For such peptides, a static predicted structure may represent only one possible conformation, resulting in consistently poorer agreement across models. These findings highlight an important limitation of current predictors: they remain strongly biased toward compact, ordered structural states learned predominantly from globular proteins.

One interesting emerging trend is that RMSD and native contact fraction (Q) indicate higher prediction quality for *α*-helix-rich and mixed peptides, whereas TM-score and GDT-TS tend to show better agreement for *β*-sheet-rich structures. This difference likely arises from the distinct sensitivities of the metrics. RMSD and native-contact fraction are more sensitive to local geometry and local packing, whereas TM-score and GDT-TS are more forgiving towards local errors and reward correct global topology. For *α*-helical and mixed peptides, predictors often get the local secondary structure right; as helices have regular *ϕ/ψ* geometry, predictable local hydrogen-bonding. By contrast, TM-score and GDT-TS emphasize global topological similarity after structural superposition rather than exact coordinate-level agreement. As a result, even when their local helical geometry are reproduced well, small differences in helix length, curvature, or orientation can lower TM-score and GDT-TS, which are more sensitive to overall fold agreement than to local secondary-structure correctness. *β*-sheet-rich peptides, on the other hand are typically more challenging to reproduce at the fine-grained level because strand registry and hydrogen-bond pairing are sensitive to small positional errors, which can increase RMSD and reduce native-contact agreement. However, once the model captures the overall *β*-hairpin or sheet topology, these metrics reward the correct global arrangement of strands even if local registry or individual contacts are imperfect. As a result, *β*-rich peptides can achieve comparatively higher TM-score and GDT-TS values despite somewhat poorer RMSD or contact-level agreement.

### C. Evolutionary Information vs. Language Models

Among the evaluated predictors, AlphaFold2 and ESMFold show the most consistent overall performance across the experimentally benchmarked peptide set. AlphaFold2 benefits from evolutionary and pairwise geometric information when sufficient signal is available. Interestingly, our results indicate that MSA-free predictors such as ESMFold and OmegaFold become increasingly competitive with MSA-dependent methods as peptide length increases. While AlphaFold2 generally achieves the most stable performance across all length bins, ESMFold and OmegaFold approach comparable TM-scores and GDT-TS values in the 30–49 residue range.

This observation suggests that large protein language models encode substantial structural priors directly from sequence statistics, enabling accurate prediction even in low-homology regimes where evolutionary depth is limited. However, for very short peptides (10–19 residues), the lack of sufficient sequence context reduces the effectiveness of language-model embeddings, leading to broader structural variability and higher failure rates compared with AlphaFold2. DMPfold2 remains the least consistent across most metrics, suggesting that its restraint- or constraint-based prediction strategy is less robust for short peptides, especially when long-range contacts and strong coevolutionary constraints are sparse.

### D. The Limits of Confidence Calibration

The confidence-calibration analysis further emphasizes that predicted confidence should be interpreted cautiously for short peptides. Although pLDDT and related confidence measures correlate moderately with experimentally derived LDDT, none of the evaluated methods achieve strong calibration in this regime. Correlation coefficients generally remain within the 0.5–0.7 range, indicating that confidence estimates are informative but imperfect predictors of structural correctness.

Calibration improves for peptides between 20 and 39 residues, where structural motifs become more stable and conformational heterogeneity decreases. However, performance deteriorates for highly disordered peptides and very short sequences, where multiple low-energy conformations coexist. In these systems, predictors frequently assign high confidence to structures that represent only one plausible conformational snapshot, such as a locally ordered helix, rather than the experimentally observed ensemble. These observations suggest that current confidence metrics are optimized primarily for folded proteins with unique native states and may not fully capture ensemble uncertainty in flexible peptide systems.

### E. Consensus-Based Structural Robustness in dbAMP3

The dbAMP3 analysis extends these conclusions to a biologically relevant peptide dataset lacking experimentally used 3D references. Because direct model-to-experiment validation was not possible for these antimicrobial peptides, the dbAMP3 analysis was designed as a consensus-based structural robustness assessment. Instead of asking whether a model exactly reproduces an experimental structure, the analysis asks whether independent predictors converge on similar structural solutions. This distinction is important. Agreement among AlphaFold2, RoseTTAFold2, and ESMFold does not prove that the predicted conformation is native, but it increases confidence that the structure represents a robust model-derived hypothesis. Conversely, strong disagreement among predictors indicates structural ambiguity and suggests that the peptide may be flexible, context-dependent, or sensitive to model assumptions.

The combined dbAMP3 metrics provide a practical framework for interpreting predicted antimicrobial peptide structures. Per-residue RMSD reports fine coordinate-level agreement, TM-score and GDT-TS report broader topological similarity, and model-referenced contact overlap reports preservation of local residue-residue interaction patterns. Cases where all three models show low RMSD, high TM-score, GDT-TS, and high contact overlap can be treated as high-confidence structural hypotheses. Cases with high TM-score, GDT-TS and high contact overlap but broader RMSD should be considered moderate-confidence predictions, where the models likely agree on the overall fold and local interaction network but differ in helix bending, terminal placement, or the relative orientation of structural elements. Finally, peptides with poor agreement across multiple metrics and model pairs should be treated as structurally ambiguous and may require experimental validation, membrane-associated modelling, or ensemble-based simulation. This consensus-based strategy is particularly relevant for antimicrobial peptides. Many AMPs are amphipathic, flexible, and environment-dependent. Their structures can change upon membrane binding, oligomerization, changes in solvent polarity, or interaction with lipid headgroups. Therefore, a single predicted structure in aqueous or model-free conditions may not represent the biologically active conformation. Multi-model agreement can help prioritize peptides whose predicted structures are robust across algorithms, but it cannot replace experimental or environment-specific validation. For AMP design workflows, the most useful approach is therefore not to accept every predicted structure as a final answer, but to use cross-model agreement as a filtering step before downstream molecular dynamics, membrane simulations, docking, or experimental characterization.

### F. Interpretation of Wilcoxon Test Results

The Wilcoxon signed-rank test with Holm-adjusted p-values [38], [39] was used to assess whether observed differences in model performance were statistically significant rather than due to random variation, revealing that model ranking is not fixed across peptide sizes. Figure 43 and 44 present the heatmaps for TM-score, while GDT-TS and RMSD is presented in Section G of the Appendix. The strongest and most consistent separation from AlphaFold2 was observed for DMPfold2, followed by RoseTTAFold2, indicating that these models form a lower-performing tier across most length regimes. In contrast, the separation between AlphaFold2 and the two single-sequence predictors, ESMFold and OmegaFold, was more nuanced. ESM-Fold remains one of the strongest overall models, particularly when considering the full benchmark, but the longer peptide bins reveal that OmegaFold becomes increasingly competitive. For peptides longer than approximately 29 residues, OmegaFold shows closer AF2-referenced behaviour than ESMFold in several comparisons, suggesting that its performance improves relative to ESMFold as peptide length increases. This indicates that short-peptide prediction accuracy is not governed only by the overall model architecture, but also by the sequence-length regime in which the model is applied.

A similar structure-dependent pattern emerges from the secondary-structure-stratified analysis. AlphaFold2 remains the most stable reference point, while DMPfold2 and RoseTTAFold2 show stronger separation across most structural classes. However, the relative behaviour of ESMFold and OmegaFold changes with secondary-structure content. In *α*-helix-rich and mixed peptides, OmegaFold shows weaker separation from AlphaFold2 than ESMFold, suggesting that OmegaFold can match or surpass ESMFold-like performance in these structural regimes. This is important because it prevents an overly simple ranking in which ESMFold is always treated as the second-best predictor. Instead, the statistical analysis suggests a more conditional interpretation, ESMFold performs strongly overall, but OmegaFold becomes particularly competitive for longer peptides and for *α*-helical or mixed secondary-structure classes. Our observation supports a regime-dependent view of model performance rather than a single universal ordering of predictors.

This study shows that modern structure predictors can provide useful structural hypotheses for short peptides, but their reliability depends strongly on peptide length, secondary-structure content, disorder, and metric choice. AlphaFold2 and ESMFold are the most consistent overall performers in the experimental benchmark, while RoseTTAFold2 and OmegaFold provide useful complementary predictions and DMPfold2 shows weaker robustness for this peptide regime. The results also show that no single metric fully captures prediction quality: RMSD, native-contact recovery, TM-score, GDT-TS, and confidence calibration each report different aspects of structural agreement. For peptides without experimental structures, such as those in dbAMP3, multi-model consensus provides a rational way to estimate structural robustness. However, predictions for very short, disordered, or environment-sensitive peptides should be interpreted as conformational hypotheses rather than definitive native structures.

## VII. Conclusion

In this study, we systematically evaluated the capabilities and limitations of five prominent deep-learning-based structure prediction models when applied to short peptide sequences. While these predictive architectures have revolutionized the structural modeling of large, globular domains, our findings reveal that their reliability on short sequences is heavily dependent on peptide length, secondary structure composition, and the presence of underlying structural cooperativity.

We found that AlphaFold2 and ESMFold emerged as the most consistently strong predictors across the benchmark, whereas DMPfold2 and RoseTTAFold2 showed larger and more frequent deviations from the experimental reference structures. However, the paired statistical analysis also showed that this ranking was not universal; OmegaFold became increasingly competitive in longer peptides and in *α*-helix-rich or mixed secondary-structure classes. These results suggest that MSA-free predictors can achieve strong short-peptide structure prediction performance, but their relative strengths depend on peptide length and structural class rather than following a single fixed ordering. Across all models, predictions for *α*-helical and mixed-structure peptides were markedly more accurate than those for *β*-sheet-rich and disordered peptides. This underscores a persistent algorithmic limitation, as current predictors remain strongly biased toward the well-defined, compact folds learned from globular proteins.

Furthermore, our evaluation reveals that no single metric fully captures prediction quality. The moderate correlation between predicted confidence (pLDDT) and actual structural accuracy highlights that high confidence scores do not definitively guarantee correct native structures for highly flexible, short sequences. Consequently, we propose a multi-model consensus strategy demonstrated here on the uncharacterized dbAMP3 dataset as a practical filter to estimate structural robustness. Ultimately, predictions for very short, environment-sensitive, or intrinsically disordered peptides must be interpreted as conformational hypotheses rather than definitive native structures.

## Acknowledgements

The authors gratefully acknowledge Dr. Raju Halder, Associate Professor in the Department of Computer Science and Engineering at the Indian Institute of Technology Patna, for providing computational resources and support that enabled portions of the experimental work. The authors also thank Anshuman Jaysingh of Washington State University for insightful discussions and valuable suggestions that contributed to the development of this work.

## Data Availability

All data generated and analyzed during this study are available in the Zenodo repository with DOI: 10.5281/zenodo.21073120. This includes the datasets and predictions ran on the dbAMP 3.0 dataset.

## Appendix A Characteristics of AlphaFold2

**Fig. 23:**
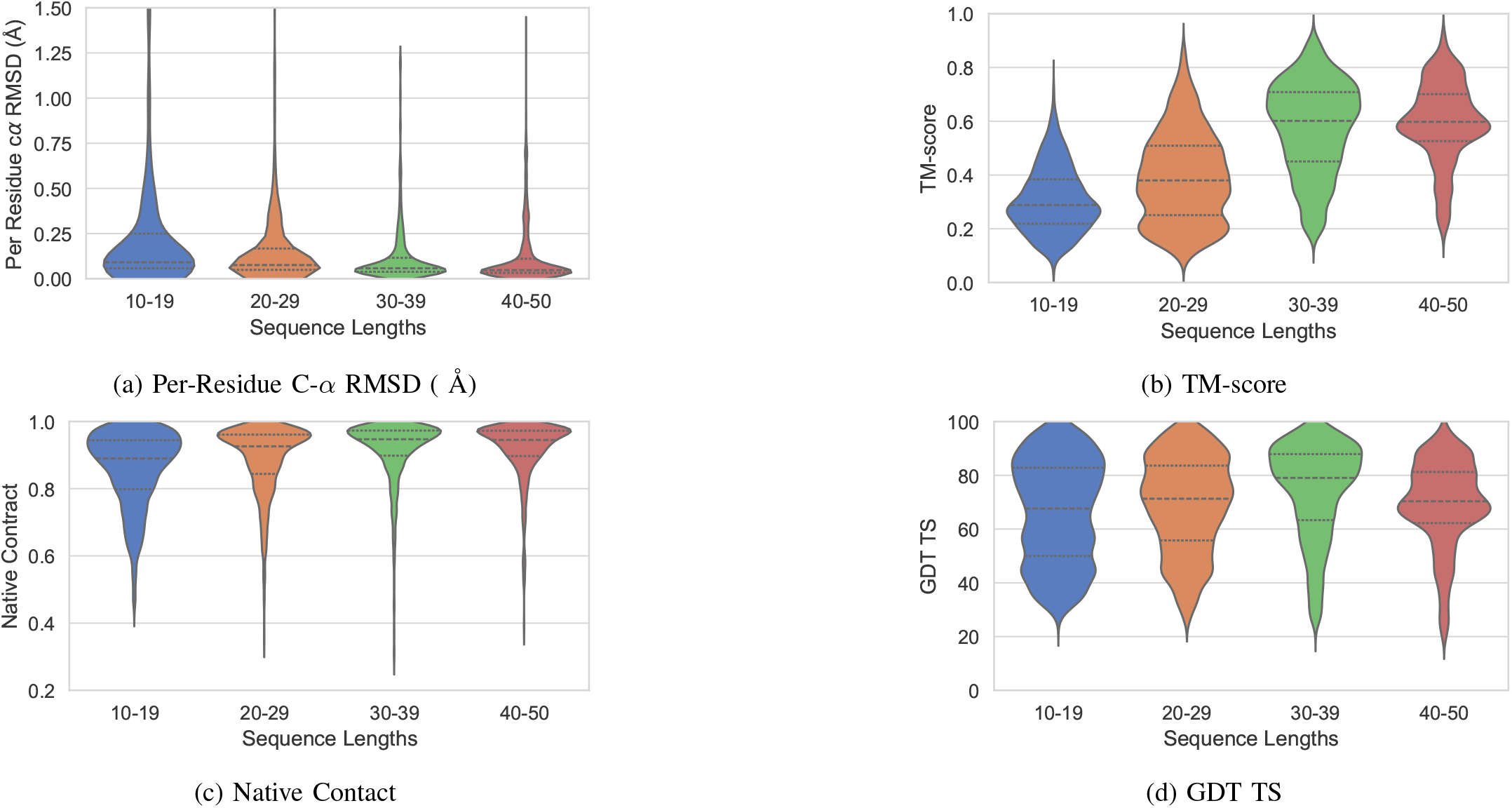
Length-wise. Comparison of Metrics for AlphaFold2

**Fig. 24:**
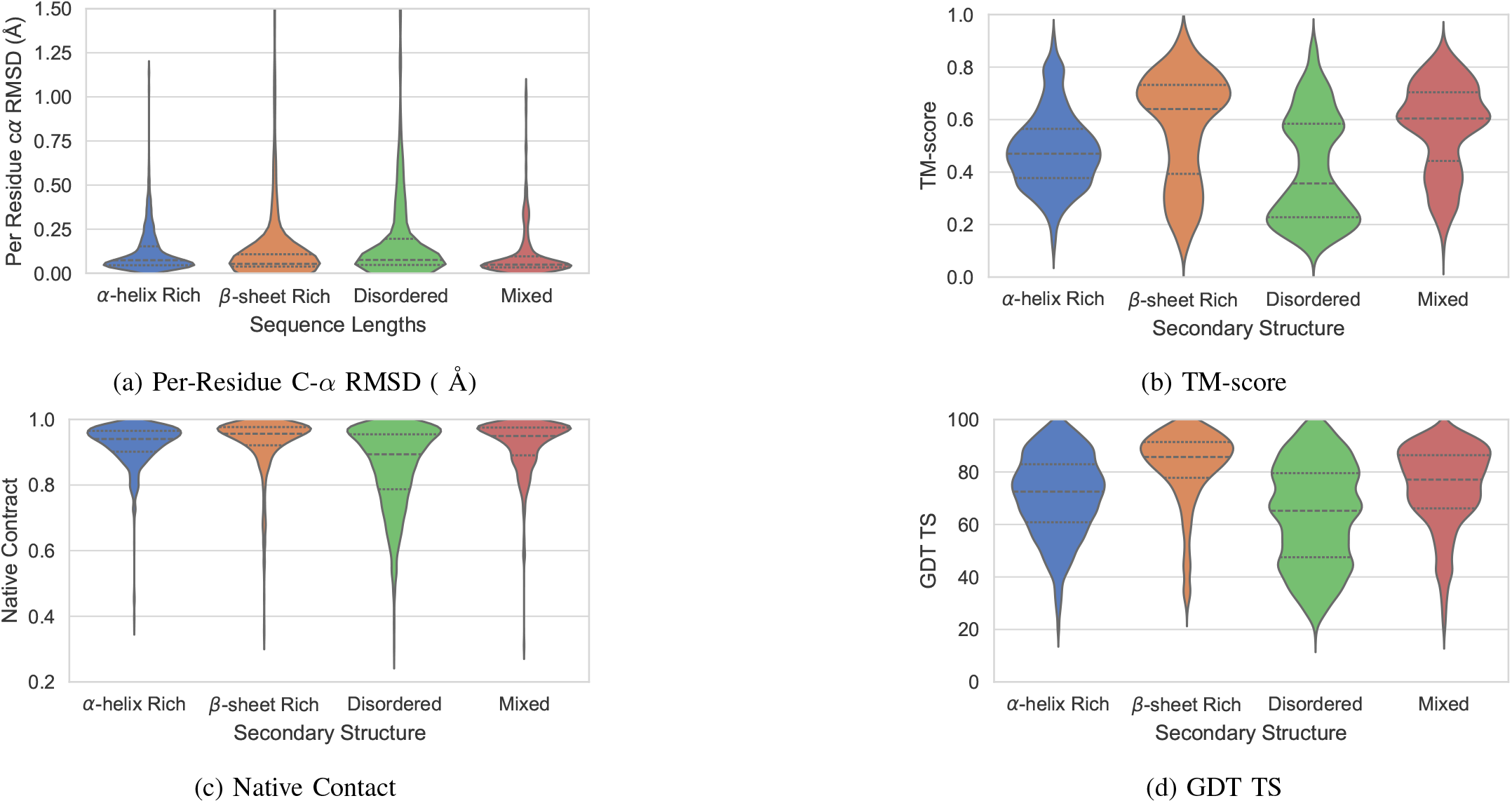
Secondary Structure-wise. Comparison of Metrics for AlphaFold2

**Fig. 25:**
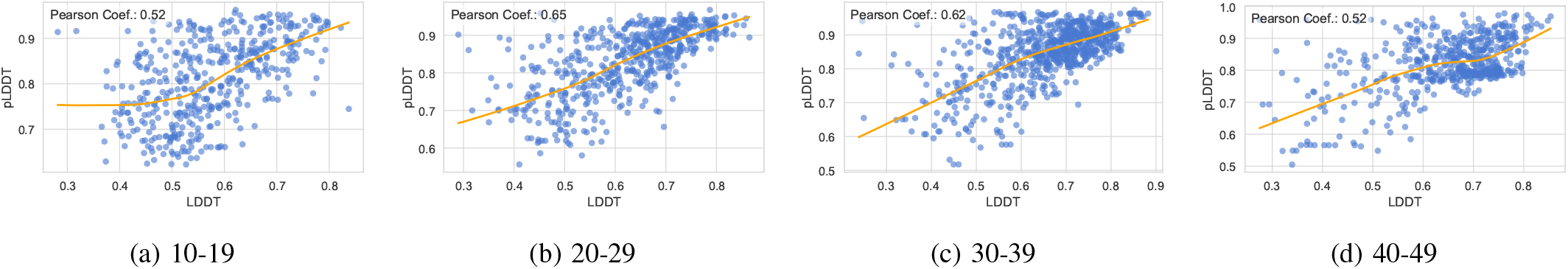
Length-wise. Correlation of LDDT v/s pLDDT for AlphaFold2

**Fig. 26:**
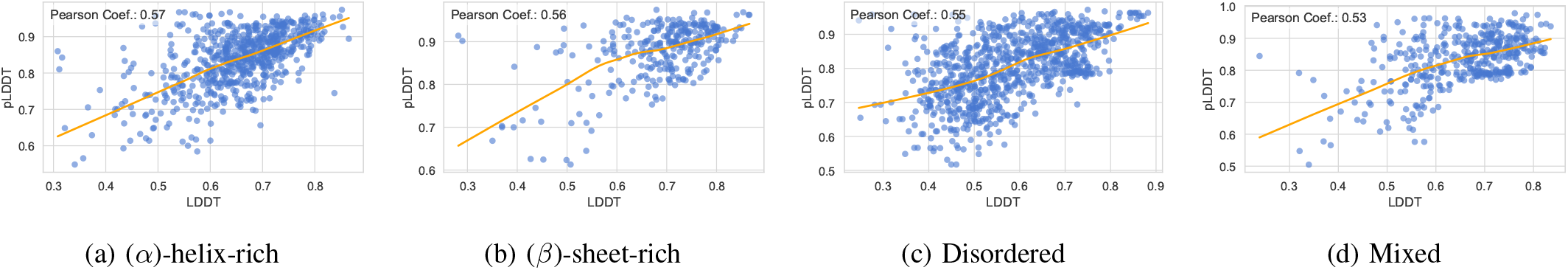
Secondary Structure-wise. Correlation of LDDT v/s pLDDT for AlphaFold2

**TABLE III:** Summary statistics of AlphaFold2 for distributions of Native Contact, TM-score and GDT TS across our benchmark dataset.

| Metric | Statistic | Length-wise |  |  |  |
| --- | --- | --- | --- | --- | --- |
|  |  | 10-19 | 20-29 | 30-39 | 40-49 |
| RMSD | Mean ± SD | 0.30 ± 0.54 | 0.17 ± 0.37 | 0.12 ± 0.16 | 0.12 ± 0.16 |
|  | Median (IQR) | 0.09 (0.06-0.25) | 0.08 (0.05-0.17) | 0.06 (0.04-0.12) | 0.05 (0.03-0.11) |
| **Q** | Mean ± SD | 0.86 ± 0.11 | 0.89 ± 0.10 | 0.91 ± 0.10 | 0.92 ± 0.09 |
|  | Median (IQR) | 0.89 (0.80-0.94) | 0.93 (0.84-0.96) | 0.95 (0.90-0.97) | 0.94 (0.90-0.97) |
| TM-score | Mean ± SD | 0.31 ± 0.12 | 0.39 ± 0.17 | 0.57 ± 0.17 | 0.59 ± 0.15 |
|  | Median (IQR) | 0.29 (0.22-0.38) | 0.38 (0.25-0.51) | 0.60 (0.45-0.71) | 0.60 (0.53-0.70) |
| GDT TS | Mean ± SD | 66.25 ± 19.15 | 69.03 ± 17.75 | 74.17 ± 17.83 | 69.26 ± 16.27 |
|  | Median (IQR) | 67.65 (50.00-82.81) | 71.30 (55.77-83.62) | 79.05 (63.33-87.90) | 70.35 (62.23-81.25) |
| Metric | Statistic | Secondary Structure-wise |  |  |  |
| | | ($\alpha$)-helix-rich | ($\beta$)-sheet-rich | Disordered | Mixed |
| RMSD | Mean ± SD | 0.12 ± 0.12 | 0.19 ± 0.45 | 0.21 ± 0.40 | 0.11 ± 0.15 |
|  | Median (IQR) | 0.07 (0.05-0.15) | 0.05 (0.04-0.11) | 0.08 (0.05-0.20) | 0.05 (0.03-0.10) |
| **Q** | Mean ± SD | 0.92 ± 0.07 | 0.93 ± 0.09 | 0.86 ± 0.12 | 0.92 ± 0.08 |
|  | Median (IQR) | 0.94 (0.90-0.97) | 0.96 (0.92-0.98) | 0.89 (0.79-0.95) | 0.95 (0.89-0.98) |
| TM-score | Mean ± SD | 0.48 ± 0.15 | 0.57 ± 0.21 | 0.41 ± 0.20 | 0.57 ± 0.17 |
|  | Median (IQR) | 0.47 (0.38-0.56) | 0.64 (0.39-0.73) | 0.36 (0.23-0.58) | 0.60 (0.44-0.70) |
| GDT TS | Mean ± SD | 71.15 ± 15.37 | 81.87 ± 14.47 | 63.72 ± 19.14 | 74.29 ± 15.47 |
|  | Median (IQR) | 72.50 (60.83-82.95) | 85.71 (77.78-91.41) | 65.22 (47.50-79.55) | 77.08 (66.13-86.41) |

**TABLE IV:** Summary statistics of AlphaFold2 (without MSA) for distributions of Native Contact, TM-score and GDT TS across our benchmark dataset.

| Metric | Statistic | Length-wise |  |  |  |
| --- | --- | --- | --- | --- | --- |
|  |  | 10-19 | 20-29 | 30-39 | 40-49 |
| **Q** | Mean ± SD | 0.86 ± 0.11 | 0.86 ± 0.13 | 0.86 ± 0.14 | 0.80 ± 0.17 |
|  | Median (IQR) | 0.89 (0.79-0.95) | 0.91 (0.81-0.95) | 0.91 (0.80-0.96) | 0.86 (0.68-0.94) |
| TM-score | Mean ± SD | 0.30 ± 0.12 | 0.37 ± 0.16 | 0.49 ± 0.17 | 0.45 ± 0.16 |
|  | Median (IQR) | 0.28 (0.21-0.37) | 0.37 (0.24-0.48) | 0.49 (0.36-0.63) | 0.45 (0.31-0.57) |
| GDT TS | Mean ± SD | 65.30 ± 19.33 | 66.12 ± 18.55 | 65.43 ± 19.40 | 52.62 ± 18.49 |
|  | Median (IQR) | 66.18 (47.73-82.35) | 68.97 (51.72-80.68) | 67.95 (50.68-81.67) | 53.26 (35.33-66.30) |
| Metric | Statistic | Secondary Structure |  |  |  |
| | | ($\alpha$)-helix-rich | ($\beta$)-sheet-rich | Disordered | Mixed |
| **Q** | Mean ± SD | 0.90 ± 0.10 | 0.87 ± 0.14 | 0.80 ± 0.15 | 0.84 ± 0.14 |
|  | Median (IQR) | 0.93 (0.88-0.96) | 0.94 (0.82-0.96) | 0.83 (0.70-0.92) | 0.89 (0.78-0.95) |
| TM-score | Mean ± SD | 0.45 ± 0.14 | 0.50 ± 0.19 | 0.34 ± 0.16 | 0.45 ± 0.17 |
|  | Median (IQR) | 0.45 (0.37-0.54) | 0.50 (0.33-0.66) | 0.30 (0.22-0.46) | 0.42 (0.32-0.59) |
| GDT TS | Mean ± SD | 67.78 ± 16.71 | 73.76 ± 18.66 | 55.92 ± 19.35 | 60.54 ± 19.93 |
|  | Median (IQR) | 69.74 (56.67-80.26) | 79.35 (63.24-88.33) | 54.17 (39.40-71.30) | 62.84 (42.38-77.27) |

## Appendix B Characteristics of RoseTTAFold2

**Fig. 27:**
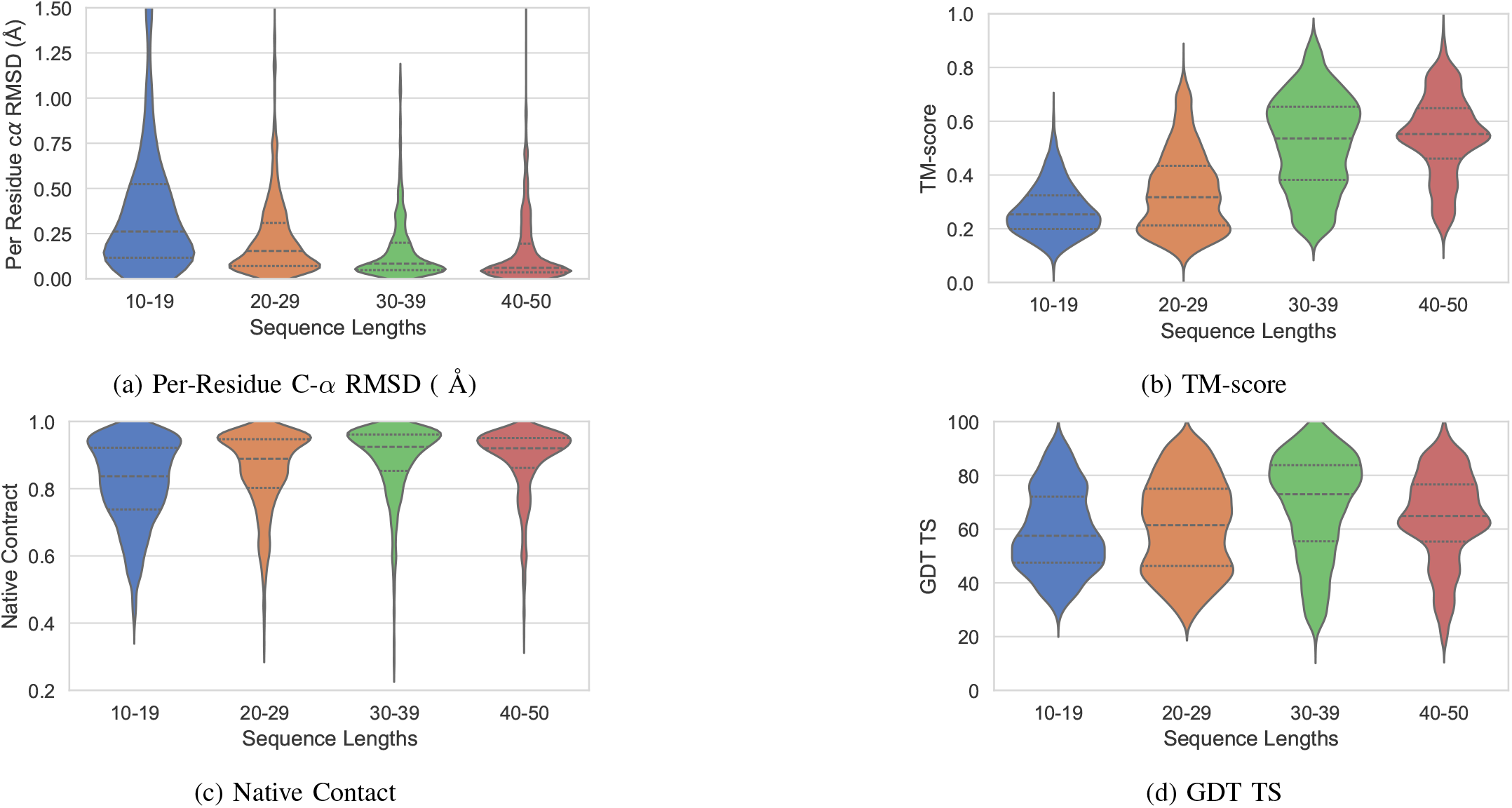
Length-wise. Comparison of Metrics for RoseTTAFold2

**Fig. 28:**
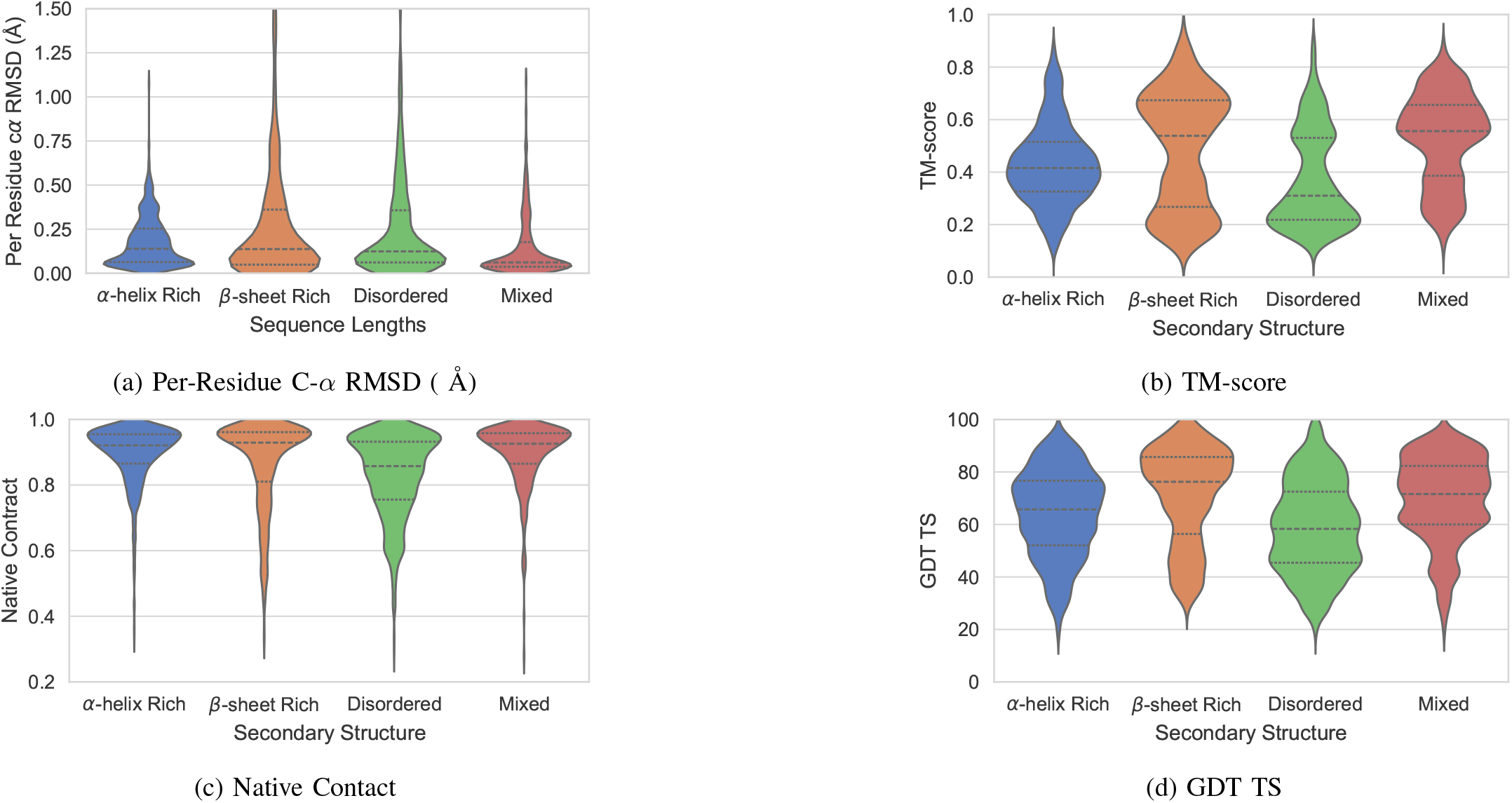
Secondary Structure-wise. Comparison of Metrics for RoseTTAFold2

**Fig. 29:**
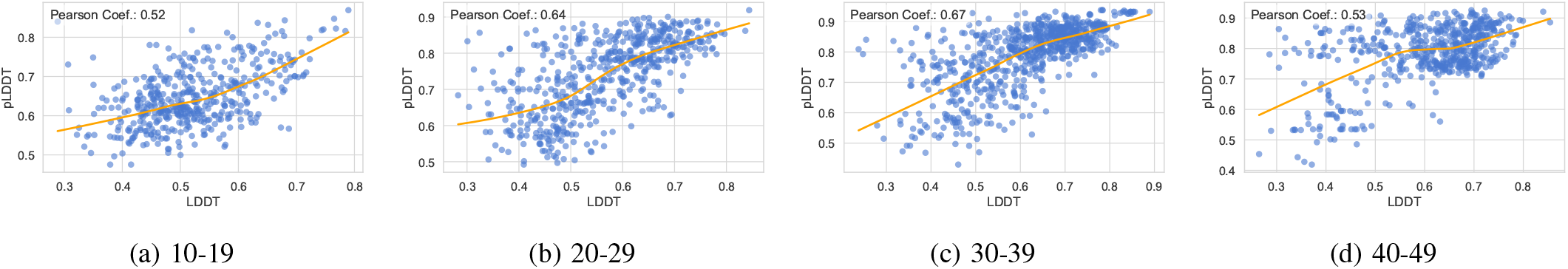
Length-wise. Correlation of LDDT v/s pLDDT for RoseTTAFold2

**Fig. 30:**
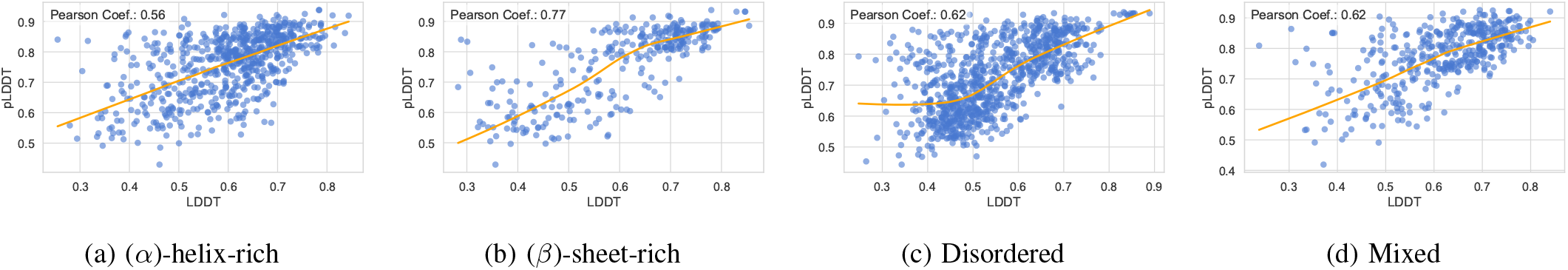
Secondary Structure-wise. Correlation of LDDT v/s pLDDT for RoseTTAFold2

**TABLE V:** Summary statistics of RoseTTAFold2 for distributions of Native Contact, TM-score and GDT TS across our benchmark dataset.

| Metric | Statistic | Length-wise |  |  |  |
| --- | --- | --- | --- | --- | --- |
|  |  | 10-19 | 20-29 | 30-39 | 40-49 |
| RMSD | Mean ± SD | 0.43 ± 0.54 | 0.23 ± 0.23 | 0.16 ± 0.17 | 0.15 ± 0.19 |
|  | Median (IQR) | 0.26 (0.12-0.52) | 0.15 (0.07-0.31) | 0.08 (0.05-0.20) | 0.06 (0.04-0.19) |
| **Q** | Mean ± SD | 0.82 ± 0.12 | 0.86 ± 0.12 | 0.89 ± 0.11 | 0.88 ± 0.10 |
|  | Median (IQR) | 0.84 (0.74-0.92) | 0.89 (0.80-0.95) | 0.92 (0.85-0.96) | 0.92 (0.86-0.95) |
| TM-score | Mean ± SD | 0.27 ± 0.09 | 0.34 ± 0.15 | 0.52 ± 0.17 | 0.55 ± 0.15 |
|  | Median (IQR) | 0.25 (0.20-0.32) | 0.32 (0.21-0.43) | 0.54 (0.38-0.65) | 0.55 (0.46-0.65) |
| GDT TS | Mean ± SD | 59.50 ± 15.37 | 61.09 ± 17.25 | 68.68 ± 18.95 | 63.99 ± 16.87 |
|  | Median (IQR) | 57.50 (47.50-72.06) | 61.46 (46.30-75.00) | 72.97 (55.47-83.78) | 64.88 (55.36-76.60) |
| Metric | Statistic | Secondary Structure-wise |  |  |  |
| | | ($\alpha$)-helix-rich | ($\beta$)-sheet-rich | Disordered | Mixed |
| RMSD | Mean ± SD | 0.18 ± 0.14 | 0.28 ± 0.43 | 0.27 ± 0.35 | 0.14 ± 0.17 |
|  | Median (IQR) | 0.14 (0.06-0.25) | 0.14 (0.05-0.36) | 0.12 (0.06-0.36) | 0.06 (0.04-0.18) |
| **Q** | Mean ± SD | 0.90 ± 0.09 | 0.87 ± 0.14 | 0.83 ± 0.13 | 0.90 ± 0.10 |
|  | Median (IQR) | 0.92 (0.87-0.95) | 0.93 (0.81-0.96) | 0.86 (0.76-0.93) | 0.93 (0.86-0.96) |
| TM-score | Mean ± SD | 0.43 ± 0.15 | 0.49 ± 0.22 | 0.37 ± 0.18 | 0.53 ± 0.17 |
|  | Median (IQR) | 0.42 (0.33-0.52) | 0.54 (0.27-0.67) | 0.31 (0.22-0.53) | 0.56 (0.39-0.66) |
| GDT TS | Mean ± SD | 64.03 ± 16.65 | 70.90 ± 18.34 | 58.90 ± 17.45 | 69.44 ± 16.28 |
|  | Median (IQR) | 65.74 (52.03-76.67) | 76.27 (56.38-85.71) | 58.33 (45.41-72.50) | 71.59 (60.00-82.31) |

**TABLE VI:** Summary statistics of RoseTTAFold2 (without MSA) for distributions of Native Contact, TM-score and GDT TS across our benchmark dataset.

| Metric | Statistic | Length-wise |  |  |  |
| --- | --- | --- | --- | --- | --- |
|  |  | 10-19 | 20-29 | 30-39 | 40-49 |
| **Q** | Mean ± SD | 0.82 ± 0.12 | 0.84 ± 0.12 | 0.82 ± 0.13 | 0.76 ± 0.15 |
|  | Median (IQR) | 0.85 (0.74-0.92) | 0.87 (0.78-0.93) | 0.85 (0.74-0.93) | 0.79 (0.66-0.89) |
| TM-score | Mean ± SD | 0.26 ± 0.09 | 0.31 ± 0.13 | 0.41 ± 0.15 | 0.39 ± 0.13 |
|  | Median (IQR) | 0.25 (0.19-0.31) | 0.29 (0.21-0.40) | 0.39 (0.30-0.50) | 0.36 (0.28-0.49) |
| GDT TS | Mean ± SD | 58.75 ± 14.90 | 57.60 ± 16.14 | 56.60 ± 17.31 | 47.04 ± 15.03 |
|  | Median (IQR) | 57.14 (47.22-69.12) | 57.69 (44.23-70.54) | 54.69 (42.97-69.35) | 44.57 (35.00-58.15) |
| Metric | Statistic | Secondary Structure |  |  |  |
| | | ($\alpha$)-helix-rich | ($\beta$)-sheet-rich | Disordered | Mixed |
| **Q** | Mean ± SD | 0.90 ± 0.09 | 0.78 ± 0.15 | 0.77 ± 0.13 | 0.81 ± 0.13 |
|  | Median (IQR) | 0.92 (0.87-0.95) | 0.80 (0.67-0.91) | 0.78 (0.67-0.87) | 0.83 (0.74-0.91) |
| TM-score | Mean ± SD | 0.41 ± 0.13 | 0.36 ± 0.17 | 0.29 ± 0.11 | 0.39 ± 0.14 |
|  | Median (IQR) | 0.40 (0.32-0.49) | 0.33 (0.22-0.47) | 0.26 (0.21-0.35) | 0.35 (0.28-0.48) |
| GDT TS | Mean ± SD | 62.27 ± 16.13 | 57.30 ± 17.49 | 49.19 ± 14.14 | 54.40 ± 17.08 |
|  | Median (IQR) | 63.71 (51.12-74.36) | 54.76 (43.06-70.59) | 47.50 (38.46-59.02) | 52.27 (40.76-66.96) |

## Appendix C Characteristics of ESMFold

**Fig. 31:**
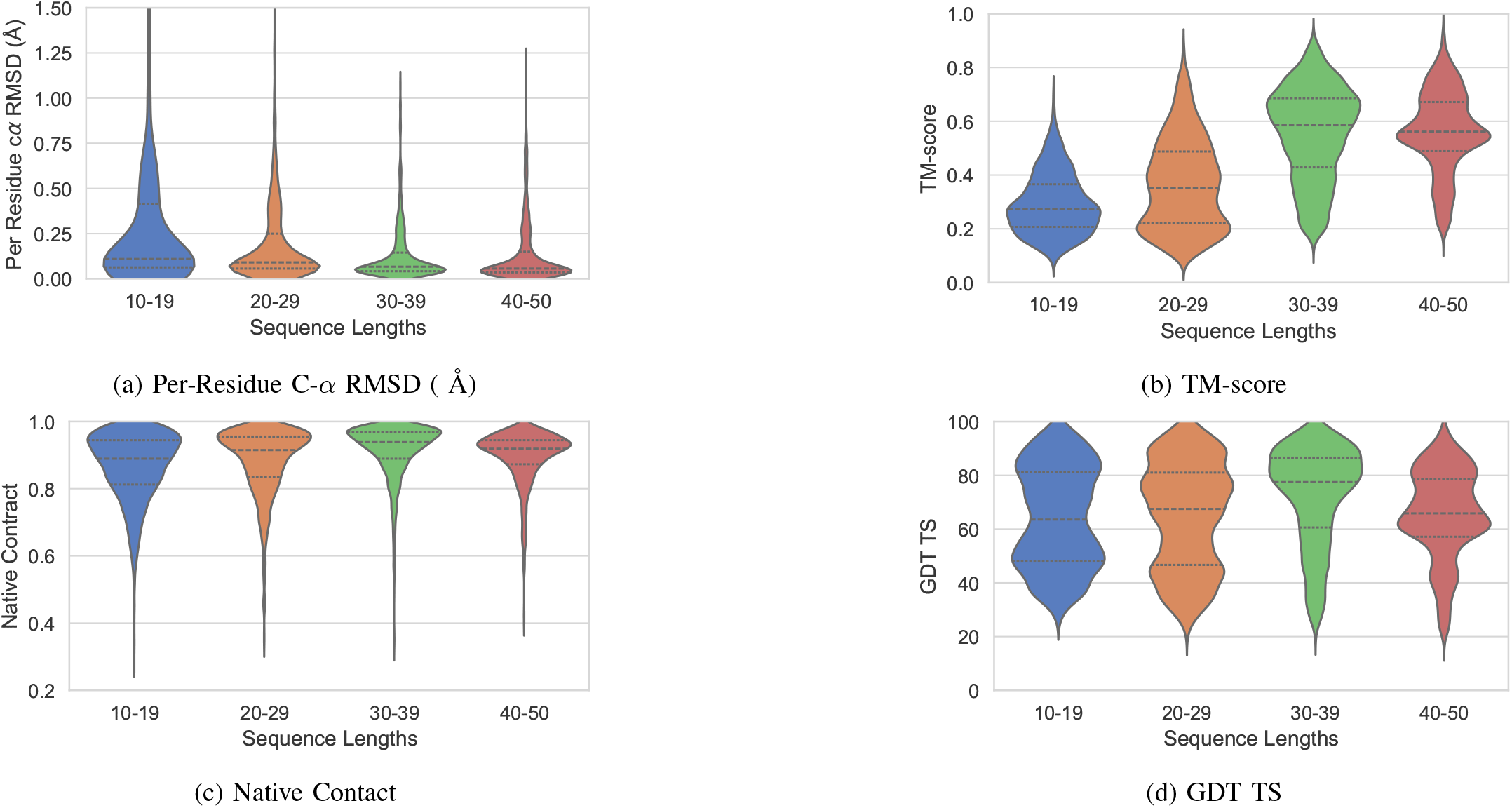
Length-wise. Comparison of Metrics for ESMFold

**Fig. 32:**
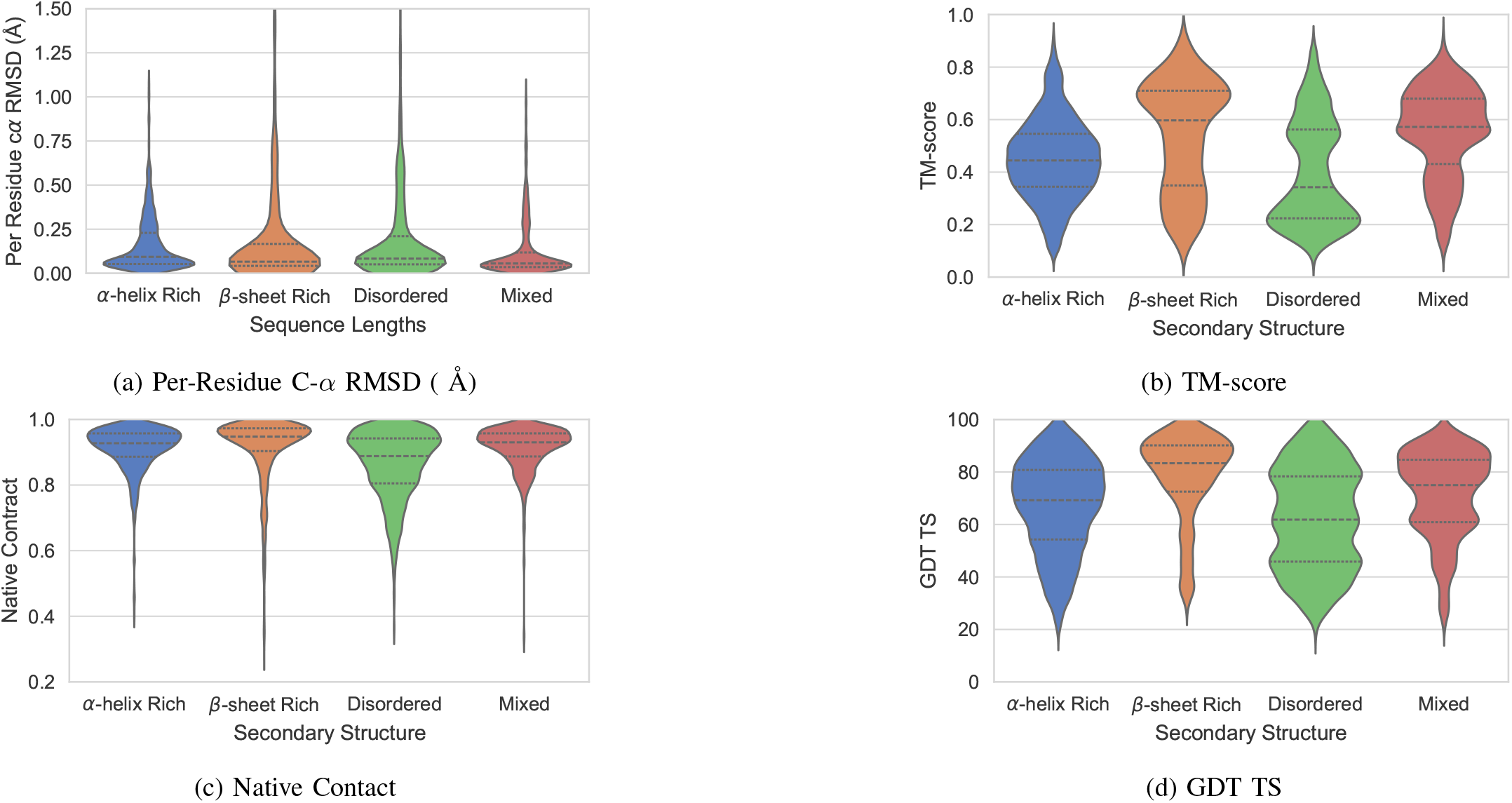
Secondary Structure-wise. Comparison of Metrics for ESMFold

**Fig. 33:**
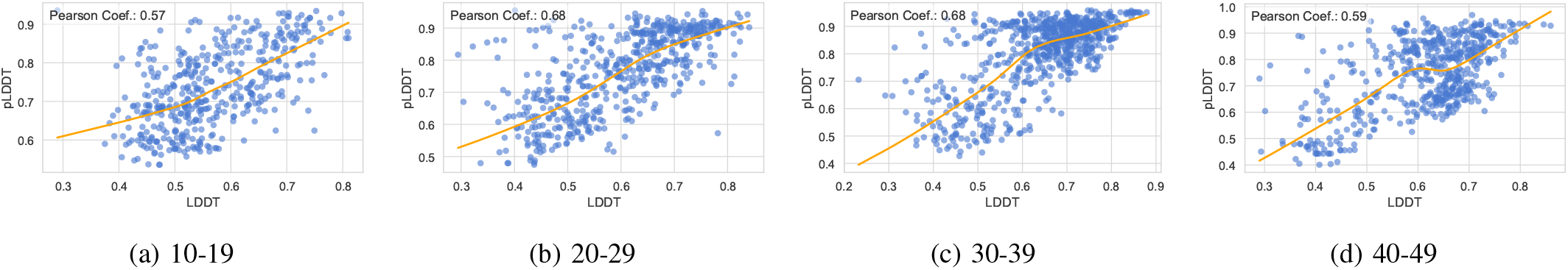
Length-wise. Correlation of LDDT v/s pLDDT for ESMFold

**Fig. 34:**
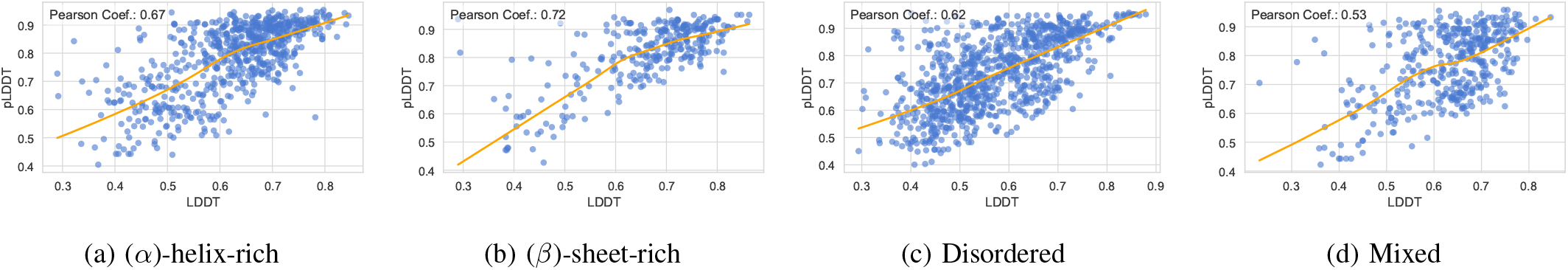
Secondary Structure-wise. Correlation of LDDT v/s pLDDT for ESMFold

**TABLE VII:** Summary statistics of ESMFold for distributions of Native Contact, TM-score and GDT TS across our benchmark dataset.

| Metric | Statistic | Length-wise |  |  |  |
| --- | --- | --- | --- | --- | --- |
|  |  | 10-19 | 20-29 | 30-39 | 40-49 |
| RMSD | Mean ± SD | 0.36 ± 0.60 | 0.21 ± 0.28 | 0.13 ± 0.15 | 0.13 ± 0.17 |
|  | Median (IQR) | 0.11 (0.06-0.42) | 0.09 (0.06-0.25) | 0.07 (0.04-0.14) | 0.06 (0.04-0.15) |
| **Q** | Mean ± SD | 0.87 ± 0.10 | 0.88 ± 0.10 | 0.91 ± 0.08 | 0.90 ± 0.08 |
|  | Median (IQR) | 0.89 (0.81-0.94) | 0.91 (0.83-0.95) | 0.94 (0.89-0.97) | 0.92 (0.87-0.94) |
| TM-score | Mean ± SD | 0.29 ± 0.11 | 0.37 ± 0.17 | 0.55 ± 0.17 | 0.56 ± 0.15 |
|  | Median (IQR) | 0.27 (0.21-0.37) | 0.35 (0.22-0.49) | 0.59 (0.43-0.69) | 0.56 (0.49-0.67) |
| GDT TS | Mean ± SD | 64.27 ± 18.67 | 64.81 ± 19.81 | 72.28 ± 18.44 | 65.44 ± 16.59 |
|  | Median (IQR) | 63.55 (48.21-81.25) | 67.50 (46.67-81.00) | 77.50 (60.59-86.57) | 65.85 (57.14-78.66) |
| Metric | Statistic | Secondary Structure-wise |  |  |  |
| | | ($\alpha$)-helix-rich | ($\beta$)-sheet-rich | Disordered | Mixed |
| RMSD | Mean ± SD | 0.16 ± 0.16 | 0.22 ± 0.46 | 0.23 ± 0.38 | 0.12 ± 0.16 |
|  | Median (IQR) | 0.09 (0.05-0.23) | 0.07 (0.04-0.17) | 0.08 (0.05-0.21) | 0.06 (0.04-0.12) |
| **Q** | Mean ± SD | 0.91 ± 0.07 | 0.91 ± 0.10 | 0.86 ± 0.10 | 0.91 ± 0.07 |
|  | Median (IQR) | 0.93 (0.89-0.96) | 0.95 (0.90-0.97) | 0.89 (0.81-0.94) | 0.93 (0.89-0.96) |
| TM-score | Mean ± SD | 0.45 ± 0.15 | 0.54 ± 0.21 | 0.40 ± 0.20 | 0.55 ± 0.17 |
|  | Median (IQR) | 0.44 (0.34-0.55) | 0.60 (0.35-0.71) | 0.34 (0.22-0.56) | 0.57 (0.43-0.68) |
| GDT TS | Mean ± SD | 66.90 ± 17.65 | 78.26 ± 16.57 | 61.92 ± 19.15 | 71.40 ± 16.45 |
|  | Median (IQR) | 69.23 (54.29-80.77) | 83.33 (72.49-90.15) | 61.84 (45.83-78.33) | 75.00 (60.87-84.66) |

## Appendix D Characteristics of OmegaFold

**Fig. 35:**
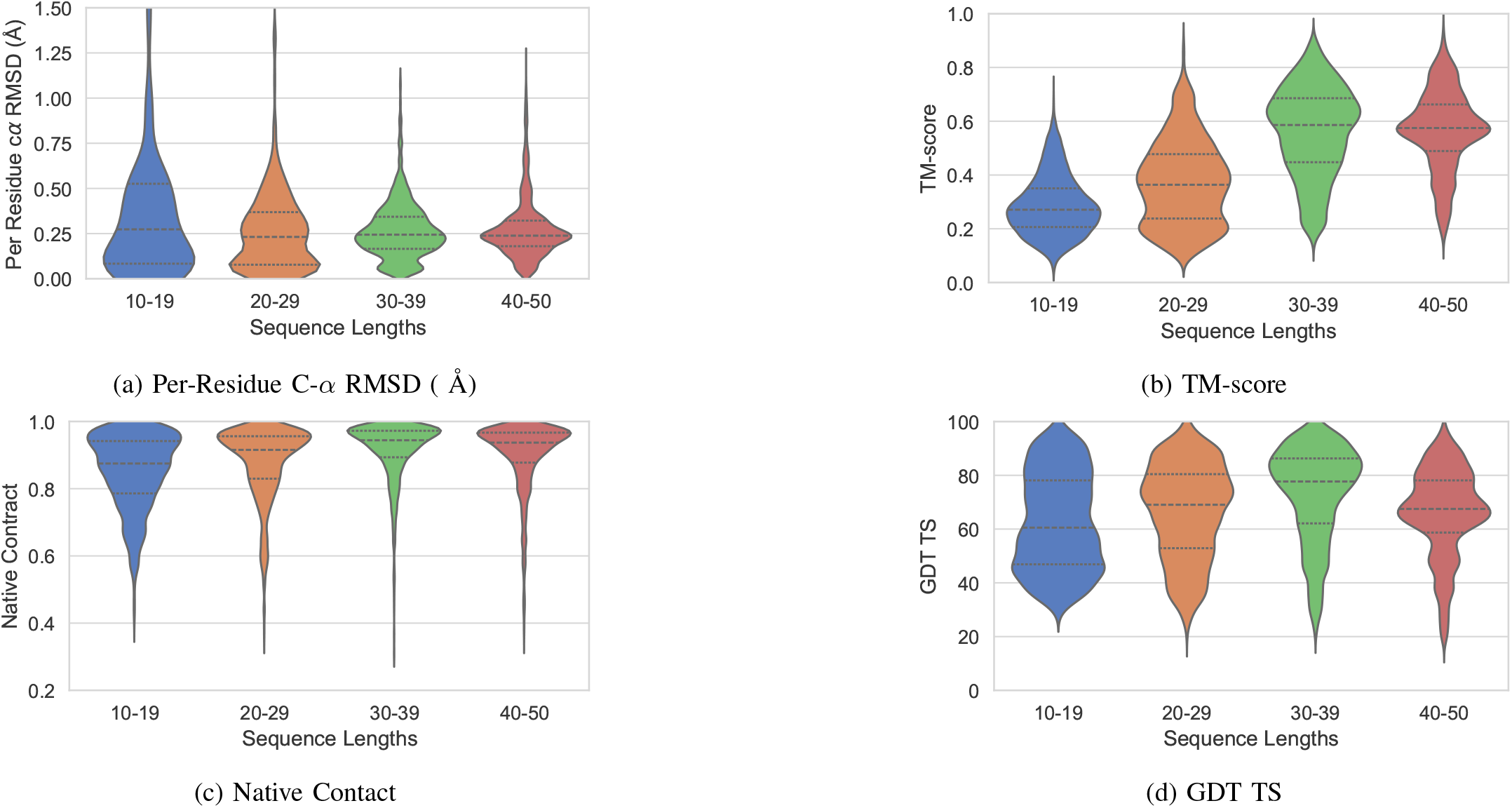
Length-wise. Comparison of Metrics for OmegaFold

**Fig. 36:**
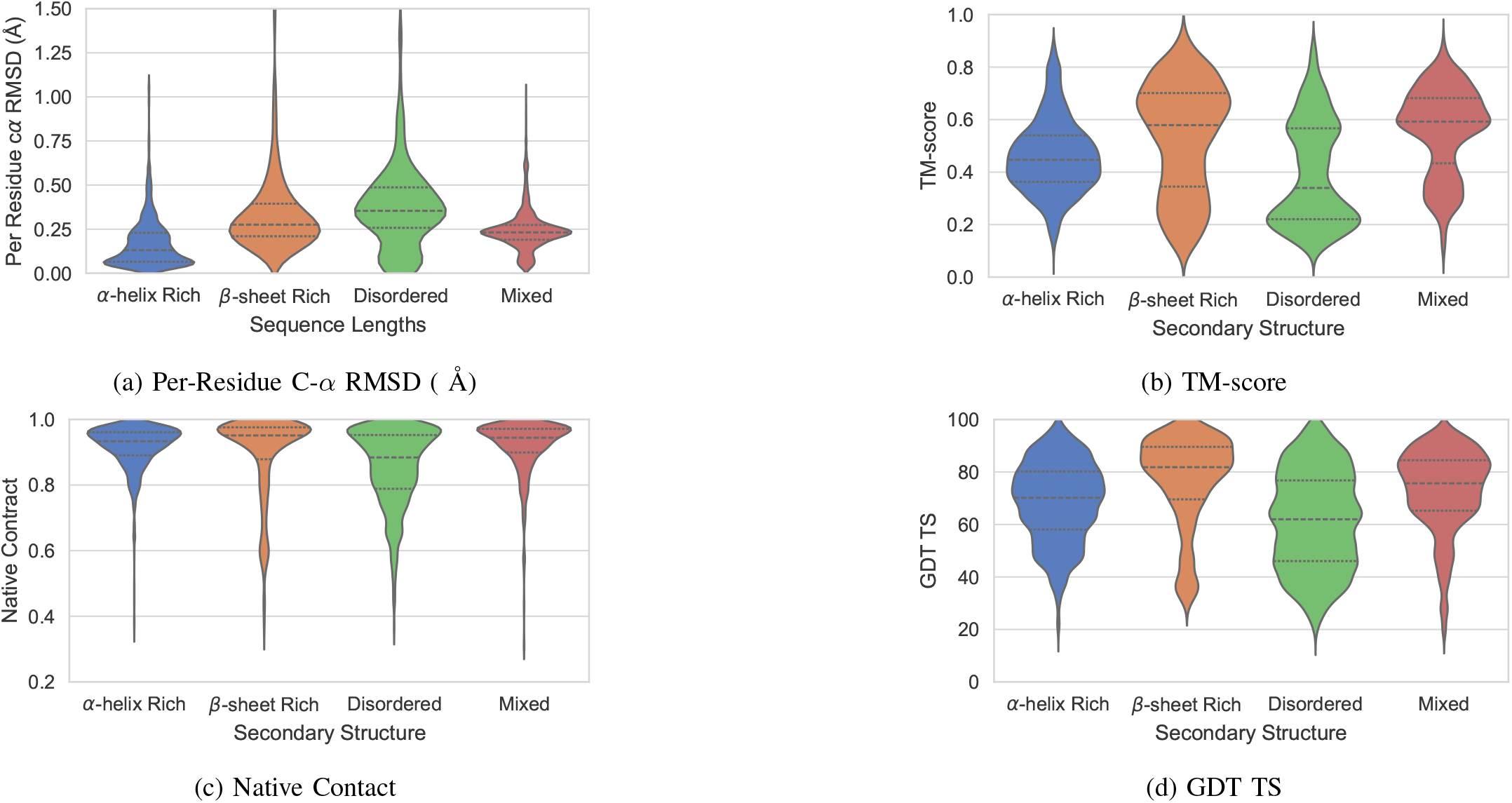
Secondary Structure-wise. Comparison of Metrics for OmegaFold

**TABLE VIII:** Summary statistics of OmegaFold for distributions of Native Contact, TM-score and GDT TS across our benchmark dataset.

| Metric | Statistic | Length-wise |  |  |  |
| --- | --- | --- | --- | --- | --- |
|  |  | 10-19 | 20-29 | 30-39 | 40-49 |
| RMSD | Mean ± SD | 0.46 ± 0.62 | 0.29 ± 0.34 | 0.27 ± 0.16 | 0.27 ± 0.17 |
|  | Median (IQR) | 0.27 (0.08-0.53) | 0.23 (0.08-0.37) | 0.24 (0.17-0.34) | 0.24 (0.18-0.32) |
| **Q** | Mean ± SD | 0.85 ± 0.11 | 0.88 ± 0.11 | 0.91 ± 0.09 | 0.90 ± 0.10 |
|  | Median (IQR) | 0.87 (0.79-0.94) | 0.92 (0.83-0.96) | 0.94 (0.89-0.97) | 0.94 (0.88-0.97) |
| TM-score | Mean ± SD | 0.29 ± 0.11 | 0.37 ± 0.16 | 0.56 ± 0.17 | 0.57 ± 0.15 |
|  | Median (IQR) | 0.27 (0.21-0.35) | 0.36 (0.24-0.48) | 0.59 (0.45-0.69) | 0.57 (0.49-0.66) |
| GDT TS | Mean ± SD | 62.58 ± 18.09 | 66.38 ± 17.68 | 72.93 ± 17.67 | 66.08 ± 16.36 |
|  | Median (IQR) | 60.53 (46.88-78.12) | 69.05 (52.88-80.43) | 77.70 (62.11-86.29) | 67.50 (58.70-78.12) |
| Metric | Statistic | Secondary Structure-wise |  |  |  |
| | | ($\alpha$)-helix-rich | ($\beta$)-sheet-rich | Disordered | Mixed |
| RMSD | Mean ± SD | 0.17 ± 0.14 | 0.39 ± 0.41 | 0.43 ± 0.43 | 0.24 ± 0.12 |
|  | Median (IQR) | 0.13 (0.07-0.23) | 0.28 (0.21-0.39) | 0.35 (0.26-0.49) | 0.23 (0.19-0.27) |
| **Q** | Mean ± SD | 0.92 ± 0.07 | 0.89 ± 0.13 | 0.86 ± 0.12 | 0.92 ± 0.08 |
|  | Median (IQR) | 0.93 (0.89-0.96) | 0.95 (0.88-0.98) | 0.88 (0.79-0.95) | 0.94 (0.90-0.97) |
| TM-score | Mean ± SD | 0.46 ± 0.14 | 0.53 ± 0.21 | 0.39 ± 0.20 | 0.56 ± 0.17 |
|  | Median (IQR) | 0.45 (0.36-0.54) | 0.58 (0.34-0.70) | 0.34 (0.22-0.57) | 0.59 (0.43-0.68) |
| GDT TS | Mean ± SD | 68.63 ± 15.18 | 76.70 ± 17.55 | 61.50 ± 18.62 | 72.97 ± 15.53 |
|  | Median (IQR) | 70.16 (58.04-80.19) | 81.82 (69.56-89.58) | 61.96 (46.05-76.79) | 75.68 (65.22-84.46) |

## Appendix E Characteristics of DMPfold2

**Fig. 37:**
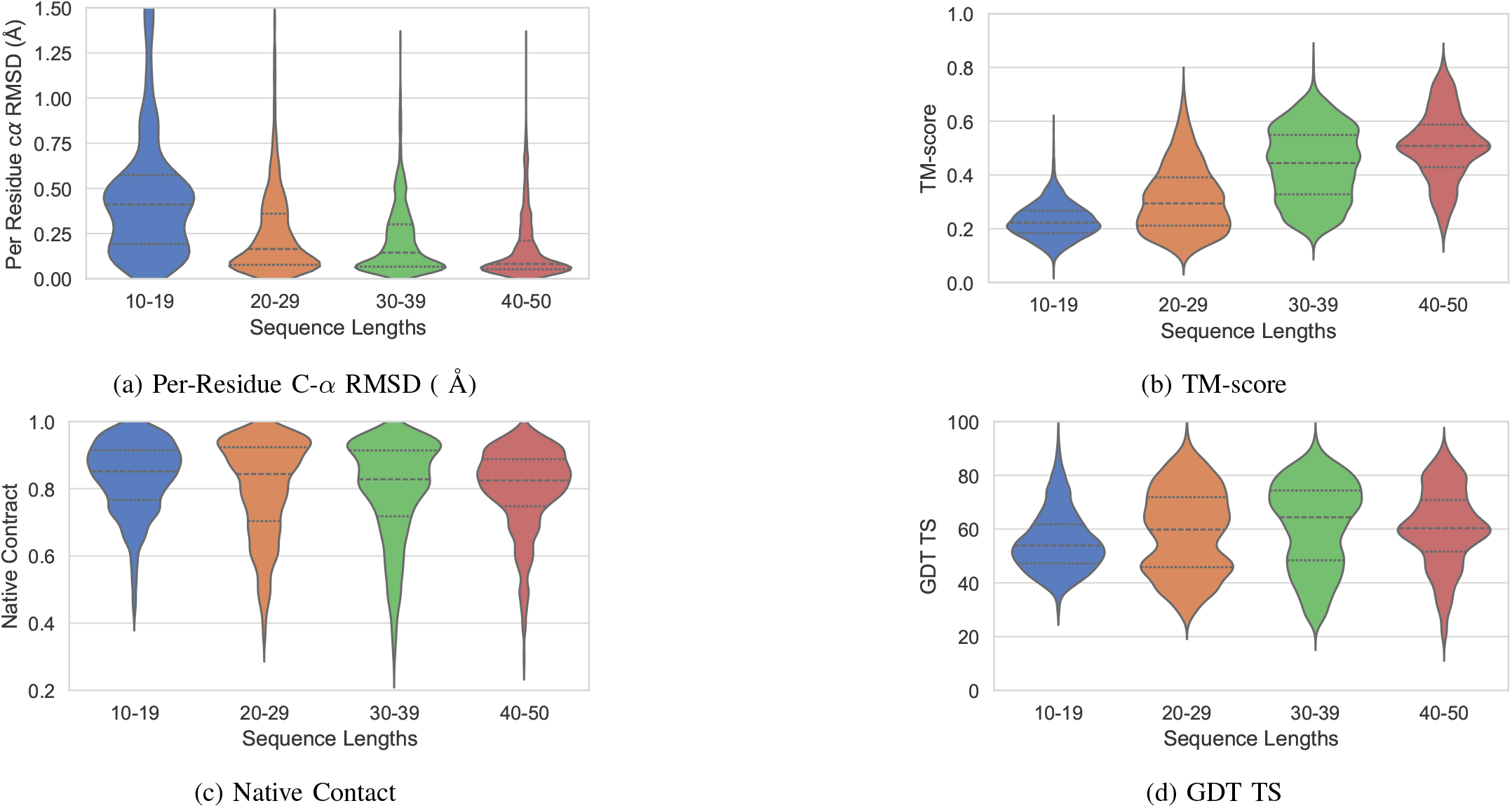
Length-wise. Comparison of Metrics for DMPfold2

**Fig. 38:**
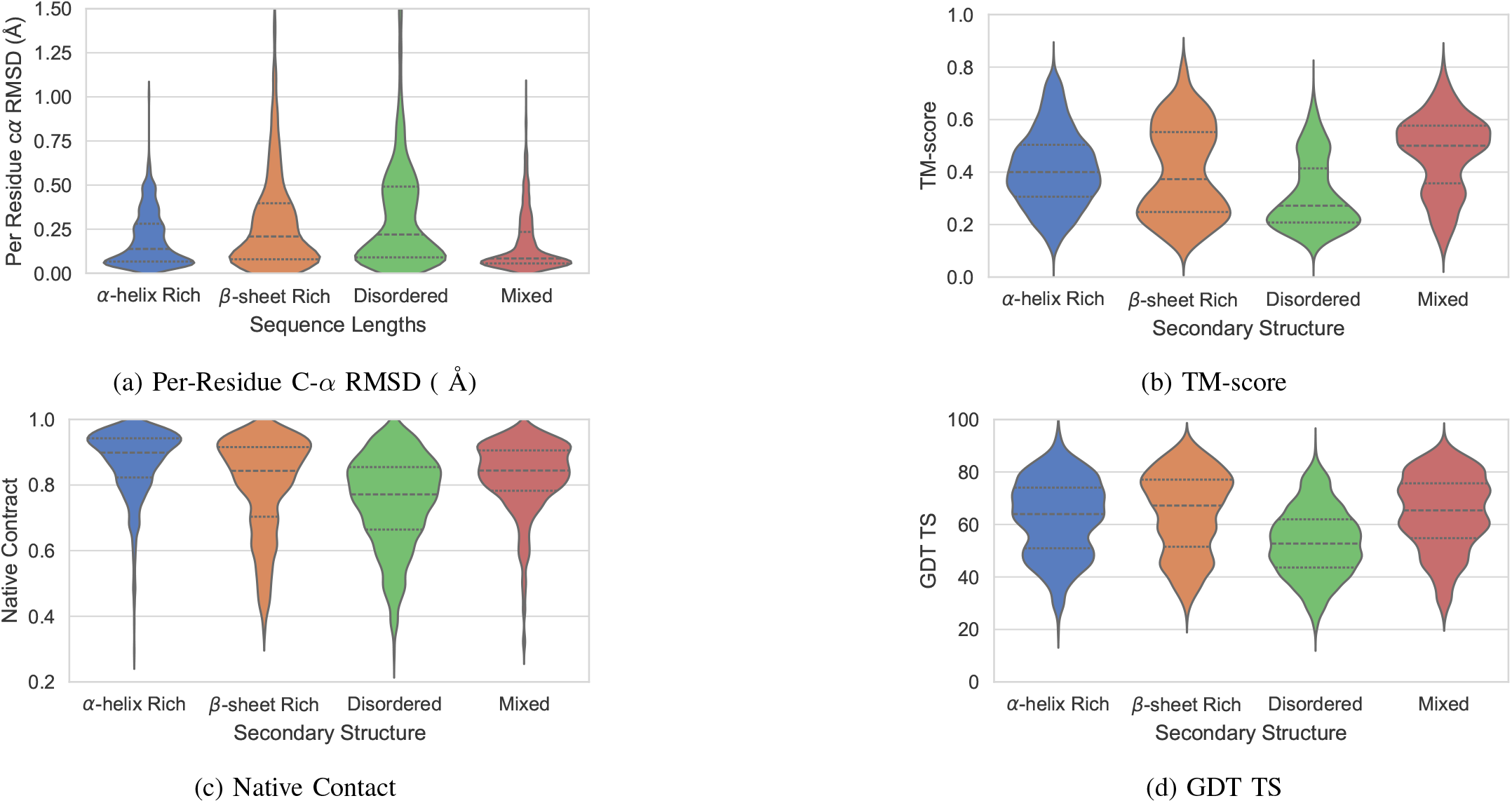
Secondary Structure-wise. Comparison of Metrics for DMPfold2

**Fig. 39:**
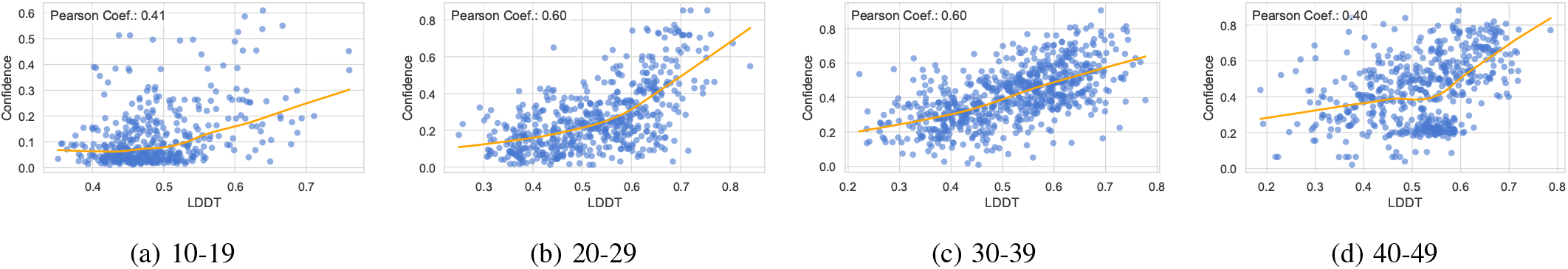
Length-wise. Correlation of LDDT v/s Confidence Scores for DMPfold2

**Fig. 40:**
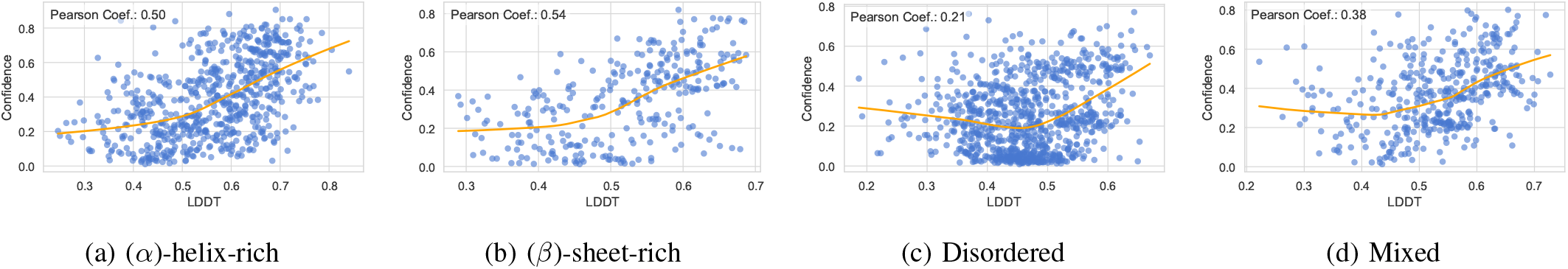
Secondary Structure-wise. Correlation of LDDT v/s Confidence for DMPfold2

**TABLE IX:** Summary statistics of DMPfold2 for distributions of Native Contact, TM-score and GDT TS across our benchmark dataset.

| Metric | Statistic | Length-wise |  |  |  |
| --- | --- | --- | --- | --- | --- |
|  |  | 10-19 | 20-29 | 30-39 | 40-49 |
| RMSD | Mean ± SD | 0.49 ± 0.42 | 0.25 ± 0.25 | 0.21 ± 0.19 | 0.16 ± 0.17 |
|  | Median (IQR) | 0.41 (0.19-0.57) | 0.17 (0.08-0.36) | 0.14 (0.07-0.30) | 0.08 (0.05-0.21) |
| **Q** | Mean ± SD | 0.83 ± 0.11 | 0.80 ± 0.14 | 0.79 ± 0.15 | 0.80 ± 0.12 |
|  | Median (IQR) | 0.85 (0.77-0.91) | 0.84 (0.70-0.92) | 0.83 (0.72-0.91) | 0.82 (0.75-0.89) |
| TM-score | Mean ± SD | 0.23 ± 0.06 | 0.31 ± 0.13 | 0.44 ± 0.13 | 0.51 ± 0.13 |
|  | Median (IQR) | 0.22 (0.18-0.27) | 0.29 (0.21-0.39) | 0.44 (0.33-0.55) | 0.51 (0.43-0.59) |
| GDT TS | Mean ± SD | 55.38 ± 11.37 | 59.02 ± 15.69 | 61.32 ± 16.23 | 60.37 ± 14.53 |
|  | Median (IQR) | 53.85 (47.22-61.76) | 59.82 (45.83-71.88) | 64.39 (48.39-74.34) | 60.33 (51.63-70.83) |
| Metric | Statistic | Secondary Structure-wise |  |  |  |
| | | ($\alpha$)-helix-rich | ($\beta$)-sheet-rich | Disordered | Mixed |
| RMSD | Mean ± SD | 0.19 ± 0.15 | 0.31 ± 0.35 | 0.34 ± 0.33 | 0.16 ± 0.16 |
|  | Median (IQR) | 0.14 (0.07-0.28) | 0.21 (0.08-0.40) | 0.22 (0.09-0.49) | 0.08 (0.06-0.23) |
| **Q** | Mean ± SD | 0.87 ± 0.10 | 0.80 ± 0.15 | 0.75 ± 0.14 | 0.83 ± 0.11 |
|  | Median (IQR) | 0.90 (0.82-0.94) | 0.84 (0.70-0.92) | 0.77 (0.66-0.85) | 0.84 (0.78-0.91) |
| TM-score | Mean ± SD | 0.41 ± 0.14 | 0.40 ± 0.18 | 0.31 ± 0.14 | 0.47 ± 0.15 |
|  | Median (IQR) | 0.40 (0.31-0.50) | 0.37 (0.25-0.55) | 0.27 (0.21-0.41) | 0.50 (0.36-0.58) |
| GDT TS | Mean ± SD | 62.61 ± 14.56 | 64.36 ± 15.49 | 53.04 ± 13.11 | 64.27 ± 14.34 |
|  | Median (IQR) | 63.95 (50.93-74.04) | 67.19 (51.52-77.08) | 52.72 (43.59-61.96) | 65.36 (54.79-75.68) |

**TABLE X:** Summary statistics of DMPfold2 (without MSA) for distributions of Native Contact, TM-score and GDT TS across our benchmark dataset.

| Metric | Statistic | Length-wise |  |  |  |
| --- | --- | --- | --- | --- | --- |
|  |  | 10-19 | 20-29 | 30-39 | 40-49 |
| **Q** | Mean ± SD | 0.83 ± 0.11 | 0.77 ± 0.16 | 0.70 ± 0.15 | 0.64 ± 0.16 |
|  | Median (IQR) | 0.85 (0.76-0.91) | 0.80 (0.64-0.90) | 0.70 (0.58-0.82) | 0.64 (0.52-0.75) |
| TM-score | Mean ± SD | 0.22 ± 0.06 | 0.29 ± 0.12 | 0.36 ± 0.10 | 0.37 ± 0.11 |
|  | Median (IQR) | 0.22 (0.18-0.26) | 0.28 (0.20-0.37) | 0.34 (0.29-0.43) | 0.35 (0.29-0.43) |
| GDT TS | Mean ± SD | 53.84 ± 11.04 | 55.23 ± 15.28 | 50.42 ± 13.77 | 43.93 ± 13.04 |
|  | Median (IQR) | 52.27 (46.15-60.00) | 55.00 (42.50-66.67) | 48.03 (39.86-60.53) | 41.30 (34.38-51.53) |
| Metric | Statistic | Secondary Structure |  |  |  |
| | | ($\alpha$)-helix-rich | ($\beta$)-sheet-rich | Disordered | Mixed |
| **Q** | Mean ± SD | 0.84 ± 0.12 | 0.70 ± 0.16 | 0.67 ± 0.16 | 0.70 ± 0.14 |
|  | Median (IQR) | 0.87 (0.79-0.93) | 0.71 (0.58-0.84) | 0.67 (0.55-0.79) | 0.70 (0.60-0.80) |
| TM-score | Mean ± SD | 0.38 ± 0.13 | 0.32 ± 0.12 | 0.26 ± 0.08 | 0.35 ± 0.09 |
|  | Median (IQR) | 0.38 (0.29-0.47) | 0.30 (0.23-0.42) | 0.26 (0.20-0.31) | 0.34 (0.29-0.41) |
| GDT TS | Mean ± SD | 58.10 ± 14.96 | 53.33 ± 14.52 | 45.36 ± 11.09 | 48.87 ± 12.92 |
|  | Median (IQR) | 58.78 (45.83-69.19) | 52.21 (40.95-64.06) | 44.32 (37.07-52.94) | 47.06 (39.06-57.24) |

## Appendix F DBAMP3 Statistics

**TABLE XI:**
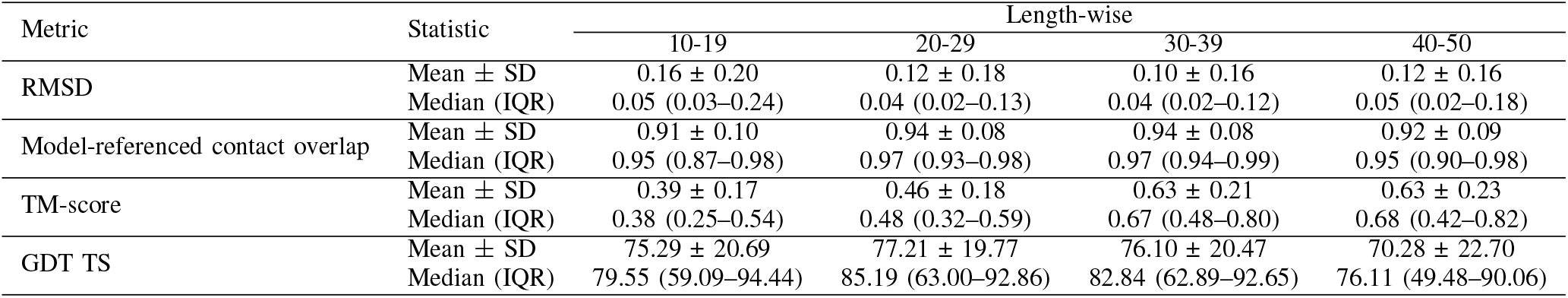
Summary statistics of AlphaFold2 vs. ESMFold for distributions of Native Contact, TM-score and GDT TS across dbAMP3 dataset.

| Metric | Statistic | Length-wise |  |  |  |
| --- | --- | --- | --- | --- | --- |
|  |  | 10-19 | 20-29 | 30-39 | 40-50 |
| RMSD | Mean ± SD | 0.16 ± 0.20 | 0.12 ± 0.18 | 0.10 ± 0.16 | 0.12 ± 0.16 |
|  | Median (IQR) | 0.05 (0.03–0.24) | 0.04 (0.02–0.13) | 0.04 (0.02–0.12) | 0.05 (0.02–0.18) |
| Model-referenced contact overlap | Mean ± SD | 0.91 ± 0.10 | 0.94 ± 0.08 | 0.94 ± 0.08 | 0.92 ± 0.09 |
|  | Median (IQR) | 0.95 (0.87–0.98) | 0.97 (0.93–0.98) | 0.97 (0.94–0.99) | 0.95 (0.90–0.98) |
| TM-score | Mean ± SD | 0.39 ± 0.17 | 0.46 ± 0.18 | 0.63 ± 0.21 | 0.63 ± 0.23 |
|  | Median (IQR) | 0.38 (0.25–0.54) | 0.48 (0.32–0.59) | 0.67 (0.48–0.80) | 0.68 (0.42–0.82) |
| GDT TS | Mean ± SD | 75.29 ± 20.69 | 77.21 ± 19.77 | 76.10 ± 20.47 | 70.28 ± 22.70 |
|  | Median (IQR) | 79.55 (59.09–94.44) | 85.19 (63.00–92.86) | 82.84 (62.89–92.65) | 76.11 (49.48–90.06) |

**TABLE XII:** Summary statistics of AlphaFold2 vs. RoseTTAFold2 for distributions of Native Contact, TM-score and GDT TS across dbAMP3 dataset.

| Metric | Statistic | Length-wise |  |  |  |
| --- | --- | --- | --- | --- | --- |
|  |  | 10-19 | 20-29 | 30-39 | 40-50 |
| RMSD | Mean ± SD | 0.21 ± 0.18 | 0.18 ± 0.15 | 0.13 ± 0.15 | 0.12 ± 0.14 |
|  | Median (IQR) | 0.16 (0.06–0.34) | 0.15 (0.05–0.28) | 0.08 (0.03–0.19) | 0.08 (0.02–0.19) |
| Model-referenced contact overlap | Mean ± SD | 0.85 ± 0.14 | 0.89 ± 0.12 | 0.92 ± 0.10 | 0.91 ± 0.10 |
|  | Median (IQR) | 0.90 (0.78–0.96) | 0.93 (0.84–0.97) | 0.96 (0.91–0.98) | 0.95 (0.88–0.97) |
| TM-score | Mean ± SD | 0.35 ± 0.14 | 0.39 ± 0.17 | 0.57 ± 0.20 | 0.61 ± 0.23 |
|  | Median (IQR) | 0.32 (0.24–0.45) | 0.37 (0.24–0.51) | 0.57 (0.41–0.74) | 0.65 (0.42–0.81) |
| GDT TS | Mean ± SD | 69.06 ± 18.60 | 67.22 ± 19.25 | 70.42 ± 20.16 | 68.98 ± 23.28 |
|  | Median (IQR) | 67.86 (53.85–86.54) | 67.50 (52.50–84.26) | 72.02 (56.43–88.24) | 74.42 (51.60–89.38) |

**TABLE XIII:** Summary statistics of RoseTTAFold2 vs. ESMFold for distributions of Native Contact, TM-score and GDT TS across dbAMP3 dataset.

| Metric | Statistic | Length-wise |  |  |  |
| --- | --- | --- | --- | --- | --- |
|  |  | 10-19 | 20-29 | 30-39 | 40-50 |
| RMSD | Mean ± SD | 0.15 ± 0.22 | 0.15 ± 0.21 | 0.14 ± 0.19 | 0.13 ± 0.19 |
|  | Median (IQR) | 0.06 (0.04–0.14) | 0.06 (0.03–0.20) | 0.05 (0.02–0.19) | 0.05 (0.02–0.19) |
| Model-referenced contact overlap | Mean ± SD | 0.86 ± 0.12 | 0.88 ± 0.12 | 0.90 ± 0.11 | 0.88 ± 0.11 |
|  | Median (IQR) | 0.90 (0.80–0.96) | 0.92 (0.82–0.97) | 0.94 (0.87–0.98) | 0.92 (0.83–0.97) |
| TM-score | Mean ± SD | 0.33 ± 0.13 | 0.38 ± 0.19 | 0.56 ± 0.21 | 0.59 ± 0.23 |
|  | Median (IQR) | 0.31 (0.23–0.42) | 0.36 (0.22–0.52) | 0.56 (0.39–0.75) | 0.62 (0.36–0.79) |
| GDT TS | Mean ± SD | 67.78 ± 18.17 | 66.88 ± 19.46 | 69.37 ± 21.07 | 66.53 ± 23.58 |
|  | Median (IQR) | 65.00 (52.63–84.72) | 66.67 (50.89–84.48) | 71.62 (52.34–88.67) | 71.43 (44.38–87.50) |

## Appendix G Additional Wilcoxon-P Test Heatmaps

**Fig. 41:**
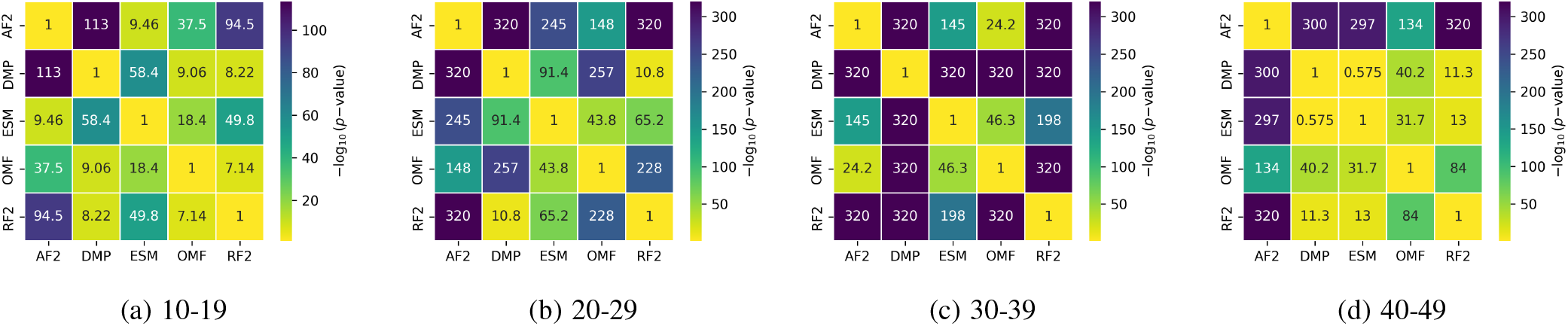
Wilcoxon signed-rank test significance heatmap for pairwise model comparisons using RMSD across peptide length bins.

**Fig. 42:**
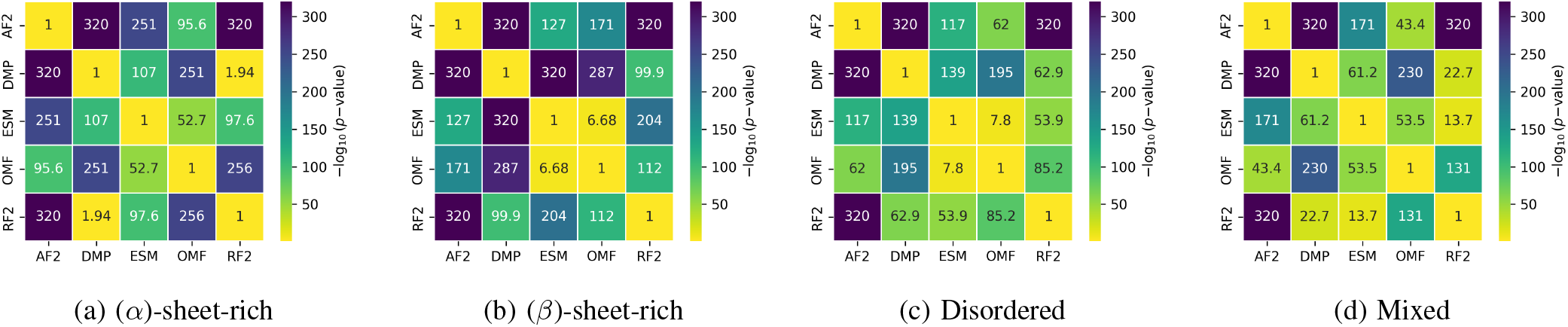
Wilcoxon signed-rank test significance heatmap for pairwise model comparisons using RMSD across secondary-structure classes.

**Fig. 43:**
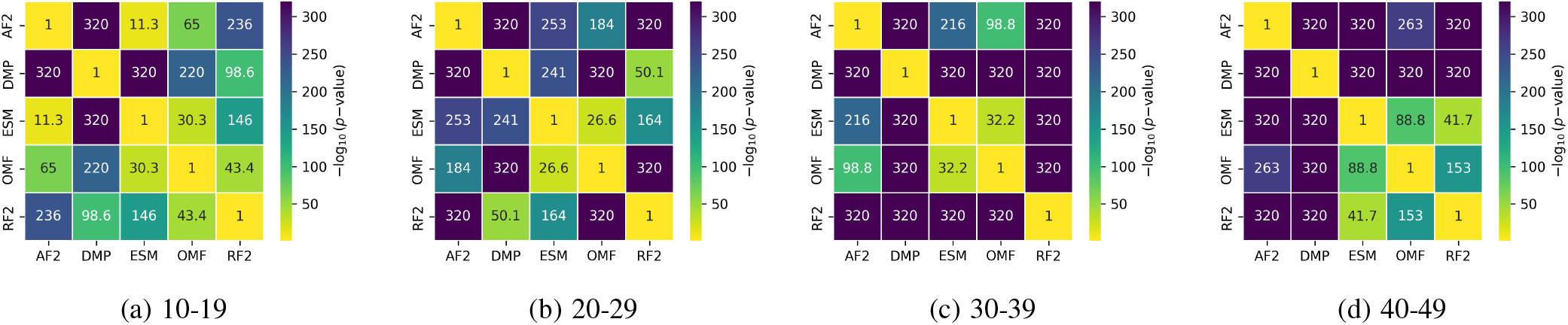
Wilcoxon signed-rank test significance heatmap for pairwise model comparisons using GDT-TS across peptide length bins.

**Fig. 44:**
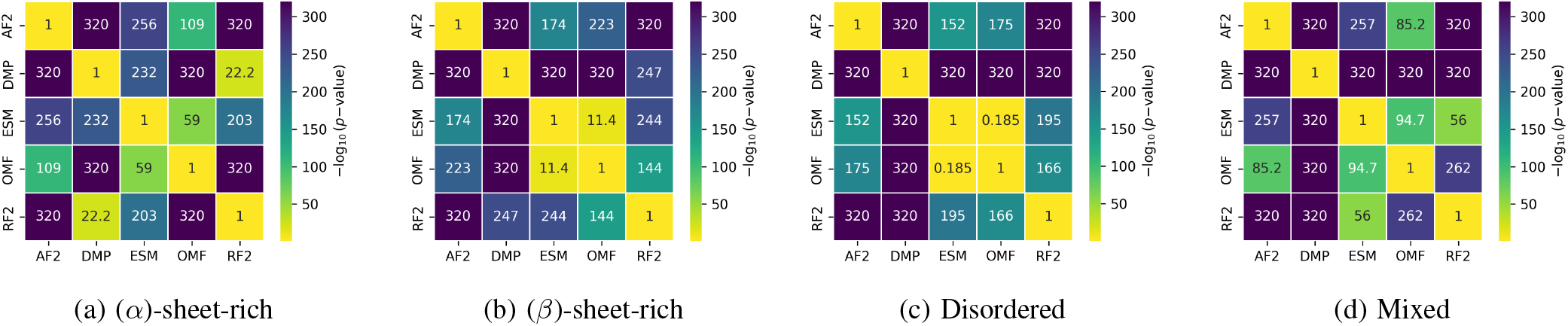
Wilcoxon signed-rank test significance heatmap for pairwise model comparisons using GDT-TS across secondary-structure classes.

## Appendix H PDB Dataset Statistics

**Fig. 45:**
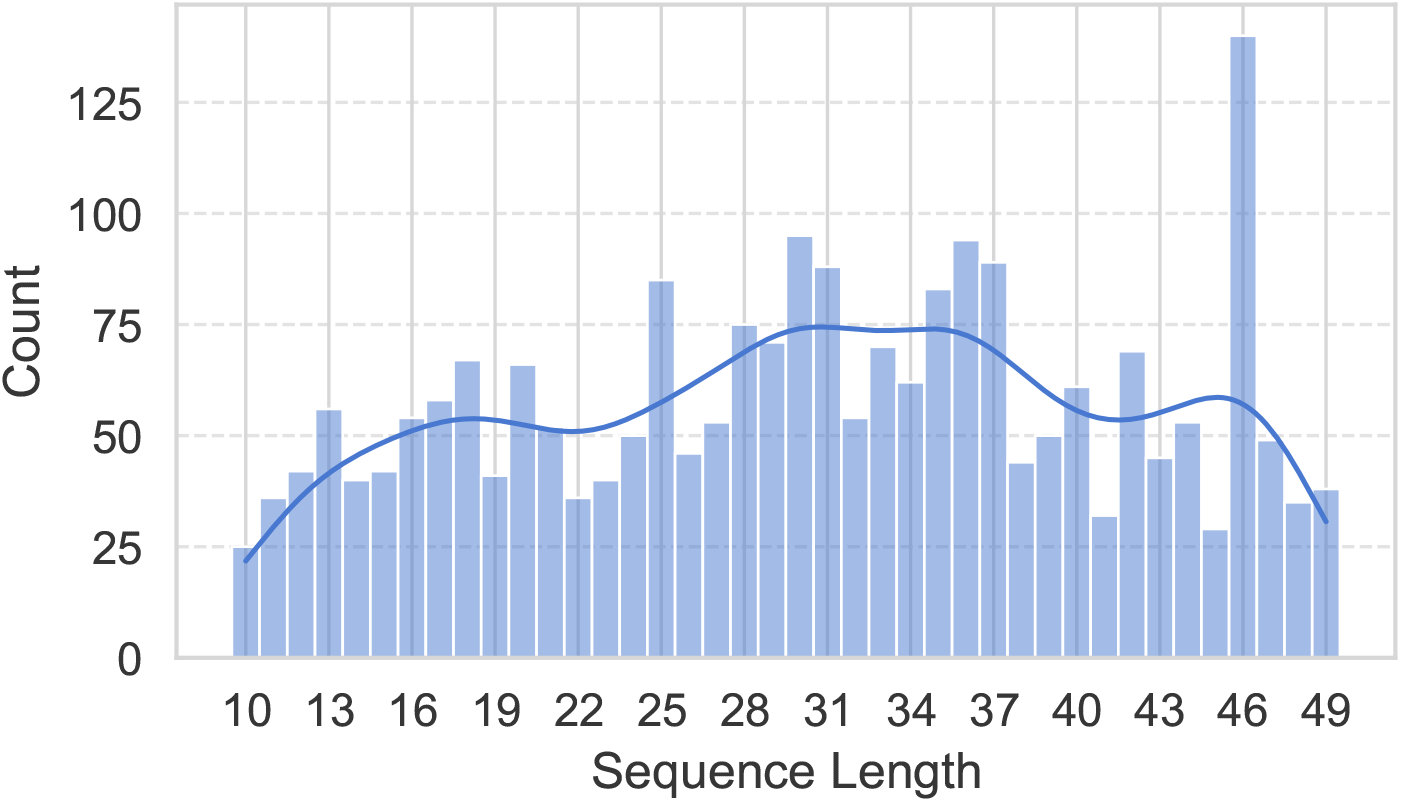
Distribution of Sequence Length

**Fig. 46:**
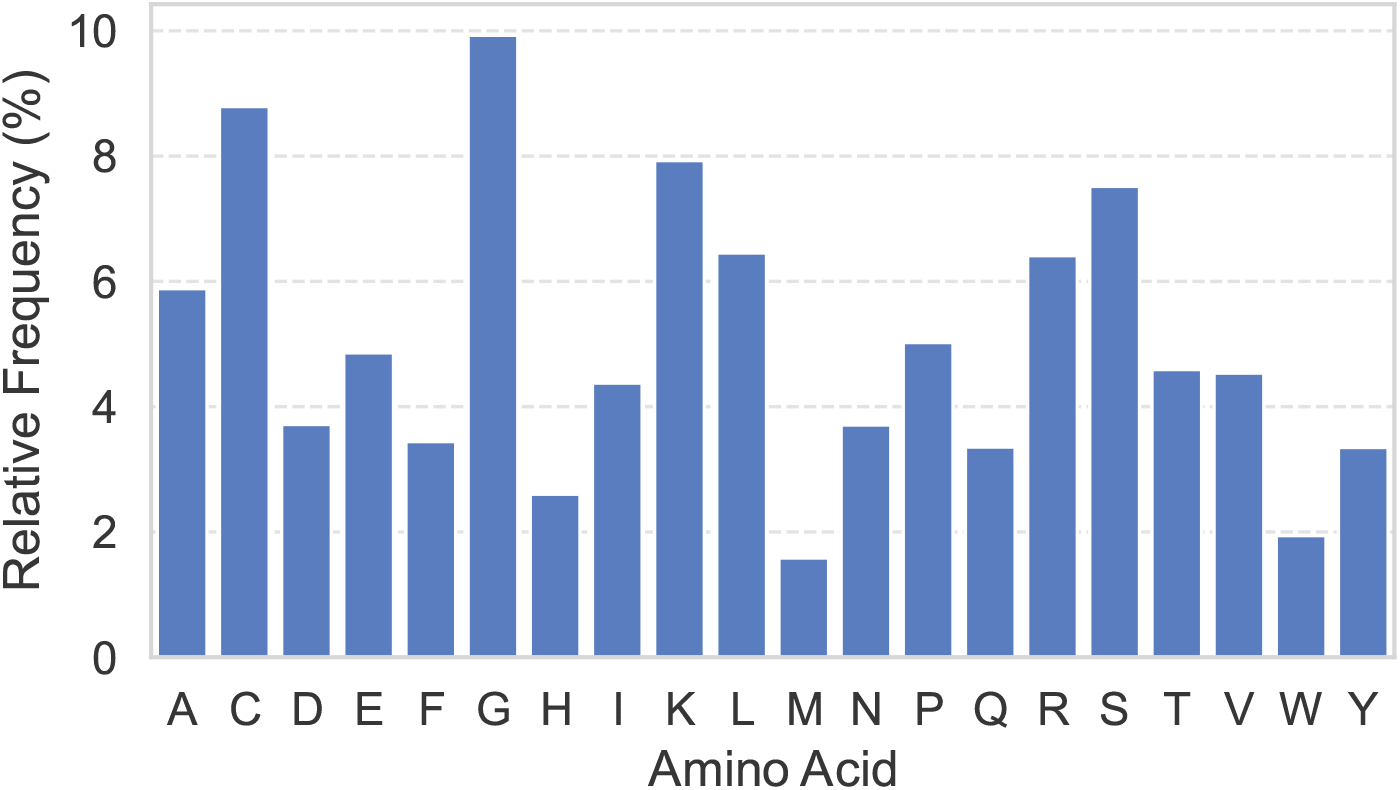
Distribution of Amino Acid Composition

## Appendix I Structural Comparison and Evaluation Metrics

Predicted peptide structures were evaluated by comparing them to their corresponding native PDB structures using several widely adopted structural similarity metrics.

### A. Root Mean Square Deviation (RMSD)

The **Root Mean Square Deviation (RMSD)** was computed between predicted and native structures to quantify the average deviation in atomic coordinates. RMSD provides a global measure of structural similarity after optimal superposition.

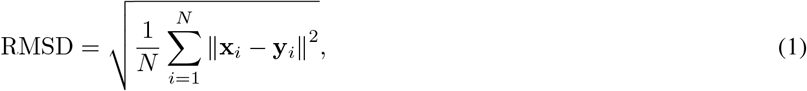

where *N* is the number of aligned C_*α*_ atoms, and **x**_*i*_ and **y**_*i*_ are the Cartesian coordinates of residue *i* in the predicted and reference structures, respectively.

### B. Template Modeling(TM)-score

The **TM-score**[34] measures global structural similarity while accounting for protein length and provides a normalized score ranging between 0 and 1, where higher values indicate greater structural similarity.

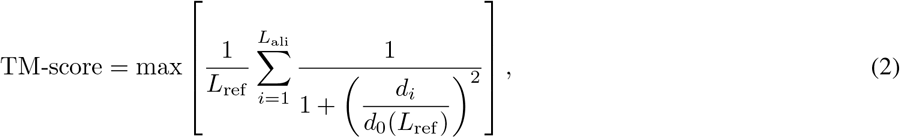

where *L*_ref_ is the length of the reference structure, *L*_ali_ is the number of aligned residues, and *d*_*i*_ is the distance between the *i*-th aligned residue pair after superposition. The length-dependent normalization factor is given by

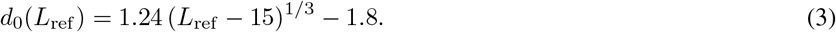

Physically, TM-score can be understood as a topology-sensitive measure; residue pairs that are close after superposition contribute strongly, whereas poorly aligned residue pairs contribute much less. Unlike RMSD, TM-score is less dominated by a small number of large coordinate deviations and is therefore useful for assessing whether the overall fold or topology is preserved.

### C. Global Distance Test (GDT)-Total Score(TS)

The **GDT-TS** metric evaluates structural similarity by calculating the fraction of residues that can be superimposed within predefined distance thresholds.

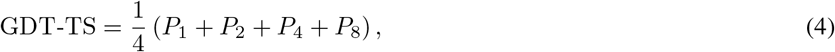

where *P*_*d*_ denotes the percentage of aligned residues whose C_*α*_ distance from the reference structure is less than *d* Å after optimal superposition. Thus, *P*_1_, *P*_2_, *P*_4_, and *P*_8_ correspond to thresholds of 1, 2, 4, and 8 Å, respectively, following the LGA/GDT framework [35]. GDT-TS asks what fraction of the peptide can be placed close to the reference structure at increasingly relaxed distance cutoffs. A high GDT-TS indicates that a large portion of the structure can be aligned within these thresholds, even if some residues deviate substantially.

### D. Native Contact Analysis

As a physics-based descriptor of structural nativeness, the fraction of **native contacts**, *Q*, was also calculated. The **Q**, was calculated using a soft-contact switching function following Best et al. [37]. In the present implementation, all non-hydrogen atom pairs were considered as candidate contacts, excluding atom pairs from residues separated by fewer than two positions in sequence. The candidate contact set was defined as

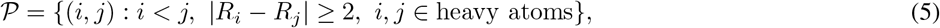

where *R*_*i*_ and *R*_*j*_ are the residue indices of atoms *i* and *j*, respectively. Native contacts were identified from the reference structure as atom pairs whose reference distance 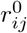 was below the cutoff distance *r*_cut_:

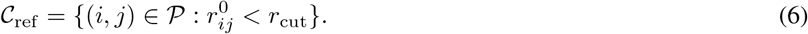

In this work, the cutoff distance was set to

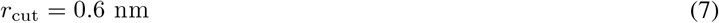

For each reference contact (*i, j*) ∈ *C*_ref_, the contact contribution in the model structure was evaluated using the switching function

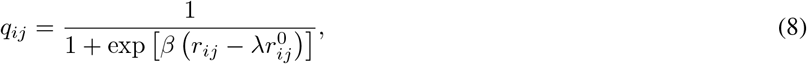

where *r*_*ij*_ is the atom-pair distance in the model structure, 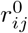 is the reference distance, *λ* is the distance-tolerance factor, and *β* controls the steepness of the switching function. The parameter values used in this work were

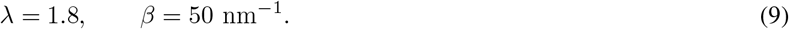

The total fraction of native contacts was calculated by averaging over all reference contacts:

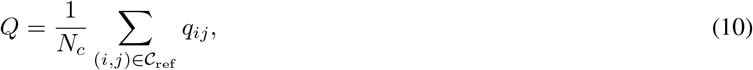

where *N*_*c*_ is the total number of reference contacts,

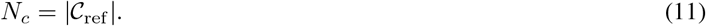

The **Q** ranges from 0 to 1, with larger values indicating greater preservation of the reference contact network.

### E. Local Distance Difference Test (LDDT)

Local structural accuracy was assessed using the **Local Distance Difference Test (LDDT)**[40], which evaluates the preservation of local atomic distances without requiring global structural superposition.

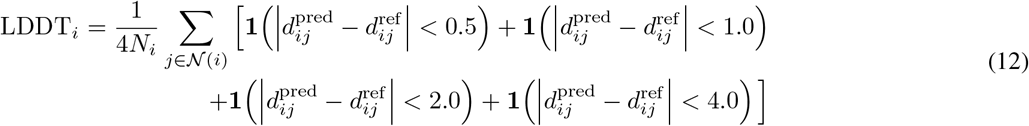

where *N* (*i*) is the set of neighboring residues or atoms of residue *i* within a predefined cutoff in the reference structure, *N*_*i*_ = |*N* (*i*)|, and **1**(*·*) is the indicator function. The global LDDT score is then obtained by averaging over all residues:

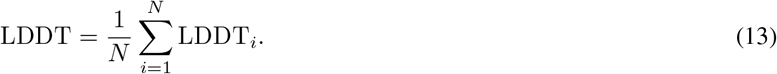

Several prediction models provide internal confidence estimates for predicted structures. In particular, AlphaFold-based models produce the **predicted Local Distance Difference Test (pLDDT)** score as a residue-level confidence measure. To evaluate the reliability of model confidence estimates, we analyzed the relationship between pLDDT scores and experimentally derived LDDT values. This comparison provides insight into how well model-predicted confidence correlates with actual structural accuracy.

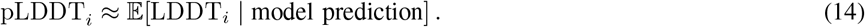

To assess confidence calibration, the Pearson correlation coefficient between experimentally derived LDDT and model-predicted pLDDT was computed as

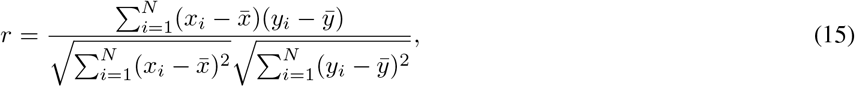

where *x*_*i*_ and *y*_*i*_ denote the LDDT and pLDDT values, respectively, and 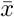 and 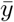 are their corresponding sample means.

Prediction accuracy across models and peptide categories was analyzed using a combination of descriptive statistics and distribution-based visualizations. For structural similarity metrics, including RMSD, TM-score, GDT-TS, and native contact recovery, violin plots were used to illustrate the distribution of values across different prediction models and dataset buckets. To assess the relationship between model confidence estimates and structural accuracy, scatter plots of lDDT versus Plddt were generated. A regression line was fitted to quantify the correlation between predicted confidence scores and experimentally derived structural similarity metrics.

All statistical analyses were performed separately across the two dataset stratification schemes: (1) Length-based buckets, (2) Secondary structure-based buckets. Within each bucket, prediction performance was compared: (1) Across different models, to evaluate relative model performance, (2) Across peptide categories, to assess how structural characteristics influence prediction accuracy. This bucketed analysis enables a detailed understanding of how model performance varies across peptides of different lengths and structural compositions.

## Notes

https://doi.org/10.5281/zenodo.21073120

### Competing Interest Statement

The authors have declared no competing interest.

